# The Divisive-Normalization Model of V1 Neurons: A Comprehensive Comparison of Physiological Data and Model Predictions

**DOI:** 10.1101/081497

**Authors:** Tadamasa Sawada, Alexander A. Petrov

## Abstract

The physiological responses of simple and complex cells in the primary visual cortex (V1) have been studied extensively and modeled at different levels. At the functional level, the divisive normalization model (DNM, Heeger, 1992) has accounted for a wide range of single-cell recordings in terms of a combination of linear filtering, nonlinear rectification, and divisive normalization. We propose standardizing the formulation of the DNM and implementing it in software that takes static grayscale images as inputs and produces firing-rate responses as outputs. We also review a comprehensive suite of 30 empirical phenomena and report a series of simulation experiments that qualitatively replicate dozens of key experiments with a standard parameter set consistent with physiological measurements. This systematic approach identifies novel falsifiable predictions of the DNM. We show how the model simultaneously satisfies the conflicting desiderata of flexibility and falsifiability. Our key idea is that, while adjustable parameters are needed to accommodate the diversity *across* neurons, they must be fixed for a given *individual* neuron. This requirement introduces falsifiable constraints when this single neuron is probed with multiple stimuli. We also present mathematical analyses and simulation experiments that explicate some of these constraints.

## 1 Introduction

The primary visual cortex (V1) is the most studied cortical area. Beginning with the seminal studies of Hubel and Wiesel (1959, 1962), V1 neurons have been studied extensively in physiology for half a century (see Albrecht, Geisler, & Crane, 2003; Andoni, Tan, & Priebe, 2013; Angelucci & Shushruth, 2013; Ferster & Miller, 2000; Fitzpatrick, 2000; Hubel & Wiesel, 1977; Lamme, 2003; Lennie & Movshon, 2005, for reviews). A wide range of models have been proposed to account for various properties of V1 neurons at various levels of analysis (see Albrecht, Geisler, Frazor, & Crane, 2002; Carandini et al., 2005; Carandini, Heeger, & Movshon, 1999; Graham, 1992, 2011; Grossberg, 1988; Hubel & Wiesel, 1977; Sompolinsky & Shapley, 1997, for reviews).

These models of the V1 neurons can be classified into three types: namely, *descriptive, functional*, and *structual* models (Albrecht et al., 2003, p. 759; see also Herz, Gollisch, Machens, & Jaeger, 2006). A *descriptive model* applies statistical regression techniques to summarize and regularize a set of empirical measurements with a mathematical equation. These measurements can be characterized by a small number of parameters of this equation. For example, the tuning bandwidth of a V1 neuron can be estimated by fitting a Gaussian function to its empirical tuning-curve. A *functional model* aims to characterize a variety of response properties within the context provided by a visual information-processing algorithm. For example, a simple cell can be modeled approximately as a linear filter followed by rectification (e.g., Movshon, Thompson, & Tolhurst, 1978c). Ideally, a *functional* model can take the stimulus image as input and calculate the response. This makes it possible to perform simulation experiments with a functional model by presenting it with a set of stimuli and examining the predicted responses. Typically, *functional* models of V1 neurons involve various combinations of linear and nonlinear operations defined via algebraic equations. The goal of a *functional* model is to characterize the neuron’s response to a given stimulus. All models discussed in the present article are of this type. Note that a *functional* model can be regarded as an intermediate step toward a *structural* model (Marr, 1982, ch. 1.2). A *structural model* aims to characterize some aspect of the biophysical and/or biochemical processing mechanisms in the early visual system. Typically, it is formulated either as an algorithm representing a neural circuit or as a system of differential equations (see Ben-Yishai, Bar-Or, & Sompolinsky, 1995; Brosch & Neumann, 2014; Chance & Abbott, 2000; Ellias & Grossberg, 1975; Grossberg, 1988; Izhikevich, 2007; Kough & Poggio, 2008; Schwabe, Obermayer, Angelucci, & Bressloff, 2006; Shapley & Xing, 2013; Somers, Nelson, & Sur, 1995; Somers et al., 1998; Zhaoping, 2011, for reviews and examples). *Structural* models are beyond our scope here.

The present review is based on a functional model of the static (steady-state) properties of simple and complex cells in V1. The temporal dynamics of the neuronal response is beyond our present scope. (See, e.g., Albrecht et al., 2002; Heeger, 1993; Albrecht et al., 2003, for reviews on temporal dynamics. We will revisit this issue in Section 4, General Discussion.) Most experiments discussed below used prolonged (steady-state) stimuli such as drifting and/or flickering gratings with relatively long durations (e.g., a few seconds). Under these conditions, the transient response triggered by the stimulus onset can be bracketed out of the analysis to a good approximation, and the most important dependent variable is the steady-state firing rate of the neuron as measured by a post-stimulus time histogram. The models discussed below are functional models that take a single grayscale image as input and produce a single number for each neuron simulated that we take to be “the response” to a given stimulus.

Within this circumscribed scope, there is “a fairly well agreed on standard model of V1 response properties, usually involving a combination of linear filtering, half-wave rectification and squaring, and response normalization” (Carandini et al., 2005, p. 10590). Whereas it is still unknown how well this *divisive normalization model (DNM)* can account for the full complexity of the V1 population code for time-varying naturalistic stimuli (Olshausen & Field, 2005; but see Rust & Movshon, 2005), it is consistent with much of the available data to a good approximation. The DNM was developed over a number of years during which it combined experimental (e.g., R. L. De Valois, Albrecht, & Thorell, 1982; Hubel & Wiesel, 1959; Movshon et al., 1978c) and theoretical (e.g., Grossberg, 1973) contributions, as well as interdisciplinary explorations of the correspondence between the physiological data and the mathematical formalisms (e.g., Albrecht & Geisler, 1991; Heeger, 1992b; Carandini & Heeger, 1994). Versions of this model have been applied to a broad spectrum of data ranging from single-cell recordings (see, e.g., Albrecht et al., 2003; Heeger, 1992b; Carandini & Heeger, 2012, for reviews), multi-electrode population recordings (e.g., Busse, Wade, & Carandini, 2009; Goris, Wichmann, & Henning, 2009), and psychophysical experiments (e.g., Itti, Koch, & Braun, 2000; Olzak & Thomas, 1999, 2003; To, Lovell, Troscianko, & Tolhurst, 2010; Jesús & Laparra, 2010).

Unfortunately, there is no standard formulation of the DNM. Different authors and publications use different mathematical expressions and idiosyncratic parameterizations. There is a clear family resemblance across these model variants—the verbal description quoted above summarizes the core DNM ideas—but the main results are scattered across dozens of journal articles spanning decades of experimental and theoretical development. A typical article reports neurophysiological data collected with a certain restricted class of stimuli, and then reports the fit of a DNM variant customized for the particular application. This practice makes it difficult to compare the results across studies.

Here, we propose a standard formulation of the DNM and test its validity with respect to a comprehensive suite of empirical phenomena (listed in Table 1 below). This article is organized around a series of figures with side-by-side comparisons between data patterns illustrating various properties of real V1 neurons and the corresponding patterns simulated with the DNM.

**Table 1:** Phenomena accounted for by the divisive normalization model (Equation 13) in our simulation experiments. RF = receptive field, CF = contrast response function.

| N. | Phenomenon | Figure |
| --- | --- | --- |
| 1. | Size tuning: The receptive field (RF) has limited spatial extent | 5 |
| 2. | The measured RF diameter increases as the grating contrast decreases | 6 |
| 3. | The measured RF diameter decreases for non-preferred orientations | 7 |
| 4. | The measured RF diameter decreases for non-preferred spatial frequencies | 8 |
| 5. | The response decreases monotonically as the hole in an annular grating increases | 9 |
| 6. | The response relationship with hole size is nearly invariant to stimulus contrast | 9 |
| 7. | The contrast response function (CF) has a characteristic sigmoidal shape | 3 |
| 8. | Supersaturation effect: The CF can slope downwards at very high contrasts | 10 |
| 9. | The CF is affected by visual noise added to the grating stimulus | 11 |
| 10. | The CF is scaled down for gratings with non-preferred orientations | 3 |
| 11. | The CF is scaled down for gratings with non-preferred spatial frequencies | 3 |
| 12. | The CF is affected by the size of the grating patch | 12 |
| 13. | Orientation tuning: The response is maximal for the preferred orientation | 2 |
| 14. | Spatial-frequency tuning: The response is maximal for the preferred frequency | 2 |
| 15. | Orientation bandwidths become narrower as the size of grating patch increases | 13 |
| 16. | Frequency bandwidths become narrower as the size of grating patch increases | 13 |
| 17. | Orientation bandwidths tend to be invariant with respect to the grating contrast | 14 |
| 18. | Frequency bandwidths tend to become narrower as the contrast decreases | 14 |
| 19. | The spatial-frequency tuning function has a secondary peak for square gratings | 15 |
| 20. | The CF shifts leftward for square gratings compared to sinusoidal ones | 15 |
| 21. | Cross-orientation suppression: orientation tuning of the mask grating | 16 |
| 22. | Cross-orientation suppression: spatial-frequency tuning of the mask grating | 16 |
| 23. | Cross-orientation suppression: the mask contrast affects the CF | 17 |
| 24. | Surround suppression: orientation tuning of the surrounding grating | 18 |
| 25. | Surround suppression: spatial-frequency tuning of the surrounding grating | 18 |
| 26. | Surround suppression: the contrast of an annular grating affects the CF | 19 |
| 27. | Surround suppression: the orientation of an annular grating affects the CF | 20 |
| 28. | Mapping the receptive field of a simple cell with a light spot | 21 |
| 29. | Mapping the RF of a simple cell using the reverse correlation method | 22 |
| 30. | Comparison of the mapped RF of a simple cell and its spatial-frequency tuning | 23 |

In addition to consolidating a large and scattered literature, our systematic approach sets the stage for identifying novel falsifiable predictions of the DNM. An adequate functional model should satisfy the conflicting desiderata of flexibility and falsifiability at the same time (Popper, 1959; Roberts & Pashler, 2000). On the one hand, the model needs to be flexible enough to accommodate the variety of V1 neurons with some adjustable parameters. The data reviewed below were recorded from neurons from different species (e.g., cats, new/old world monkeys, rabbits, rodents, and ferrets) under different conditions (e.g., anesthesia vs. alertness) and different experimental protocols. Furthermore, there is substantial variability within a sample of neurons recorded from a single animal under constant conditions. Clearly, adjustable parameters are needed to accommodate this diversity. Note also that it is easy to make the model more flexible by adding more parameters. On the other hand, if the model becames so flexible as to be able to fit any response pattern, it would become devoid of all empirical content (Roberts & Pashler, 2000). Whereas it might still be useful as a descriptive formalism for a succinct characterization of properties such as tuning bandwidths, such a model would not constrain our theories about the functional organization of the visual system. To have empirical content, the model must be restrictive enough to rule out at least some possible data patterns.

While the model’s parameters may have to be adjusted flexibly to accommodate the diversity across neurons, they must be fixed for a given *individual* neuron. This introduces falsifiable constraints when this single neuron is probed with a judiciously-chosen suite of stimuli. This article presents mathematical analyses and simulation experiments that explicate some of these constraints. These novel theoretical results emerge naturally from our side-by-side examination of dozens of phenomena within the framework of a single model. This type of analysis focuses on the *qualitative* patterns that can be produced by a model under a given parameterization (Pitt, Kim, Navarro, & Myung, 2006). It contrasts with the typical approach in the experimental literature where, with some notable exceptions (e.g., Tadmor & Tolhurst, 1989), one or more models were compared in terms of their *quantitative* fits to physiological data pertaining to a single phenomenon.

Our review is of potential interest to several groups of readers. First, readers interested in the neurophysiology of simple and complex cells in the primary visual cortex will find a systematic series of figures with representative data from classic experiments, as well as their interpretation under the DNM. Single-cell data published over a 50-year span were digitized from select figures in the original reports and are replotted below. Second, readers interested in functional modeling of the early visual system will find systematic exposition and motivation of the DNM, as well as its empirical grounding. Third, expert modelers of the early visual system will find mathematical derivations and simulation experiments that identify novel falsifiable predictions of the DNM. Last but not least, modelers who need an off-the-shelf front-end to a larger model (e.g, Petrov, Dosher, & Lu, 2005, 2006; Jacobs, 2009) will find a general-purpose parameterization of the DNM and a standard parameter set (Table 2 below) that is consistent with almost all phenomena listed in Table 1. The model was implemented as a software program for Matlab (The MathWorks, 2015). This software takes a static grayscale image as input and produces a matrix of firing-rate responses for a population of DNM neurons centered on a single retinal location and tuned for a range of orientations and spatial frequencies.

The rest of the article is organized as follows: Section 2 presents the DNM, its proposed parameterization, and computational implementation. Section 3 reviews over two dozen empirical phenomena and interprets them through the lens of the DNM. It also reports mathematical analyses, simulation results, and some novel predictions. Finally, there is a general discussion (Section 4) followed by mathematical appendices.

**Table 2:**
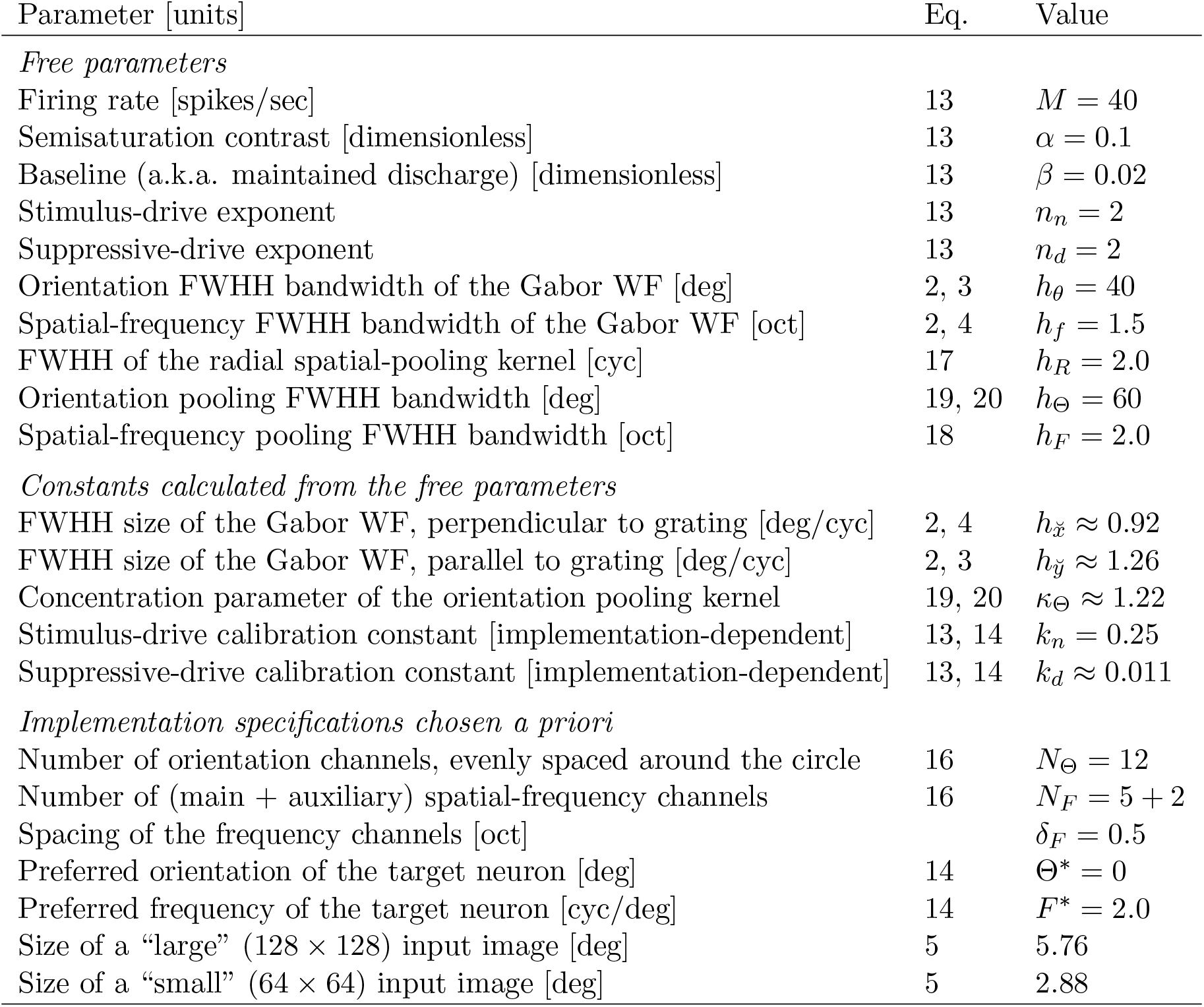
Parameters of the divisive normalization model (Equation 13). The values for the ten free parameters are compatible with typical neurophysiological measurements of representative simple and complex cells. Moreover, the model under this *standard parameterization* accounts qualitatively for the phenomena in Table 1. All simulation results in this article were produced with these parameter values unless explicitly stated otherwise. FWHH = full width at half height, WF = weighting function, cyc = wave cycles, oct = octaves = log[cyc/deg], Eq = Equation.

## 2 Models

The essential components of the divisive normalization model (DNM) are the linear filters, the static nonlinearities, and divisive normalization. These components will be described below. But, before this is done, we must acknowledge an important pre-processing step,namely, *light adaptation* (or *luminance gain control*). Adaptation is primarily accomplished in the retina (Shapley & Enroth-Cugell, 1984). It matches the limited dynamic range of the neurons to the locally-prevalent luminance. The DNM does not model this light adaptation explicitly. It simply assumes it has been incorporated into the encoding of the input images. This assumption is justified in situations when the stimuli are embedded in a large uniform gray background, and when the visual system has adapted to the baseline luminance level *L*_*b*_. The input to the model is a matrix *I*(*x,y*) of local contrast around this fixed baseline:

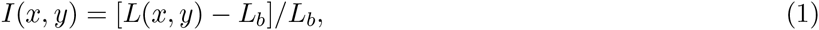

where *L*(*x, y*) is the luminance at coordinates *x* and *y*. In this notation, a sinusoidal grating with contrast *c* modulates between *I*_*min*_ = -*c* and *I*_*max*_ = +*c*, and has zero mean. The maximal possible contrast (*c*_*max*_ = 1) of a grating is attained when the lowest intensity is zero and the highest intensity is twice the mean.

### 2.1 Linear model

The majority of V1 neurons respond selectively to a variety of stimulus features including position, size, orientation, and spatial frequency (e.g., R. L. De Valois, Albrecht, & Thorell, 1982; Hubel & Wiesel, 1959, 1968; Pollen & Ronner, 1982; Watkins & Berkley, 1974). A typical V1 neuron responds to stimulation within a circumscribed region called the *(classical) receptive field (RF)*. Different neurons have RFs centered on different positions, and V1, as a whole, forms a *topographic map* (e.g., Schwartz, Tootell, Silverman, Switkes, & De Valois, 1985). This population code is beyond our scope. We are modeling a representative individual neuron. Note that, in this article, the origin of the xy coordinate system is placed conventionally at the center of the neuron’s receptive field.

Hubel and Wiesel (1959, 1962) introduced the influential distinction between *simple* and complex cells in V1. According to their original definition (Hubel & Wiesel, 1962), simple cells have four characteristic properties: (1) distinct excitatory and inhibitory sub-regions within the RF, (2) spatial summation within a given subregion, (3) mutual antagonism between sub-regions, and (4) the responses to novel stimuli can be predicted to a good approximation on the basis of the spatial arrangement of the sub-regions. These four properties would be expected from a linear spatio-temporal filter and they would motivate the application of Linear-Systems Theory (e.g., Lathi, 2005) to the study of spatial vision (e.g., R. L. De Valois & De Valois, 1988; Graham, 1989; Maffei & Fiorentini, 1973; Shapley & Lennie, 1985). Quantitative tests have identified a subpopulation of V1 neurons that exhibit these linear properties to a good approximation (e.g., Andrews & Pollen, 1979; Movshon et al., 1978c; Pollen & Ronner, 1982; see Albrecht et al., 2003, for review). This Linear-Systems approach is reinforced by a rich body of psychophysical data (e.g., Campbell & Robson, 1968; Cornsweet, 1970) and theory that supports the existence of *channels* selective for orientation and spatial frequency (R. L. De Valois & De Valois, 1988; Graham, 1989). But, two caveats should be kept in mind here (Albrecht et al., 2003). First, the match between the measured single-cell responses and the linear-systems predictions is always approximate, never exact, because they are systematic nonlinearities (discussed below). Second, the simple- vs. complex-cell distinction probably denotes the endpoints of a continuum rather than a sharp dichotomy (Mechler & Ringach, 2002). These caveats notwithstanding, this distinction has proven its theoretical utility and is widely used in V1 models.

The spatial layout of simple-cell receptive fields has been mapped out using local stimulus probes (e.g., Hubel & Wiesel, 1959, 1962, 1968; Movshon, Thompson, & Tolhurst, 1978b, see Section 3.3.6 for further references). The RF of a typical simple cell consists of alternating excitatory and inhibitory sub-regions (Hubel & Wiesel, 1962; Maffei & Fiorentini, 1973; R. L. De Valois, Albrecht, & Thorell, 1982; Sengpiel, Sen, & Blakemore, 1997). This 2D spatial pattern can be approximated well by the *Gabor function* in Equation 2 (Daugman, 1980, 1985; Field & Tolhurst, 1986; J. P. Jones & Palmer, 1987a, 1987b; Kulikowski, Marčelja, & Bishop, 1982; Marčelja, 1980; Ringach, 2002). Mathematically, a Gabor function 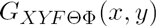 is a product of a Gaussian envelope and a sinusoidal grating:

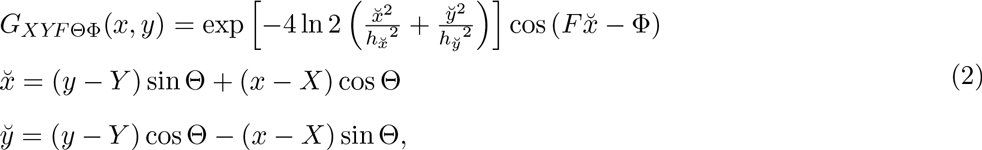

where *x* and *y* (degrees of visual angle, deg) are positions across the image, and *X* and *Y* define the center of the RF. The grating has spatial^1^ frequency *F* (cycles/deg, cpd), phase Φ, and orientation Θ. The parameters 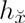 and 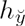 control the full width at half height (FWHH) of the Gaussian envelope along the orthogonal and parallel directions as shown in Figure 1. See Appendix A for more details.

**Figure 1:**
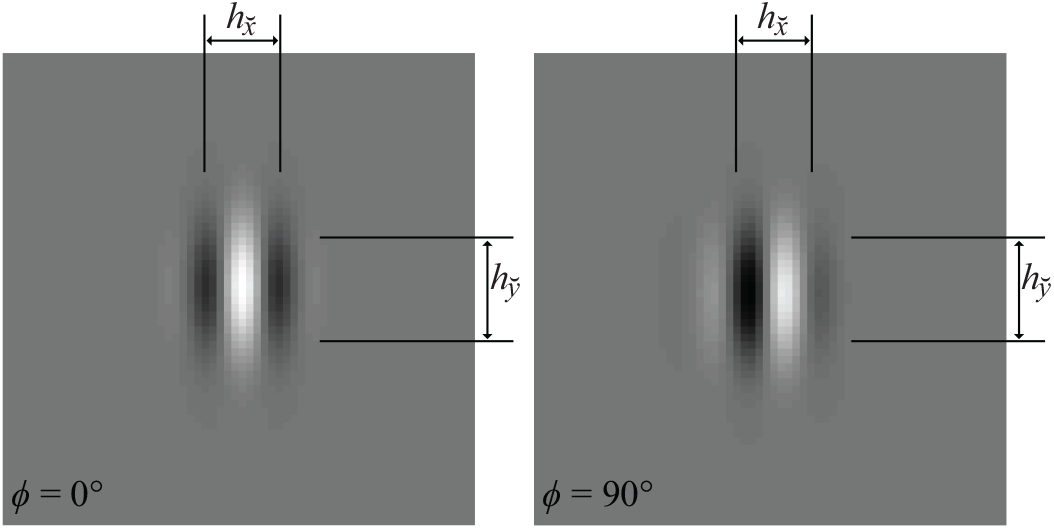
Examples of Gabor patches with phase Φ = 0° and Φ = 90°, vertical orientation (Θ = 0°), and spatial frequency *F =* 2 cpd (Equation 2). The parameters 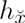 and 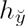 control the full width at half height (FWHH) of the Gaussian envelope along the directions that are perpendicular and parallel to the grating, respectively.

The alternating arrangement of excitatory and inhibitory RF regions makes the simple cell selectively responsive (or *tuned)* to the orientation and frequency of the stimuli. It is very informative to probe the neuron with a battery of sinusoidal gratings covering a range of orientations and frequencies. Such probing in the *frequency (Fourier) domain* complements the local probing in the space domain for simple cells, and is necessary when studying complex cells because they do not respond to local probes. There is abundant evidence that most excitatory neurons in V1 (as well as many other visual cortical areas) are tuned for orientation and spatial frequency (e.g., Albrecht, De Valois, & Thorell, 1980; Watkins & Berkley, 1974; Pollen & Ronner, 1982, 1983, see Section 3.3.3 for further references). Figure 2 illustrates typical empirical^2^ *tuning curves* with respect to orientation and spatial frequency. For example, the tuning curve in Figure 2A has a peak at 0° and FWHH of approximately 45°, which measure this neuron’s *preferred orientation* and the *orientation bandwidth,* respectively.

**Figure 2:**
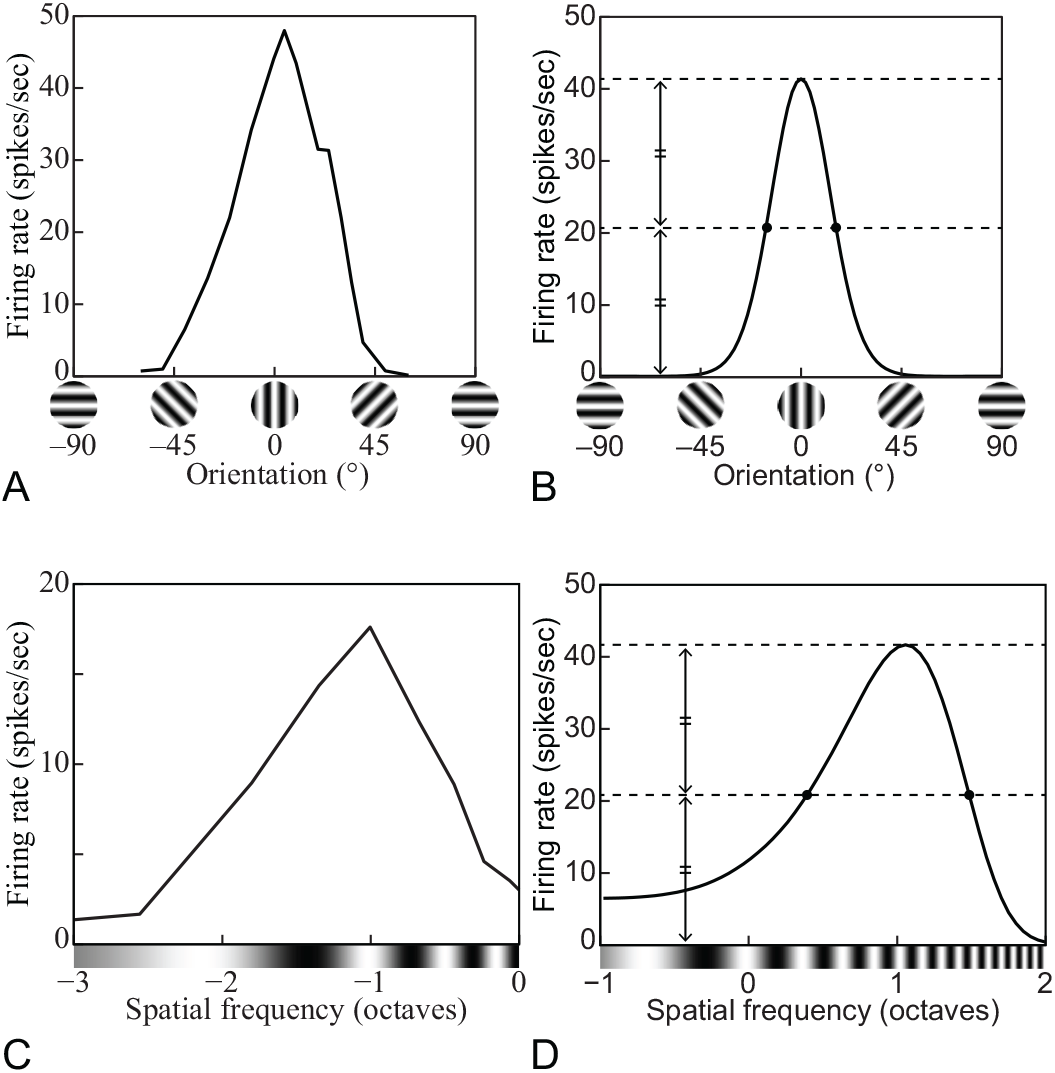
(A) A representative orientation tuning curve—in this case from a V1 complex cell of an anesthetized cat. Replotted from Rose and Blakemore (1974, Figure 1). (B) Orientation tuning curve of the DNM neuron with default parameters, probed with gratings with 100% contrast, 2.88° size, and neuron’s preferred frequency. (C) A representative spatial-frequency tuning curve—in this case from a V1 simple cell of an anesthetized cat. Replotted from Li and Li (1994, Figure 7C). (D) Spatial-frequency tuning curve of the DNM neuron with default parameters, probed with gratings with 100% contrast, 2.88° size, and neuron’s preferred orientation. Dotted lines depict the full- and half-heights of the curves, and the distances between the dots depict the full-widths at half-height. (DNM = divisive normalization model, defined later in the text. See Phenomena 13 and 14 in Table 1 below.)

With a perfectly linear filter (Lathi, 2005), the response profile in the space domain completely determines the tuning in the frequency domain (via the *Fourier transform*) and vice versa (via the *inverse Fourier transform*). Specifically, consider a linear filter whose *weighting function* is a Gabor patch in the space domain (Equation 2). The tuning function of this filter is a bivariate Gaussian in the frequency domain (Graham, 1989, p. 85). Moreover, for a given frequency *F*, the orientation bandwidth 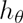 (in degrees) is inversely proportional to the size 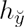 of the patch along the direction parallel to the grating:

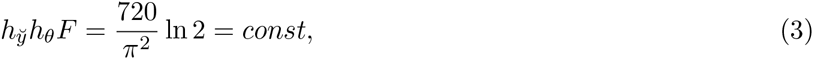

where the constant 720 ln(2)/π^2^ comes from the conversion from degrees to radians and from FWHH to standard deviation. See Appendix A for details. The reason for this inverse relationship is intuitively clear from Figure 1: To estimate the stimulus orientation with high precision (low *h*_*θ*_), it is necessary to have a large baseline for measurement along the length of the bars. An analogous inverse relationship exists between the frequency bandwidth *h*_*f*_ and the perpendicular size 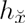. Intuitively, to estimate the stimulus frequency with high precision (low *h*_*f*_), it is necessary to be able to “count” many cycles within the width of the filter. It is more convenient to express the frequency bandwidth *h*_*f*_ in *octaves* on a log_2_ scale instead of linear units. The exact relationship (derived in Appendix A) is

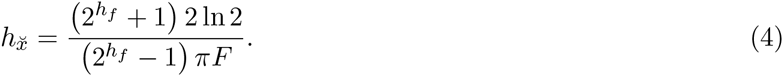

Real neurons are never perfectly linear but the tuning properties of simple and complex cells are in qualitative agreement with the predictions of the linear model. “The two-dimensional tuning curves are mostly moderately elongated along a radial axis [in the Fourier plane], and extreme or amorphous shapes (e.g., sausages, amoebas) are rare” (Lennie & Movshon, 2005, p. 2020). Typically, the preferred orientation does not depend much on the frequency of the test grating, and vice versa (Webster & De Valois, 1985). This means that the joint orientation-by-frequency tuning curve can be modeled as the product of two orthogonal dimension-specific curves (cf. Fig. 2), which is in agreement with the linear prediction.

The output *E*_*S*_ of a linear filter to a stimulus image *I* is a scalar quantity equal to the dot product^3^ of the image with the weighting function *G*_*XYFΘΦ*_ of the filter:

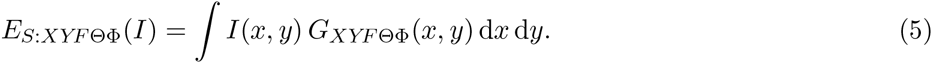

The *weighting function* (*WF*) determines the properties of the filter. All models discussed in this article use Gabor WFs (Eq. 2). The center *XY* of the Gabor patch determines the center of the receptive field (RF) of the filter (in image space), the frequency *F* of the patch determines the preferred frequency of the filter (in Fourier space), and analogously for the orientation Θ and phase Φ. Of all images with a given energy (i.e., fixed variance of the intensity distribution), the output is maximized by the stimulus that exactly matches the WF (Cauchy-Schwarz inequality, n.d.). In this sense, the *preferred stimulus* of the filter in Equation 5 is the Gabor patch in Equation 2. The absolute value of the output can be interpreted as the similarity between the stimulus and the preferred template, and the sign indicates whether the two are *in phase* or *out of phase.* (An image is in phase with a WF when the bright spots on the image line up with the excitatory regions of the WF and the dark spots line up with the inhibitory regions; it is out of phase if the alignment is the other way around.)

The linear model of a simple cell consists of a linear filtering stage (Eq. 5) followed by *halfway rectification* (Eq. 6). The linear stage is motivated by the extracellular recordings surveyed above, as well as by intracellular recordings (e.g., Jagadeesh, Wheat, & Ferster, 1998) suggesting that the fluctuations in membrane potential of simple cells around the resting potential can be modeled quantitatively in terms of linear summation of synaptic potentials. For our purposes, the output of the linear stage is referred to as the *stimulus drive E*_*S*_ to the simple cell. The stimulus drive can be positive or negative, but the firing rate of a real neuron is always non-negative. This is modeled by a rectifying nonlinearity

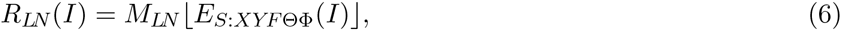

where *R*_*LN*_ is the response of the cell (in spikes/sec, sps) and *M*_*LN*_ is a parameter that, under certain calibration assumptions, defines the maximum firing rate to a preferred grating. The *halfway rectification* operator 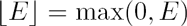 passes positive values unchanged and converts negative values to zero.

The weighting function of a linear neuron coincides with its receptive field. Consequently, the terms *WF* and *RF* are sometimes used interchangeably in the literature. We keep them distinct because the corresponding referents are distinct for nonlinear models. *WF* is a theoretical term that is defined only with respect to a model with a linear filtering stage (Eq. 5). By contrast, the RF is defined operationally for any (real or simulated) neuron as the region of the visual field where stimulus presentation induces changes in firing rate. The RF is smaller than the WF in models involving nonlinearity and/or suppression as discussed below.

Complex cells differ from simple cells because of the absence of distinct excitatory and inhibitory subregions in their RFs (Hubel & Wiesel, 1962; Watkins & Berkley, 1974). Complex cells are relatively invariant to the phase of the stimuli (Maffei & Fiorentini, 1973; Movshon et al., 1978c; R. L. De Valois, Albrecht, & Thorell, 1982; Sengpiel et al., 1997; Ibbotson, Price, & Crowder, 2005). For example, they respond indiscriminately to light and dark bars, as long as the bar stands out from the gray background. They are sensitive to the stimulus orientation and spatial frequency, however, and their tuning curves are very similar to those of simple cells.

Complex cells are usually modeled in terms of several linear filters whose outputs are nonlinearly transformed and then combined (see Bair, 2005; Martinez & Alonso, 2003; Mechler & Ringach, 2002; Spitzer & Hochstein, 1988, for reviews). “A key feature of these models is that the underlying linear filters—not the later nonlinearities—determine the set of stimuli to which the neuron will respond” (Lennie & Movshon, 2005, p. 2023). The most influential model of this class is the *energy model* (Adelson & Bergen, 1985; Pollen & Ronner, 1983; Spitzer & Hochstein, 1985; Watson & Ahumada, 1985):

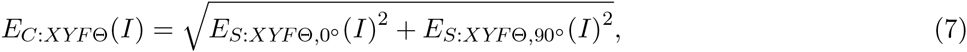

where 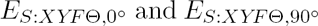 are linear filters (Eq. 5) whose WFs have identical parameters *XYF*Θ but orthogonal phases. Equation 7 produces phase-invariant output *E*_*C*_ via the trigonometric identity *sin*^2^Φ + *cos*^2^Φ = 1. The firing rate *R*_*LN*_(*I*) of the complex cell can be modeled by substituting *E*_*C*_ for *E*_*S*_ in Equation 6 (see, e.g., Emerson, Korenberg, & Citron, 1992; Heeger, 1992b; Lehky, Sejnowski, & Desimone, 2005; Szulborski & Palmer, 1990, for similar formulations). It is tempting to interpret Equation 7 as a formalization of the hierarchical feedforward simple-to-complex arrangement proposed by Hubel and Wiesel (1962). The physiological evidence, however, is more consistent with a non-hierarchical interpretation in terms of a continuum from strongly phase-sensitive (simple) cells to nearly phase-invariant (complex) cells (Bair, 2005; R. L. De Valois & De Valois, 1988; Martinez & Alonso, 2003; Mechler & Ringach, 2002). For our present purposes, it suffices to treat Equation 7 as a mathematically convenient functional description of the *stimulus drive E*_*C*_ to complex cells.

As a notational convenience, it is often useful to bundle the long subscripts into a *tuning preference vector*

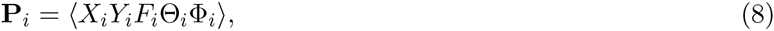

where the phase Φ_*i*_ can take a special non-numerical value to indicate a phase-invariant (complex) cell. In this notation, an image *I* induces stimulus drive *E*_P*i*_(*I*) to a neuron with index *i* and preferences P_*i*_:

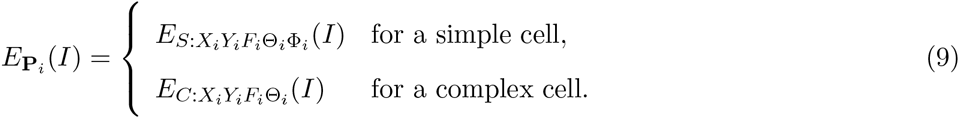

Note that *E*_P*i*_ is a linear operator for the simple cell. That is, for a fixed *i*, 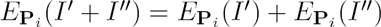 for any pair of images *I′* and *I″*, and 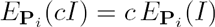 for any number *c* ≥ 0. The latter equality also holds for the complex cell.

The linear model 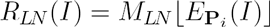 provides a quantitative account of the selectivity of V1 neurons’ responses to bars, edges, and gratings. Moreover, it provides “a credible account of the responses to a variety of more complicated targets, including checkerboards (K. K. De Valois, De Valois, & Yund, 1979), random dot textures and Glass patterns (Smith, Bair, & Movshon, 2004)” (Rust & Movshon, 2005, p. 1647). It dominated the field until the mid-1980s and set the stage for subsequent research that uncovered systematic departures from linearity, to which we now turn.

### 2.2 Hyperbolic-ratio model

The *contrast response function* (CF^4^) describes how a neuron’s response depends on the contrast of the stimulus. Consider a family of stimuli 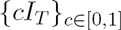 based on the same template image *I*_*T*_ but varying in contrast *c*. We assume throughout this section that the template *I*_*T*_ is normalized so that its contrast is 1.0 (cf. Eq. 1). The CFs of neurons have been measured for various templates, cortical areas, species, and conditions (e.g., Albrecht & Geisler, 1991; Albrecht et al., 2002; Albrecht & Hamilton, 1982; Carandini, Heeger, & Movshon, 1997; Dean, 1981; Derrington & Lennie, 1984; Geisler & Albrecht, 1997; C.-Y. Li & Creutzfeldt, 1984; Sclar, Maunsell, & Lennie, 1990; see Albrecht et al., 2003; Carandini et al., 1999; Graham, 2011; Heeger, 1992a; Lennie & Movshon, 2005, for reviews). Further more, Albrecht, Geisler, Frazor, and Crane (2002) pointed out that “Although there is a great deal of heterogeneity from cell to cell, it is possible to provide a description of the basic properties of the contrast response function that applies to the overwhelming majority of neurons: As the contrast increases from zero, the response increases in an accelerating fashion, remains dynamic over some limited range of contrasts, and then saturates” (p. 888). Figure 3 illustrates the characteristic sigmoidal shape characteristic of a typical CF.

**Figure 3:**
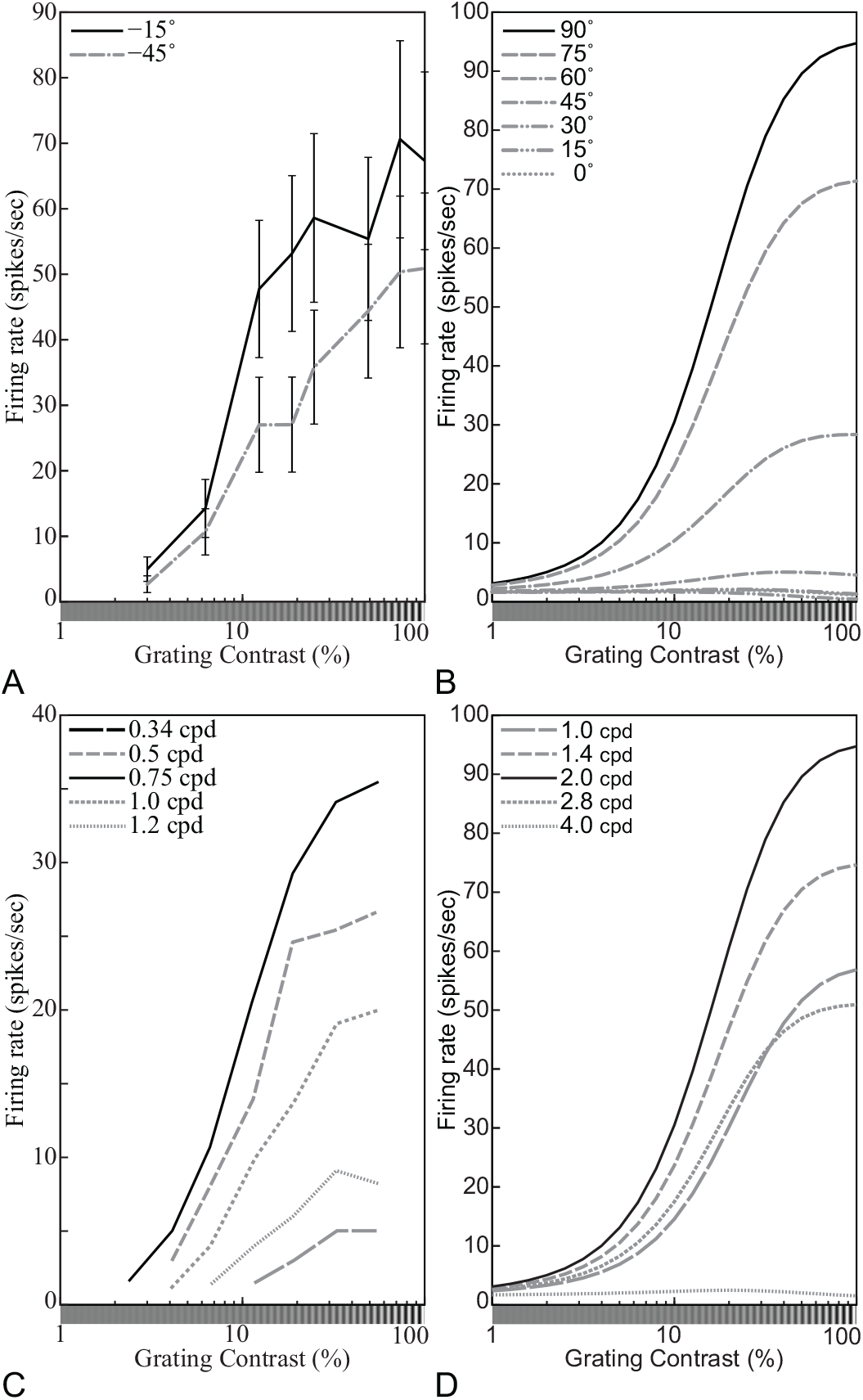
Representative contrast response functions (CFs). (A) Responses of a simple cell to drifting sinusoidal gratings spanning a range of contrasts at two orientations (see legend). Replotted from Carandini, Heeger, and Movshon (1997, Figure 4B, anesthetized macaque, error bars = ±SE). (B) CFs of the DNM neuron with default parameters, probed with gratings with the neuron’s preferred frequency (1.0 octave) and orientations shown in the legend. The size of the stimuli was equal to the measured RF diameter (0.81°). (C) Responses of a V1 neuron to drifting sinusoidal gratings with the neuron’s preferred orientation and spatial frequencies shown in the legend. Replotted from Albrecht and Hamilton (1982, Figure 7A). (D) CFs of the DNM neuron with default parameters, probed with gratings with the neuron’s preferred orientation (0°) and spatial frequencies shown in the legend. The size of the stimuli was equal to the measured RF diameter (0.81°). (DNM = divisive normalization model, defined later in the text. See Phenomena 7, 10, and 11 in Table 1 below.)

The *hyperbolic-ratio model* has been widely used as a descriptive model to fit these data (e.g., Albrecht & Hamilton, 1982, see also^5^ Naka & Rushton, 1966; Legge & Foley, 1980; Graham, 2011). The response *R*_*HB*_ of this model to a stimulus *cI*_*T*_ with contrast *c* and template *I*_*T*_ is

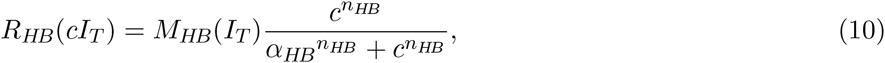

where the *semi-saturation contrast* parameter *α*_*HB*_ expresses the contrast of the image that produces one-half of the saturation level *M*_*HB*_(*I*_*T*_). Note that the latter depends on the template *I*_*T*_. When contrast is plotted on a log axis, the exponent *n*_*HB*_ controls the slope of the CF and *α*_*HB*_ controls its location (Figure 4, see Appendix B for properties of the CF plotted on a linear contrast axis). Note also that the response to a stimulus with zero contrast (i.e., a uniform gray field) is assumed to be zero.

**Figure 4:**
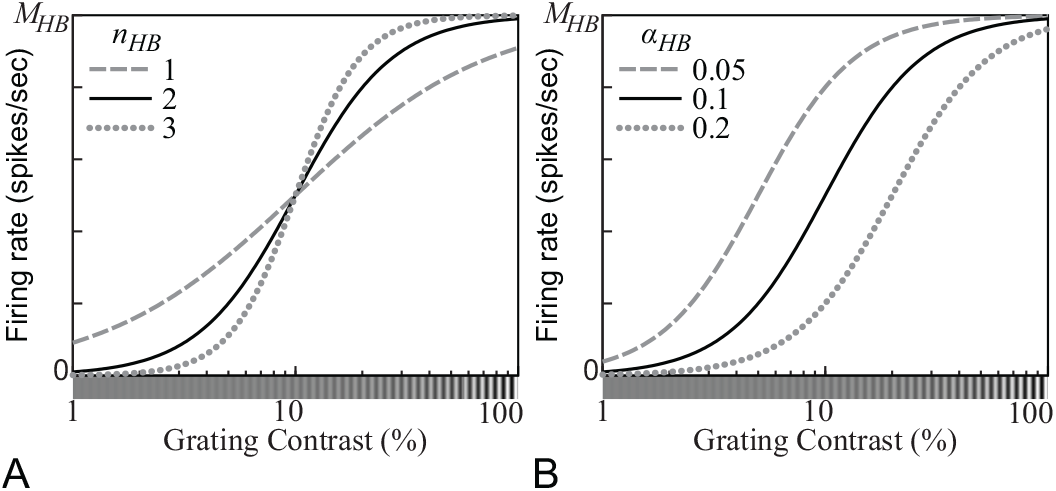
Contrast response functions (CFs) produced by the hyperbolic-ratio model (Equation 10). (A) The exponent parameter *n*_*HB*_ controls the slope of the CF as a function of the log-contrast of the stimulus. (B) The *semisaturation contrast* parameter *α*_*HB*_ controls the location of the CF. A stimulus with contrast *α*_*HB*_ elicits one-half of the saturation level *M*_*HB*_. (*α*_*HB*_ = 0.1 for panel A; *n*_*HB*_ = 2 for panel B.)

The hyperbolic-ratio model (Eq. 10) involves two nonlinear operations: exponentiation and division. The former accounts for the accelerating shape of the CF at low contrasts—that is, for the fact that the CF slope gets steeper and steeper as the contrast increases from zero. Using the logarithmic scale of the contrast *c*, the maximal slope is *n*_*HB*_/4 at *c* = *α*_*HB*_. The divisive operation of the hyperbolic-ratio model saturates the CF at high contrasts. The CF of the hyperbolic-ratio model with these properties can represent shapes of CFs of real neurons well.

Equation 10 predicts that all CFs measured for the same neuron are multiplicatively scaled replicas of each other across the entire contrast range. That is, the ratio of the neuron’s responses to images with identical contrast but different templates *I*_*T′*_ and *I*_*T″*_ is invariant with respect to *c*:

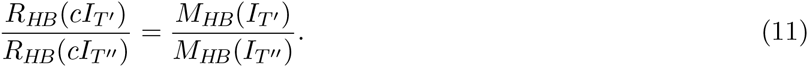

The empirical CFs (Fig. 3) of typical simple and complex cells in V1 are consistent with this prediction in many cases (e.g., Albrecht & Hamilton, 1982; Carandini et al., 1997; see Section 3.3.2 for further references). In particular, when *I*_*T′*_ and *I*_*T″*_ are gratings with different orientations, Equation 11 accounts for the approximate contrast invariance of the orientation tuning curves of typical V1 neurons (e.g., Sclar & Freeman, 1982; Skottun, Bradley, Sclar, Ohzawa, & Freeman, 1987; see Section 3.3.3 for further references). Note, however, that the spatial-frequency tuning curves of many V1 neurons have a slight but systematic dependence on contrast (e.g., Albrecht & Hamilton, 1982; Skottun et al., 1987; see Section 3.3.3 for further references and discussion).

Cortical neurons have a limited dynamic range and their firing rates saturate at high contrasts (e.g., Albrecht & Hamilton, 1982). It is important to note that this saturation is not simply an output nonlinearity because it occurs at a fixed contrast rather than at a fixed response level for different stimuli. Consider the CFs plotted on Figure 3C, for example. The responses to gratings whose frequency is 0.34 cpd saturate below 10 sps even though the same neuron can sustain firing rates above 30 sps when stimulated at its preferred frequency (0.75 cpd). Each neuron is characterized by an entire *family* of CFs—one for each stimulus template. This is why the saturation level *M*_*HB*_ depends on *I*_*T*_ in Equation 10. However, *M*_*HB*_ only acts as a scaling factor and all CFs within the family have the same shape. This shape only depends on *c*. It is for this reason that this type of divisive normalization is called a *contrast-set gain control* (or simply a *contrast gain control*).

The contrast gain control is a cortical mechanism and it should not be confused with the luminance gain control (Eq. 1) found in the retina. Because of the luminance gain control, the maximal contrast is *c*_*max*_ = 1 (or locally slightly greater for sharp-edged patches embedded within a larger field). The semisaturation contrast parameter in Equation 10 is in light-adapted units and should be set to a relatively small value (*α*_*HB*_ ≪ 1) so that the ratio *c*^*n*_*HB*_^ /(*α*_*HB*_^*n*_*HB*_^ + *c*^*n*_*HB*_^) is close to saturation for *c* ≈ 1. The asymptotic limit *c* → ∞ has no physiological interpretation.

The hyperbolic-ratio model requires the extraction of two distinct pieces of information about the stimulus. One is the contrast c, which is an intrinsic property of the image. The other is the degree of match between the input template *I*_*T*_ and the weighting function of the neuron. This match is a relational property that depends not only on the image but also on the WF. Both pieces of information are available in the image but they cannot be extracted by the application of a single filter. The stimulus drive extracted by a linear filter (Eq. 5) is a single number that confounds contrast information and degree-of-match information. If either of them is known, the other can be decoded from the stimulus drive. The two models discussed so far are complementary in this regard: The linear model characterizes stimulus selectivity, whereas the hyperbolic-ratio model describes dependence on contrast. When an arbitrary stimulus is presented, however, both pieces of information are unknown and multiple filters must be applied to resolve this ambiguity. The divisive normalization model pools the (half-squared) outputs of filters with diverse tuning preferences in order to estimate the intrinsic properties of the image. This pooled estimate is then used to normalize the stimulus drives to the individual units.

### 2.3 Divisive Normalization Model

The notion of a *normalization pool* is pivotal to the *divisive normalization (DNM)* model and sets it apart from the simpler models discussed above. The introduction of a normalization pool is motivated by three convergent lines of evidence. The first line comes from the experimental data on the contrast-set gain control outlined above (Fig. 3), coupled with the need to pool across filters with diverse tuning preferences to estimate the stimulus contrast. A second, related line comes from a priori considerations involving the so-called *noise-saturation dilemma* (Grossberg, 1988). Individual neuronal responses are noisy and they have a limited dynamic range. The brain needs to represent signals across very wide dynamic ranges. Hence the dilemma: “If the [activations of individual neurons] are sensitive to large inputs, then why do not small inputs get lost in internal system noise? If the [activations] are sensitive to small inputs, then why do they not all saturate at their maximum values in response to large inputs?” (Grossberg, 1988, p. 33). The proposed solution relies on pooled inhibition^6^ within a network of interacting neurons to normalize the individual responses relative to a dynamically adjustable baseline. Such divisive normalization has been proposed (e.g., Carandini & Heeger, 2012) as a canonical type of neural computation in a wide variety of sensory modalities, brain regions, and species. The divisive nonlinearity in the hyperbolic-ratio model (Eq. 10) is an instantiation of this general principle.

The third and most direct line of evidence motivating the introduction of a normalization pool consists of experimental demonstrations of various broadly tuned suppressive effects in V1. In *cross-orientation suppression,* for example, the response to a grating (signal) is suppressed by another grating (mask) superimposed onto the signal within the neuron’s receptive field (e.g., DeAngelis, Robson, Ohzawa, & Freeman, 1992; Morrone, Burr, & Maffei, 1982; see Section 3.3.4 for further references and discussion). In *surround suppression,* the response is suppressed by masks presented outside the classical RF (e.g., Cavanaugh, Bair, & Movshon, 2002b; C.-Y. Li & Li, 1994; see Section 3.3.5).

Thus, modeling the responses of a single individual neuron requires filtering the input image with multiple linear filters that have diverse weighting functions. Let this diverse set be indexed by *i,* the filters in the normalization pool 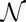 have preferences P_*i*_, and P* denote the tuning preference vector (Eq. 8) of the target neuron.

Heeger (1992b) combined all theoretical ideas introduced above into a single equation. His formulation of the divisive normalization model has been very influential (see Carandini & Heeger, 2012, for a recent review). In our notation, which is different from Heeger’s, this equation is

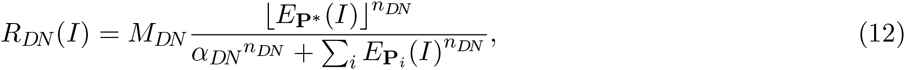

where the summation in the denominator encompasses the normalization pool 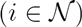. The exponent *n*_*DN*_ and the semi-saturation contrast *α*_*DN*_ are free parameters analogous to their counterparts in the hyperbolic-ratio Equation 10 (Fig. 4). Note that the firing-rate parameter *M*_*DN*_ is a constant that does not depend on stimulus *I*. This parameter determines the stimulus drive *E*_*P*\*_(*I*) to the target neuron, which can be either a simple or a complex cell (Eq. 9).

The *suppressive drive* 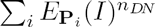 in the denominator represents the aggregated inhibitory influence impinging on the target neuron. There is evidence that this inhibitory influence combines lateral inhibition from other neurons in V1, feedforward inhibition from and within the lateral geniculate nucleus (LGN), and feedback inhibition from higher cortical areas (e.g., Sengpiel, Baddeley, Freeman, Harrad, & Blakemore, 1998; Angelucci & Shushruth, 2013; see Section 4.3 for further references and a brief discussion). These sources have different temporal and spatial properties (e.g., Bair, Cavanaugh, & Movshon, 2003; see Sect. 4.3). This is an active research area that is beyond our present scope. Equation 12 models this suppressive drive as a sum of (exponentiated) homogenous terms 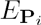, each of which is analogous to the stimulus drive *E*_*P\**_ in the numerator.

It is important to emphasize that this formula is a *functional* description only. It entails no commitment about what neurophysiological mechanisms produce the suppressive effect. The formula itself suggests a three-stage sequence of linear filtering followed by exponentiation followed by divisive normalization, and this is indeed how Equation 12 is implemented on a computer. This sequential scheme, however, is not physiologically possible because the un-normalized intermediate terms 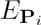 cannot be represented by substrates with a limited dynamic range such as membrane potentials or firing rates. This constraint is at the core of Grossberg’s (1988) noise-saturation dilemma. Instead, the normalization almost certainly involves dynamic inhibitory interactions within a recurrent network (e.g., Brosch & Neumann, 2014; Chance & Abbott, 2000; Ellias & Grossberg, 1975; Heeger, 1993; Kough & Poggio, 2008) in conjunction with other regulatory mechanisms (e.g., Carandini, Heeger, & Senn, 2002; Freeman, Durand, Kiper, & Carandini, 2002).

Various variants of Equation 12 have been used to account successfully for extracellular recordings (e.g., Heeger, 1992b; Carandini & Heeger, 2012), multi-electrode population recordings (e.g., Busse et al., 2009; Goris et al., 2009), and psychophysical data (e.g., Itti et al., 2000; Olzak & Thomas, 1999, 2003; To et al., 2010; Jesús & Laparra, 2010). Unfortunately, many of these applications use different mathematical formulations and idiosyncratic parameterizations. This practice makes it difficult to compare the results across studies despite the clear family resemblance of the model variants.

With the aim of consolidating this scattered literature, we propose that Equation 13 is used as the standard formulation of the divisive normalization model. *This is the main equation in the present article.* It is used in all of the simulations reported below. It was chosen on the basis of a theoretical analysis and the results of simulation experiments with several model variants that were compared informally on their ability to account for a comprehensive suite of empirical phenomena that are listed in Table 1 below. In our opinion, the following formulation achieves a good balance between flexibility and parsimony:

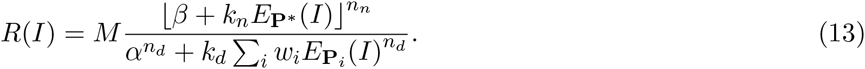

The firing-rate parameter *M* and the semisaturation contrast *α* have the same interpretation as their counterparts in Heeger’s (1992b) proposal (Eq. 12, Fig. 4). There are separate exponents *n*_*n*_ and *n*_*d*_ for the numerator and denominator, respectively. The *baseline* parameter *β* allows for non-zero responses when the stimulus drive *E*_P*\**_(*I*) is zero. The *maintained discharge* of the DNM neuron—its response to a blank stimulus (uniform gray field)—is 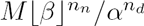 sps. Note that the firing rate of an actual simple cell in V1 can be less than its maintained discharge when the inhibitory regions in the cell’s receptive field are stimulated by a light spot (Hubel & Wiesel, 1959). This property can be modeled by Equation 13, assuming *β* > 0, but not by Equation 12. Conversely, setting *β* < 0 within the scope of the half-wave rectification operator 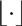 amounts to setting a threshold on the stimulus drive (the *iceberg effect,* Heeger, 1992a; Sceniak, Hawken, & Shapley, 2001; Tadmor & Tolhurst, 1989). The constraint that *β* must be fixed for a given neuron entails falsifiable predictions for the model (see Section 3.3.2 and Appendix C).

The *calibration constants k*_*n*_ and *k*_*d*_ are not free parameters. Conceptually, they are factored into the weights *w*_*i*_ in Equation 13 and the weighting function of the linear filter *E*_P*\**_ (Eq. 5). Both constants are defined with respect to a single *calibration image I_cal_*:

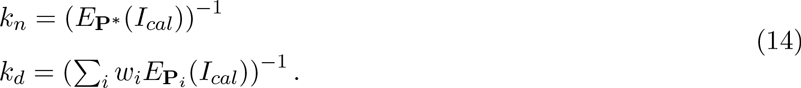

The calibration image is chosen a priori as the grating whose frequency and orientation (and phase for a simple cell) match the preferences P* of the target neuron. The contrast of *I*_*cal*_ is 1 and its spatial extent is large enough to fill both the classical receptive field and the suppressive surround. Calibrating the model in terms of an explicit image is convenient because it establishes a standardized scale for the substantive parameters *α* and *β*—they can be interpreted as equivalent contrasts. To see why, consider the *calibration family* of gratings 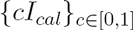 that use *I*_*cal*_ as a template and sweep a range of contrasts *c*. For this special family, Equations 13 and 14 reduce to the following variant of the hyperbolic-ratio model (Eq. 10, see also Appendix C):

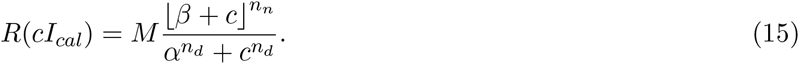

Then a positive baseline (*β* > 0) can be interpreted as the contrast of the counter-phase grating that cancels the maintained discharge of a simple cell, and a negative baseline (*β* < 0) as the contrast of the preferred (in-phase) grating that barely elicits a response.

To complete the specification of the divisive normalization model, we need to define the suppressive drive 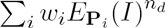 in the denominator of Equation 13. We need to specify three things: the weighting functions of the linear filters 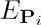, the composition of the normalization pool 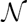 encompassed by the sum, and the pooling weights *w_i_.* We followed the common practice (e.g., Itti et al., 2000; Reynolds & Heeger, 2009) with respect to all three. First, we assume all linear filters have the same spatial-frequency bandwidth *h*_*f*_ (Eq. 4) and the same orientation bandwidth *h_θ_* (Eq. 3). These *h*_*f*_ and *h_θ_* are assumed for the stimulus drive *E_P*_* as well. Second, we assume the normalization pool tiles the space of frequencies and orientations. There is evidence (DeAngelis et al., 1992; DeAngelis, Freeman, & Ohzawa, 1994) that the suppressive effects are approximately invariant with respect to the phase of the mask grating. We model this by including only phase-invariant components *E*_*C*_ (Eq. 7) into 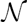. Third, we adopt the common simplifying assumption that the pooling weights *w*_*i*_ can be separated^7^ into independent pooling kernels with respect to space, frequency, and orientation:

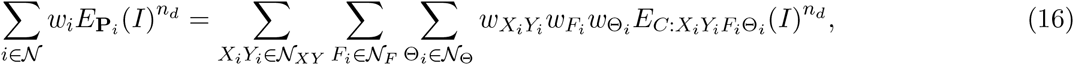

where 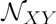 is a grid of image locations, 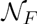 is a set of *frequency channels,* and 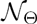 is a set of *orientation channels.* This specification is constrained by empirical data on various forms of suppression (surveyed in Section 3.3) and by considerations of symmetry, parsimony, and computational efficiency.

The spatial pooling weights 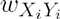 are defined by a radially symmetric 2D Gaussian kernel

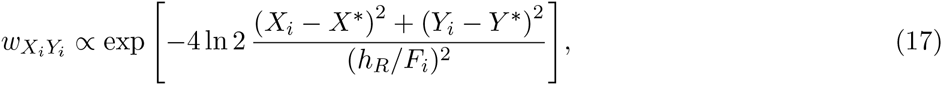

where *X*\* and *Y*\* are the coordinates of the center of the receptive field of the target neuron. The weights are defined up to a scaling factor and then calibrated by *k*_*d*_ in Equation 14. The diameter at half height of the kernel is proportional to the wavelength *1/F_i_* of each frequency channel. The *spatial pooling bandwidth h_R_* (in number of cycles) is a free parameter common to all channels. Note that the overall *suppressive field* of the model is produced by a combination of two types of spatial summation. First, the individual components 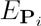 perform summation within elliptical Gabor receptive fields whose spatial dimensions 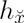 and 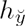 also are proportional to the respective channel wavelengths (Eqs. 3 and 4). Second, after nonlinear rectification with exponent n_*d*_, there is another summation across components centered on multiple locations *X_i_Y_i_.* Equation 17 defines the weighting function of the latter summation.

The frequency pooling weights 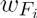 are defined by a Gaussian kernel along the log-frequency (octave) dimension:

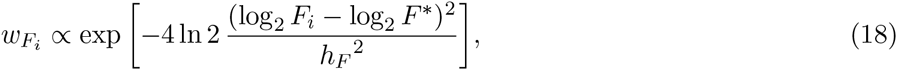

where *F*\* is the preferred frequency of the target neuron. The *frequency pooling bandwidth h_F_* (in octaves) controls the full width at half height of the kernel. This parameter is distinct from the bandwidth *h*_*f*_ of the weighting functions of the individual components 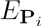. Because of the pooling, the frequency tuning of the suppressive effects is broader than *h_f_,* in agreement with the data (DeAngelis et al., 1994; C.-Y. Li & Li, 1994).

Finally, the orientation pooling weights 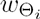 are defined by a von Mises kernel

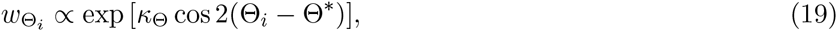

where Θ* is the preferred orientation of the target neuron. The von Mises distribution is the circular analog of the normal distribution (Fisher, 1996). The dimensionless *concentration parameter* 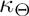 is intuitively similar to inverse variance. Equation 19 assigns 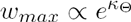 to the preferred orientation and 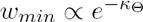 to the orthogonal orientation. The two points whose height is halfway between these two extremes occur at orientations 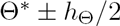, where

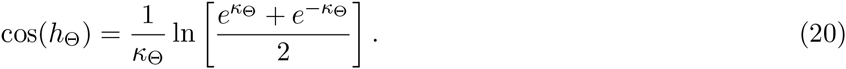

This equation establishes an invertible relationship in which 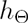 monotonically increases as 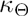 decreases. The circular uniform distribution 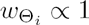 is the special case for 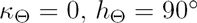. It is convenient to parameterize the DNM in terms of its *orientation pooling band*w*idth* 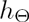 (in degrees). Again, this parameter is distinct from the bandwidth *h_θ_* of the weighting functions of the individual components 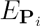 Because of the pooling, the tuning of cross-orientation suppression is broader than *h_θ_,* in agreement with the data(DeAngelis et al., 1992; Morrone et al., 1982). Unfortunately, 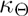 cannot be expressed as a closed-form function of 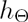, but in practice, Equation 20 is easy to solve numerically.

Overall, our formulation of the divisive normalization model has ten free parameters: *M, α, β, n_n_*, and *n*_*d*_ in Equation 13, the tuning bandwidths *h*_*f*_ and *h*_*θ*_ of the linear filters (Eq. 5), and the pooling bandwidths *h*_*R*_, *h*_*F*_, and *h*_Θ_ of the suppressive drive (Eq. 16). Five auxiliary constants 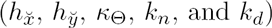 are calculated from the free parameters (cf. Table 2 below).

## 3 Simulation Experiments

The divisive normalization model (DNM, Eq. 13) was implemented as software and tested on a wide range of stimuli designed to replicate a comprehensive suite of published neurophysiological studies (Table 1).

### 3.1 Computational Implementation and Calibration

We developed software for Matlab (The MathWorks, 2015). This software takes a static grayscale image as input and produces a matrix of firing-rate responses for a population of DNM neurons. The neurons in this population have receptive fields at a single retinal location and they are tuned for a range of orientations and spatial frequencies. The details of this implementation are given in Appendix E. Briefly, the software provides tools for accomplishing two main computational tasks: constructing a DNM object for a given parameter set and calculating the responses of a model for a given input image.

A DNM object is a data structure that encapsulates the model’s parameters, weighting functions (WFs) for all linear filters, pooling weights for the suppressive drive, and various auxiliary information. Our implementation used 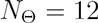 orientation channels spaced evenly at 15° increments around a circle. It used the following set of spatial-frequency channels: 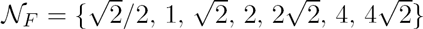 cyc/deg. The first and last channels in this set were auxiliary. They were included to contribute to the normalization of the five main channels in the middle. The WF of each channel was a Gabor function (Eq. 2). Two WFs were constructed per channel: one in sine (Φ = 90°) and one in cosine (Φ = 0°) phase (Fig. 1), for a total of 168 = 12 × 7 × 2 Gabor patches. The sizes 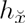 and 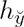 of their Gaussian envelopes were calculated from the free parameters *h*_*f*_ and *h_θ_* according to Equations 4 and 3. All images were rendered on a square 128 × 128 grid subtending 5.76 deg of visual angle. To improve efficiency, we also used a “small” 64 × 64 grid for some simulations that did not involve stimuli with extensive surrounds.

The calibration constants *k*_*n*_ and *k*_*d*_ were calculated according to Equation 14 with the aid of a calibration grating *I*_*cal*_ with unit contrast, vertical orientation (Θ* = 0°), frequency *F*\* = 2.0 cyc/deg, phase Φ* = 0°, and spatial extent covering the entire grid.

The calibrated model can be applied to an arbitrary grayscale input image. The computationally expensive operation is the calculation of the suppressive drive in Equation 16. The stimulus is convolved with each of the 168 Gabor filters to produce 84 phase-invariant suppressive terms 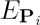 in Equation 13. The software uses the FFT algorithm (Fast Fourier Transform, Lathi, 2005) to compute these convolutions efficiently. See Appendix E for details. The suppressive drives of multiple simulated neurons can be computed as weighted linear combinations of the same 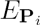 with different pooling kernels 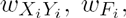 and 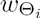 centered on different orientations and frequencies (Eqs. 16–19). Our software calculates the responses of 60 DNM complex cells and 240 DNM simple cells. The former are phase invariant (Eq. 7) and span the 12 orientations × 5 frequencies listed above (excluding the auxiliary frequencies). The latter vary also in phase: Φ = 0°, 90°, 180°, and 270°.

The channel with Θ* = 0° and *F*\* = 2.0 cyc/deg is singled out as the *target* and used to generate the DNM predictions in most figures below. Note, however, that it is only marginally more expensive to compute an entire *population code* of the input image (deCharms & Zador, 2000; Pouget, Dayan, & Zemel, 2003). Our software can be used as off-the-shelf front-end to larger models utilizing population codes. In fact, an earlier version of the software has been incorporated into such larger models (Petrov et al., 2005, 2006; Jacobs, 2009).

### 3.2 Standard Parameter Set

We propose the values listed in Table 2 as a standard parameterization of the DNM. These values were used to generate almost all of the DNM results in this article, with a few exceptions noted explicitly below. They are compatible with typical neurophysiological measurements of representative simple and complex cells, and with the phenomena in Table 1. Thus, these values are good defaults when the DNM is used as a building block for the construction of larger models of the visual system. The explicit reliance on a calibration image (Eq. 14) is designed to facilitate the reusability of parameter values across multiple applications.

A limitation inherent in the notion of standard parameterization must be acknowledged: Real V1 neurons have diverse properties that are impossible to subsume under a single parameter set. We reviewed single-cell recordings from different species (e.g., cats, new/old world monkeys, rabbits, rodents, and ferrets) obtained under different conditions (e.g., anesthesia vs. alertness, see Bereshpolova et al., 2011; Chen, Martinez-conde, Macknik, Swadlow, & Alonso, 2009; Disney, Aoki, & Hawken, 2007; Ecker et al., 2014; Goltstein, Montijn, & Pennartz, 2015; Niell & Stryker, 2010) and different experimental protocols (see Mukherjee & Kaplan, 1995; Smyth, Willmore, Baker, Thompson, & Tolhurst, 2003, for examples). Furthermore, there is substantial variability within a sample of neurons recorded from a single animal under constant conditions. Clearly, model parameters need to vary substantially to accommodate this diversity. The standard set in Table 2 is proposed as an estimate of the central tendency of a broad distribution. Our goal is to produce a *qualitative* account of the phenomena in Table 1, rather than a *quantitative* fit of a specific data set.

Now, let us discuss briefly the ten free parameters in Table 2. The firing-rate parameter *M* converts the dimensionless ratio of Equation 13 into physiologically observable units (spikes/sec). This parameter plays no role in accounting for the qualitative patterns that are our focus here, but is indispensable for quantitative fitting of actual neuronal firing rates. The semi-saturation contrast *α* was discussed in Section 2.2 and illustrated in Figure 4B. Under the DNM calibration (Eq. 13), it is expressed in dimensionless units and can be interpreted as equivalent contrast. The proposed value (*α* = 0.1) is consistent with empirical estimates obtained via hyperbolic-ratio fits to the contrast response functions of V1 neurons (Busse et al., 2009; Albrecht & Hamilton, 1982; Sclar et al., 1990; Gardner, Anzai, Ohzawa, & Freeman, 1999a). The exponents *n*_*n*_ and n*d* control the slope of the CF (Fig. 4A). Exponents greater than 1 are needed to account for the accelerating nonlinearity at low contrasts discussed in Appendix B. The standard value *n*_*n*_ = *n*_*d*_ = 2 is consistent with empirical estimates (e.g., Busse et al., 2009, p. 933; Albrecht et al., 2003, p. 752) and implements Heeger’s (1992a) half-squaring operator 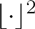. Note that Equation 13 has separate exponents for the numerator and denominator. This additional flexibility is needed for quantitative fits of physiological (e.g., Carandini & Heeger, 2012) and psychophysical (e.g., Itti et al., 2000) data.

The baseline parameter *β* can be interpreted as equivalent contrast as discussed in Section 2.3. The interpretation depends on its sign. Negative values effectively impose a threshold on the stimulus drive, whereas positive values produce a *maintained discharge* in response to a blank stimulus (uniform gray field). Many real neurons in V1 emit spikes in the absence of external stimulation, although the spontaneous firing rates typically are quite low (Allison & Bonds, 1994; Hubel & Wiesel, 1959; Hubel, 1959; R. L. De Valois, Albrecht, & Thorell, 1982, 1982; Pettigrew, Nikara, & Bishop, 1968; Squatrito, Trotter, & Poggio, 1990). For example, Hubel and Wiesel (1959) recorded maintained discharges between 0.1 and 10 spikes/sec in V1 neurons of anesthetized cats. Under the standard parameterization, Equation 13 produces 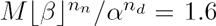 spikes/sec. The *β* parameter is in the focus of the mathematical analyses below (see Eq. 21 and Appendix C) and several simulations (e.g., Fig. 10) that explore non-standard values.

The remaining five free parameters control various bandwidths. They have diverse units (listed in Table 2) and should be interpreted with care because their empirically observable analogs depend on complex interactions among the DNM components as discussed in Section 3.3.3 below. For instance, the orientation tuning bandwidth of the model as a whole is 31.8° under the standard parameters (Fig. 13 and 14). Note that this is smaller than the bandwidth *h_θ_ =* 40° of the linear filtering stage and much smaller than the pooling bandwidth 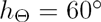 of the suppressive drive. Analogously, the frequency tuning bandwidth of the standard model as a whole is 1.11 oct, which is smaller than both *h_f_ =* 1.5 and *h_F_ =* 2.0 oct. The overall model bandwidths are within the reported ranges of neurophysiological measurements (R. L. De Valois, Albrecht, & Thorell, 1982; Kulikowski & Bishop, 1981; Movshon, Thompson, & Tolhurst, 1978a; Rose & Blakemore, 1974; Busse et al., 2009; Watkins & Berkley, 1974; Schiller, Finlay, & Volman, 1976b, 1976a). Finally, the standard FWHH of the radial spatial-pooling kernel (Eq. 17) covers *h*_*R*_ = 2.0 cycles of the neuron’s preferred frequency. This value is consistent with surround-suppression measurements (see Sections 3.3.1 and 3.3.5 for references).

### 3.3 Results

A systematic series of simulation experiments replicated the qualitative patterns characterizing the phenomena in Table 1. The simulated DNM patterns are plotted below alongside single-cell recording data from representative experiments. The physiological data were captured from the figures in their respective publications using PlotDigitizer (http://plotdigitizer.sourceforge.net) and replotted here in a unified format. All simulations used the standard parameters listed in Table 2 unless explicitly indicated otherwise.

#### 3.3.1 Size tuning

Our first simulation measured the responses of a DNM complex cell as a function of the stimulus diameter. All stimuli were gratings with orientation Θ* = 0° and frequency *F* =* 2 cpd that matched the cell’s tuning preferences. The resulting *size tuning* function is plotted in Figure 5A for gratings with maximal contrast *c* = 1. The responses increased as stimulus size increased at first, reached a peak, decreased, and finally settled to an asymptote. This matches the qualitative pattern observed in single-cell recordings of V1 neurons (Gieselmann & Thiele, 2008; H. E. Jones, Grieve, Wang, & Sillito, 2001; Sengpiel et al., 1997; Schwabe et al., 2010).

**Figure 5:**
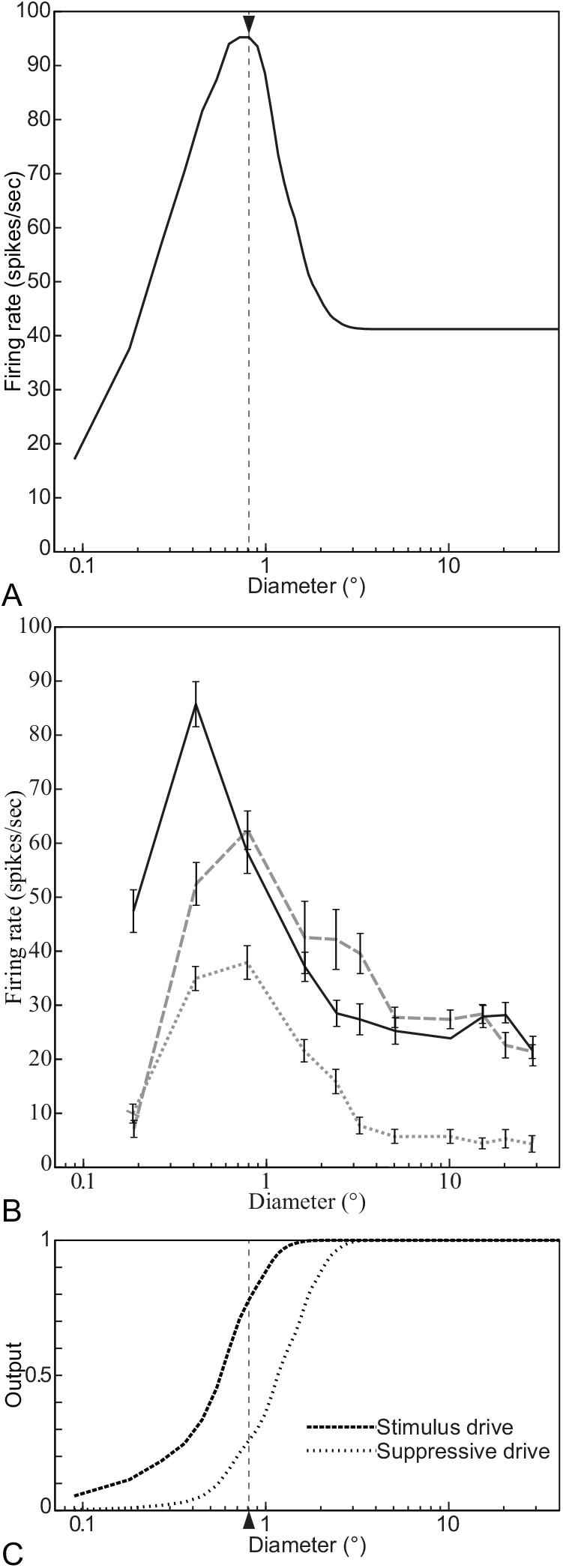
Size tuning functions of (A) a DNM complex cell with standard parameters, (B) three V1 complex cells (replotted from Schwabe et al., 2010, Fig. 2, anesthetized macaque, error bars = ±SE), and (C) the stimulus- and suppressive-drive terms of the DNM equation. The *measured RF diameter* of the DNM neuron is 0.81°· (indicated by the solid triangle in panel A). All stimuli were gratings with maximal contrast (*c* = 1) and orientation and frequency that matched the preferences of the respective neuron. (DNM = divisive normalization model. See Phenomenon 1 in Table 1.)

The non-monotonic response pattern in Figure 5 indicates that the neuron’s receptive field has limited spatial extent (Phenomenon 1 in Table 1). The diameter of the grating that induces the maximal response is often used in physiological studies to operationalize the size of the classical receptive field. For the DNM neuron, this *measured RF diameter* is MRFD ≈ 0.81° (marked by the black triangle in Figure 5A). Note that it is narrower than the FWHH sizes of the elliptical contour of the weighting function of the model’s linear stage (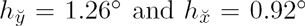, Table 2, see also Fig. 1). Other parameter sets were also tested and the MRFD tended to expand as 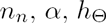, or *h*_*F*_ increased or 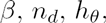 or *h*_*f*_ decreased. As a consistency check, note also that the measured asymptotic response rate (≈ 41.2 spikes/sec) matched the prediction of the hyperbolic-ratio Equation 15. This is because the gratings with very large diameters in this simulation became identical to the calibration image *I*_*cal*_, thereby satisfying the condition for applicability of Equation 15.

The nonmonotonic response pattern in the model arises from the interplay between the numerator and denominator in Equation 13. As the stimulus diameter increases, the stimulus drive (Eq. 9) rises faster but saturates earlier than the suppressive drive (Eq. 16, Fig. 5C, Gieselmann & Thiele, 2008).

The measured RF diameters of real neurons depend on the parameters of the grating. The MRFD increases as the stimulus luminance contrast decreases (Phenomenon 2, Figure 6B, Schwabe et al., 2010; Sengpiel et al., 1997; Sceniak, Ringach, Hawken, & Shapley, 1999; Kapadia, Westheimer, & Gilbert, 1999; Cavanaugh, Bair, & Movshon, 2002a; Tailby, Solomon, Peirce, & Metha, 2007; Nienborg et al., 2013). Also, the MRFD decreases for gratings with non-preferred orientations (Ph. 3, Fig. 7B, Tailby et al., 2007). The DNM neuron reproduces both phenomena (Figures 6A and 7A). The simulation that produced Figure 7A was designed to emulate the method of Tailby et al. (2007). Specifically, the stimuli in the Opt condition (gray solid line) were gratings of the preferred orientation (0°). We determined the MRFD for the DNM neuron in the Opt condition (0.81°, vertical dashed line) and then determined the orientation tuning curve for gratings with this diameter (Fig. 13A). The orientation in the Ori_∆_ condition (15.9°) was determined by the half-height points of this (symmetric) tuning curve. Probing the DNM neuron with gratings at this sub-optimal orientation produced the dotted black line in Figure 7A. It replicates the qualitative pattern of the neurophysiological data in panel B.

**Figure 6:**
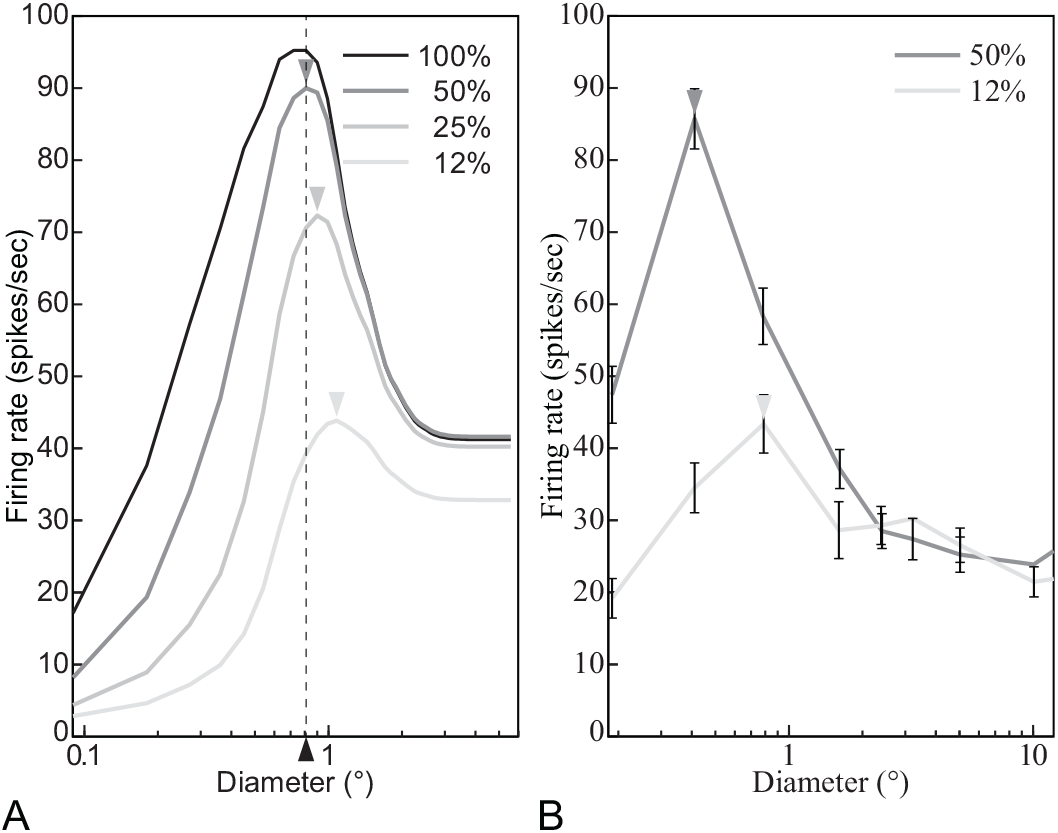
The size tuning function depends on the contrast of the stimulus grating. (A) DNM complex cell with standard parameters, (B) a V1 complex cell (replotted from Schwabe et al., 2010, Figure 2a, anesthetized macaque, error bars = ±SE). The measured RF diameter (depicted by triangles) increases as the stimulus contrast (shown in the legend) decreases. Stimulus orientation and frequency matched the preferences of the respective neuron. (DNM = divisive normalization model. See Phenomenon 2 in Table 1).

**Figure 7:**
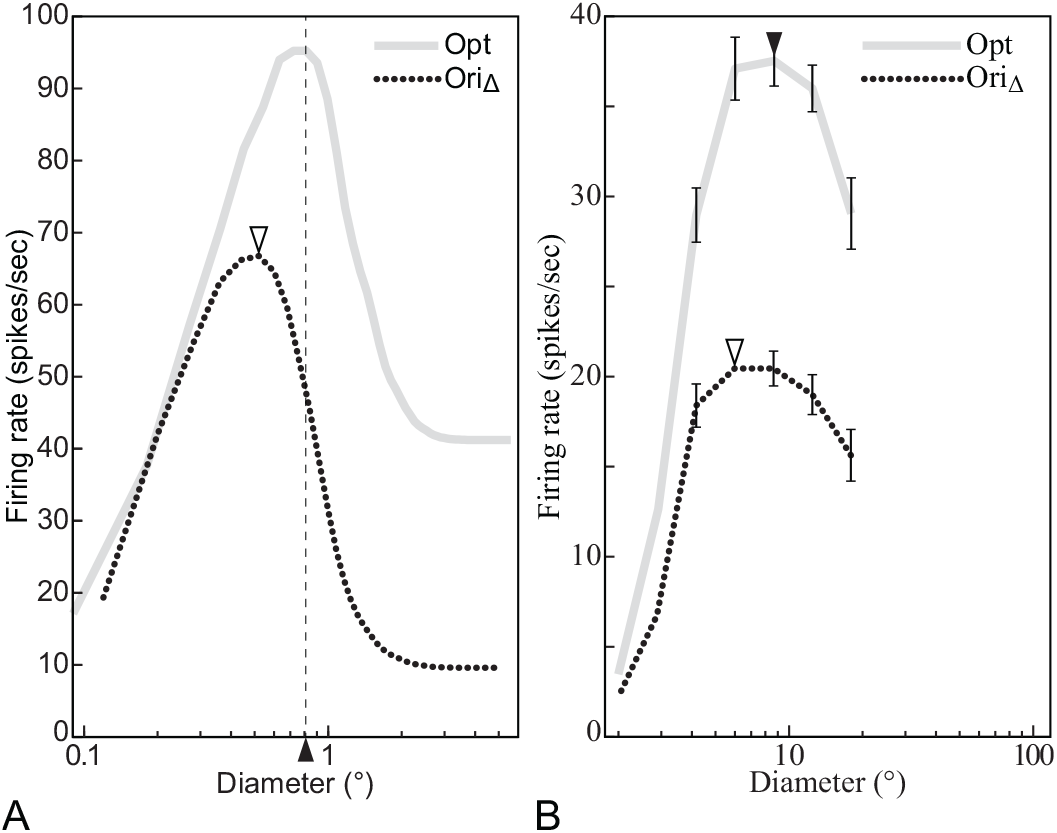
The size tuning function depends on the orientation of the stimulus grating. (A) DNM complex cell with standard parameters, (B) V1 complex cell (replotted from Tailby et al., 2007, Fig. 2b, anesthetized cat, error bars = ±SE). The grating orientation in the Opt condition matched the preferences of the respective neuron. The orientation in the Ori_∆_ condition was determined by the half-height point of the orientation tuning function (Tailby et al., 2007). See text for details. All gratings had maximal contrast (*c* = 1) and their frequency matched the preference of the respective neuron. (DNM = divisive normalization model. See Phenomenon 3 in Table 1.)

The MRFD of the model is also affected by the spatial frequency of the grating. It decreases if the frequency is either lower or higher than the DNM neuron’s preferred frequency, with stronger decreases for lower frequencies (Fig. 8A). The non-optimal stimulus frequencies (Spf_*L*_ = -0.21 oct, 0.86 cpd; Spf_*H*_ = 1.52 oct, 2.87 cpd) for this simulation were chosen at the half-height points of the model frequency-tuning curve (Fig. 13D; Tailby et al., 2007). The available recordings from real V1 neurons suggest that their MRFD tend to decrease as the stimulus frequency becomes higher than the preferred frequency, but no clear trend has been observed when the stimulus frequency becomes lower (Fig. 8B, Osaki, Naito, Sadakane, Okamoto, & Sato, 2011; see also Tailby et al., 2007). This qualitative pattern may be somewhat different than the DNM prediction. See Section 3.4 for further discussion of the relationships among stimulus size, frequency, and neuronal responses.

The size tuning of V1 neurons was also measured by using annular stimuli that overlay a circular grey “hole” in a larger circular grating with the neuron’s preferred orientation and frequency (H. E. Jones et al., 2001; Sengpiel et al., 1997). The recordings from V1 neurons decreased monotonically as the diameter of the hole increased, and then leveled off at an asymptotic level that was similar to the spontaneous discharge of the cell (Fig. 9B, Ph. 5). The hole diameter for which the responses become nearly constant is similar to the RF diameter measured with disc stimuli as discussed above. The DNM neuron shows analogous trends (Figure 9A). Recall that when disc stimuli (with no holes) are used to measure the RF diameter, the latter depends on stimulus contrast for both real and model neurons (Fig. 6). When annular stimuli are used, however, the DNM predicts approximate invariance with respect to the contrast of the grating inside the annular envelope (Phenomenon 6).

**Figure 8:**
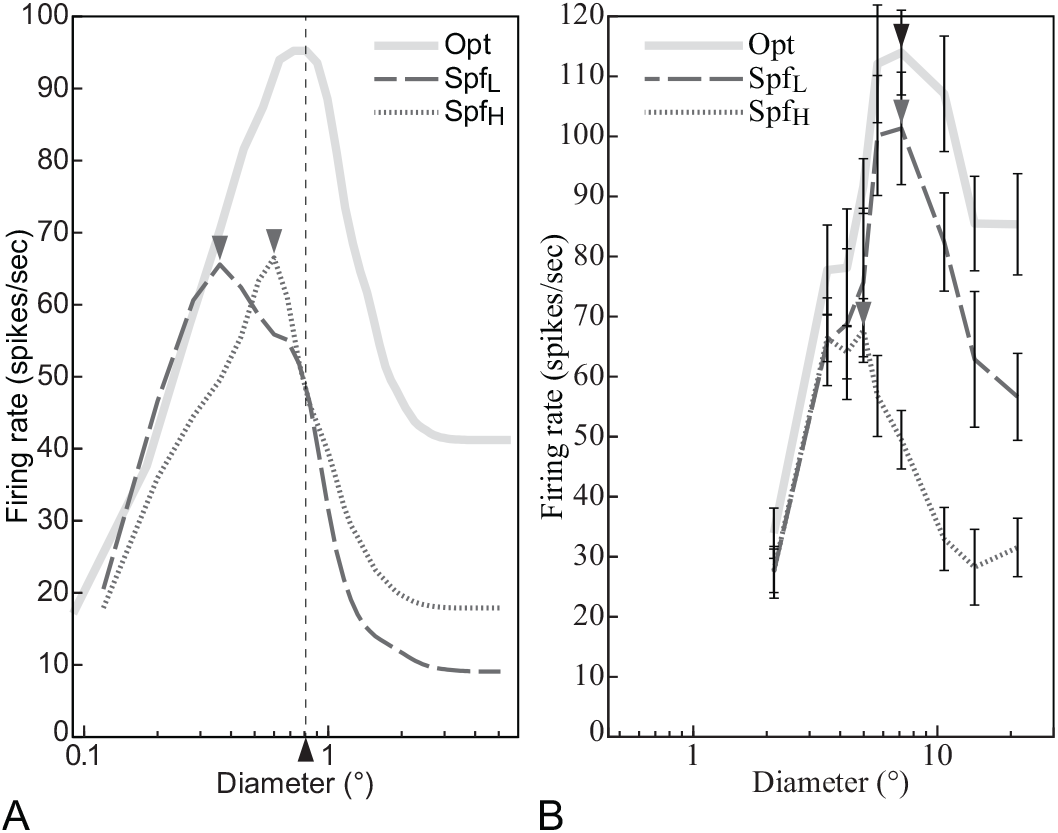
The size tuning function depends on the spatial frequency of the stimulus grating. (A) DNM complex cell with standard parameters, (B) a real V1 neuron (replotted from Osaki et al., 2011, Figures 2A and 4A, anesthetized cat, error bars = ±SE). The stimulus frequency matched the preference of the respective neuron in the Opt condition, was lower in the Spf_*L*_ condition, and higher in the Spf_*H*_ condition. The side frequencies in (A) were determined by the half-height points of the DNM frequency tuning curve. See text for details. The data in (B) were collected with Spf_*L*_ = 0.10 cpd, Opt = 0.20 cpd, and Spf_*H*_ = 0.30 cpd (Osaki et al., 2011). All gratings had maximal contrast (*c* = 1) and their orientation matched the preference of the respective neuron. (DNM = divisive normalization model. See Phenomenon 4 in Table 1.)

**Figure 9:**
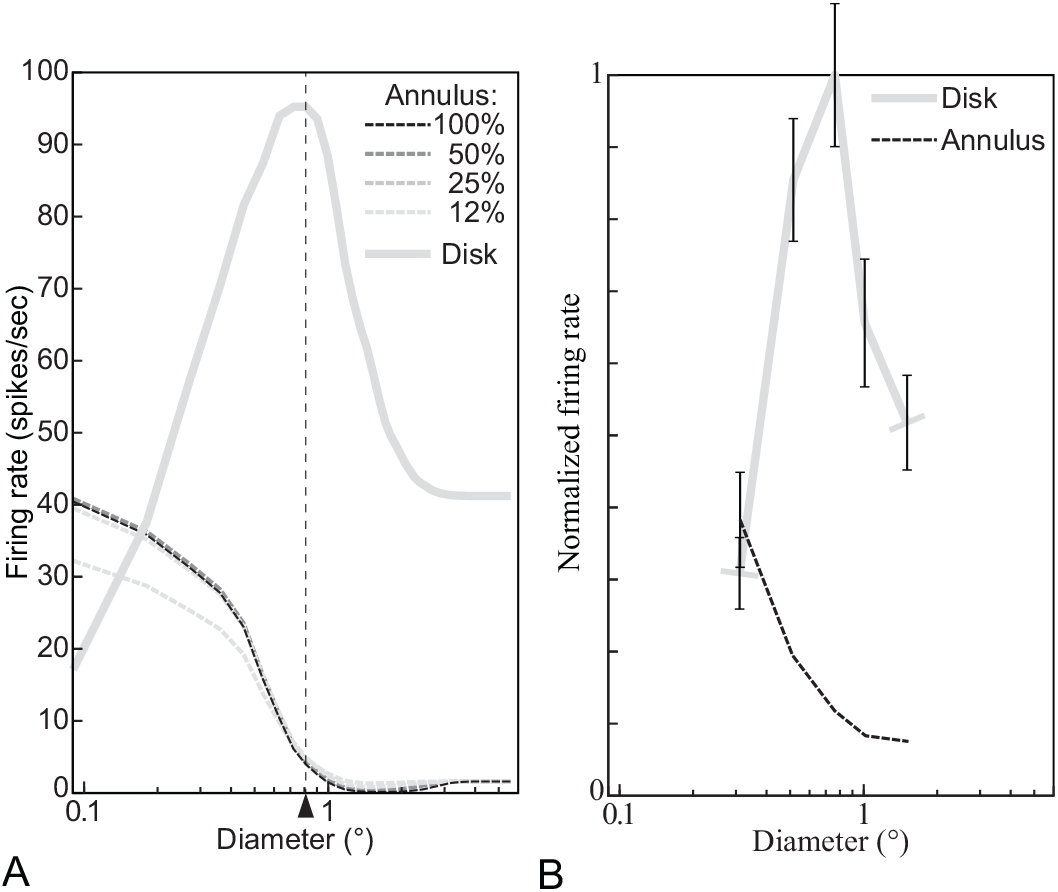
Responses to annular stimuli containing a circular grey “hole” inside a large circular grating. The X axis specifies the diameter of the hole for annular stimuli (dashed lines) and the outside diameter for disk stimuli (solid line, cf. Figure 5). (A) DNM complex cell with standard parameters. The legend indicates the contrast of the annular stimuli; the disk had 100% contrast. The measured RF diameter (0.81°) is indicated by a triangle. (B) V1 neuron (replotted from H. E. Jones et al., 2001, Fig. 1, anesthetized macaque, error bars = ±SE). All stimuli are based on gratings whose frequency and orientation matched the preference of the respective neuron. See Phenomena 5 and 6 in Table 1.

#### 3.3.2 Contrast sensitivity

The contrast sensitivity of the DNM neuron was measured using a grating whose orientation and spatial-frequency are those to which the neuron is tuned and its size is large enough (5.58°) to cover both the receptive field of the stimulus and the suppressive field of the model neuron. The model responses are plotted in Figure 10A (solid curve) as a function of the contrast (Phenomenon 7). The slope of the contrast sensitivity function becomes maximal when the contrast is at 9%, above 0 (see Sect. 2.2). It may be relevant to the psychophysical results that a differential threshold of a grating contrast becomes minimal above the contrast detection threshold (see Itti et al., 2000; Wilson, 1980 for a comparison between human- and model performance on luminance contrast discrimination).

**Figure 10:**
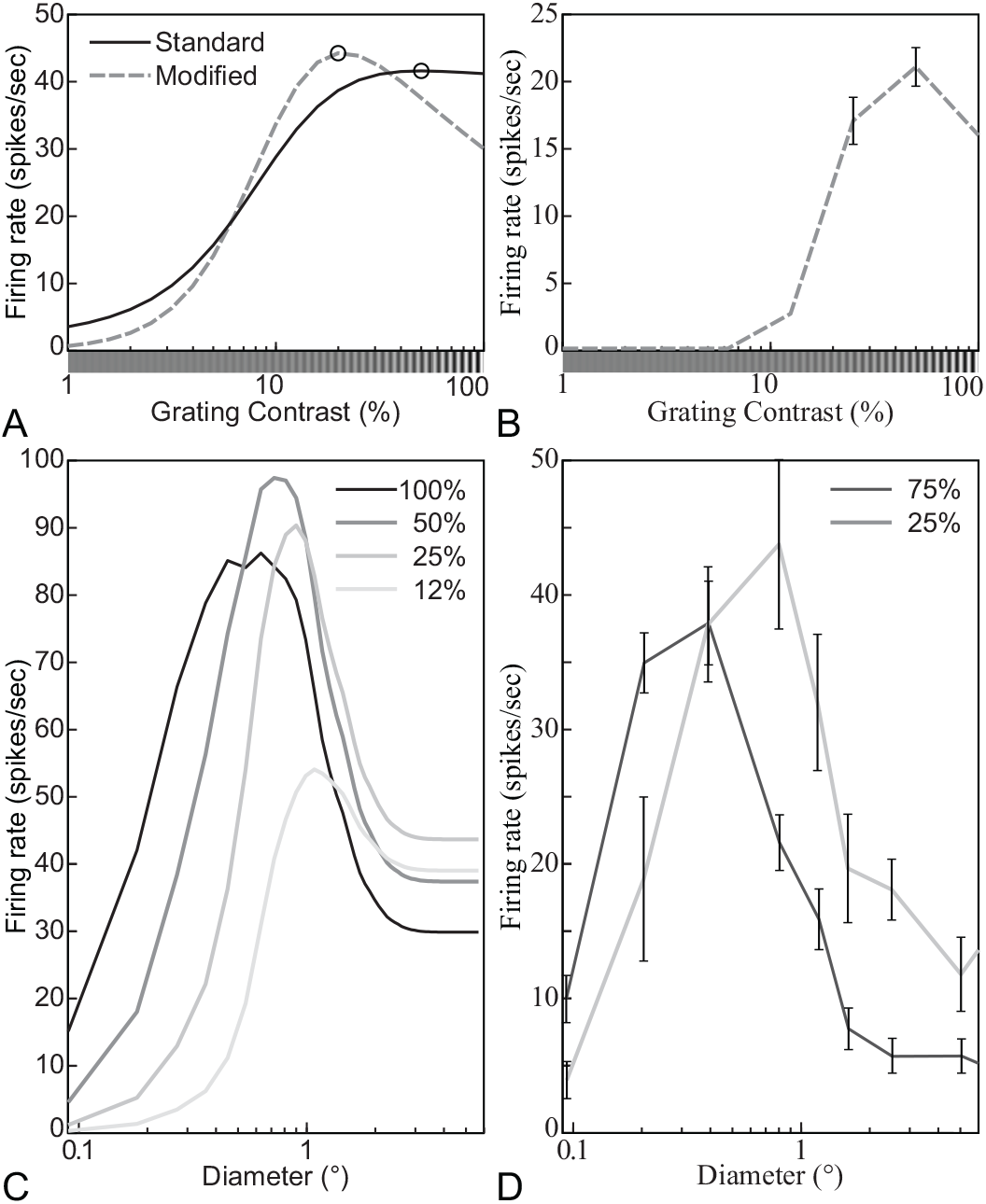
Examples of the *supersaturation effect* (Phenomenon 8 in Table 1). (A) Contrast sensitivity function of a DNM neuron with a standard (solid line) and a modified parameter set (*n*_*d*_ = 2.35, *β* = 0, *M* = 30, dashed line). (B) Contrast sensitivity function of a real neuron (replotted from Peirce, 2007, Fig. 2e, anesthetized macaque). (C) Size-tuning functions of the DNM neuron with the modified parameter set for different contrasts of a grating (shown in legend; see Fig. 6 for the standard-parameter analog). (D) Size-tuning functions of a V1 complex cell (replotted from Schwabe et al., 2010, Fig. 2C, anesthetized macaque, error bars = ±SE). Stimulus frequency and orientation matched the DNM preferences in (A) and (C). The size of the grating patch was 2.88° for (A).

Note that the contrast sensitivity function of the virtual neuron slightly decreases if the contrast is higher than 50% with the standard parameter set (indicated by a solid curve and an open circle in Figure 10A). An analogous trend is also observed from some real neurons (C.-Y. Li & Creutzfeldt, 1984; Tyler & Apkarian, 1985; Somers et al., 1998; Peirce, 2007). This is called the *supersaturation effect* (Ph. 8). This effect could be emulated only weakly by the DNM neuron with the standard parameter set. Some V1 neurons, however, do show the supersaturation effect very clearly (Figure 10B, D). Mathematical analysis (Appendix C) indicates that the model produces this effect when the baseline parameter *β* exceeds the following critical value:

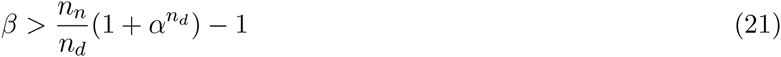

This inequality suggests that the super-saturation effect becomes stronger as *β* increases, *α* decreases, and *n_n_/n_d_* decreases (Figure 10A, C; see also Sawada & Petrov, 2015, December, for supersaturation plots produced with a different parameter set).

When visual noise is added to the preferred grating of a real neuron, the neuron’s responses tend to be scaled down by a multiplicative factor (Ph. 9, Carandini et al., 1997). This effect is observed in the model when the contrast of the tuned grating was high (e.g., 50, 25, and 12% in Figures 11A, B). When the contrast of the grating is zero or very low, adding noise increased the response of both real neurons (Squatrito et al., 1990) and the DNM neuron (Fig. 11A, B). Note that psychophysical results have shown that human performance on detecting a grating is improved by adding moderate amount of visual noise (e.g., 6% in Figure 11) to the grating (Moss, Ward, & Sannita, 2004; H. Sasaki et al., 2006; Simonotto et al., 1997). This is called *stochastic resonance.* The noise increased the response of the DNM neuron but decreased the slope of the contrast sensitivity function if the contrast of the grating was close to zero. Decreasing the slope is expected to increase the threshold of the model neuron for detecting the grating. However, if the output of one model neuron is relayed as excitatory input to another model neuron, it is possible that adding noise to the stimulus elevates the responses of the first neuron enough to trigger a response in the second neuron. In this way, moderate amounts of additive stimulus noise can improve the detection performance of the entire ensemble, despite the fact that the noise decreases the CF slope in the first stage.

**Figure 11:**
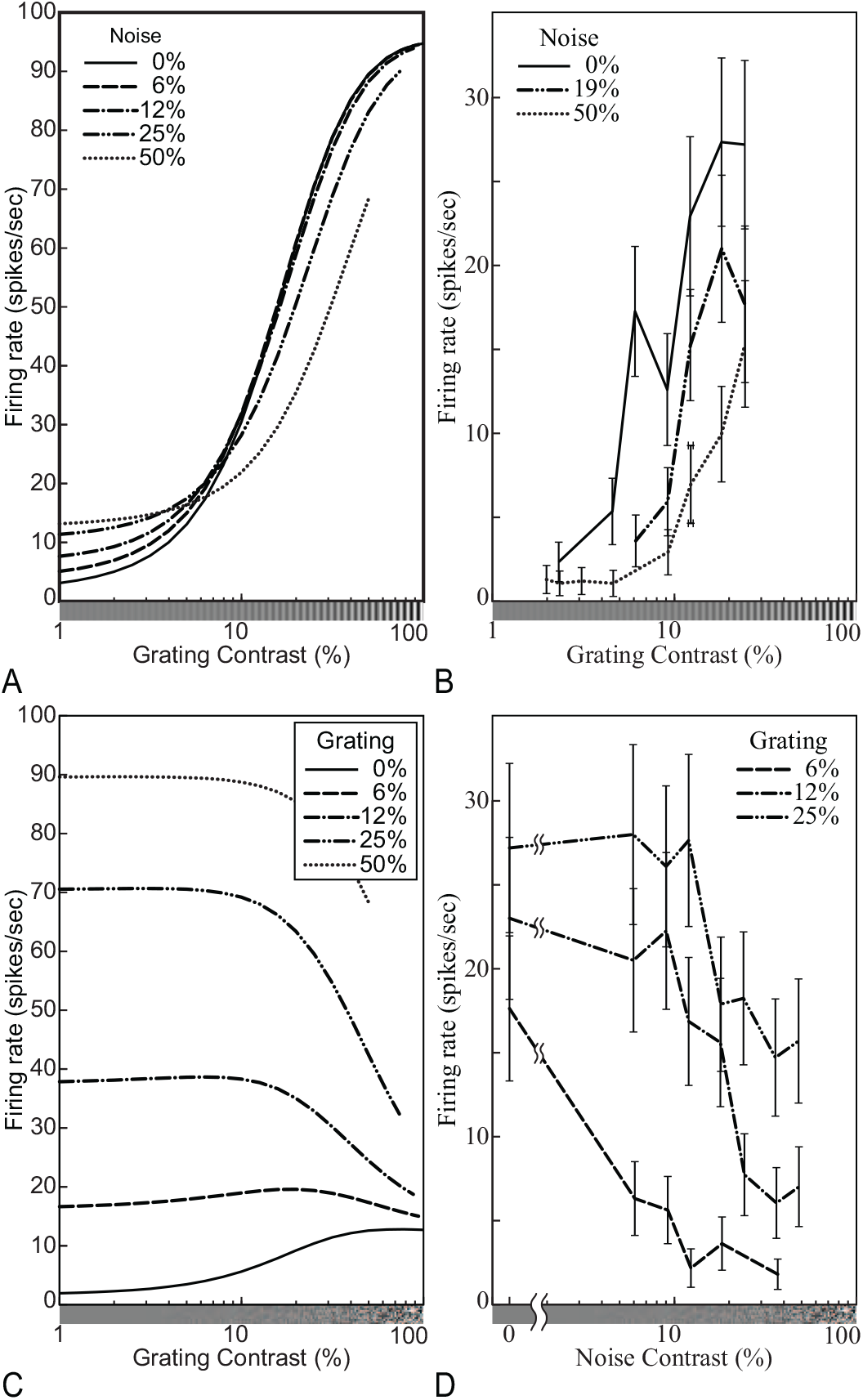
Responses of (A, C) the model neuron and (B, D) a V1 simple cell (replotted from Carandini et al., 1997, Fig. 14D and 14C, anesthetized macaque, error bars = ±SE) to a grating with a random noise pattern superimposed. The graphs in (A, B) are plotted against the contrast of the grating, whereas those in (C, D) are plotted against the contrast of the noise. (A, C) The size of the stimuli was equal to the measured RF diameter (0.81°), and their orientation and frequency matched the tuning preferences of the model neuron. See Phenomenon 9 in Table 1.

The contrast sensitivity functions of real neurons were affected by the orientation (Phenomenon 10, Tolhurst & Dean, 1991; Carandini et al., 1997) and the spatial-frequency (Phenomenon 11, Albrecht & Hamilton, 1982; Carandini et al., 1997; Albrecht et al., 2003) of the stimulus grating. The CFs of real V1 neurons were multiplicatively scaled down if the orientation or the frequency of the grating were different from the neuron’s preferred orientation or frequency (Fig. 3, Carandini et al., 1997). These trends are also observed in the model (Fig. 3B, D).

Our simulation experiments showed that the CF of the model neuron was also affected by the size of the grating patch (Fig. 12). As stimulus diameter increases up to 0.8°, the overall firing-rate increases, the CF slope at low contrasts becomes steeper, and the function becomes more saturating at high contrasts. Then, the asymptotic firing rate decreases while the slope at low contrasts is unchanged up to 3.2°. The function is unchanged even if stimulus diameter becomes larger than 3.2°. Note that this phenomenon was predicted by an analogous model of LGN neurons and was observed in a physiological experiment (Bonin, Mante, & Carandini, 2005). To the best of our knowledge, however, the phenomenon has never been observed in V1 neurons using grating patches as stimuli (see Schumer & Movshon, 1984). Note also that the supersaturation effect is observed from the DNM neuron if the stimulus diameter is 1.6° or larger.

**Figure 12:**
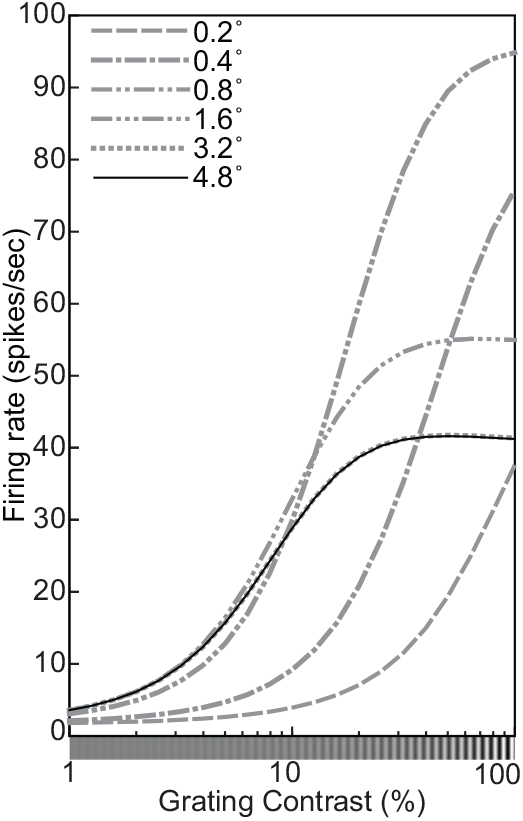
Contrast sensitivity functions of the model neuron for gratings with different diameters (indicated in the legend). The contrast, orientation, and spatial-frequency of the grating were 100%, 0°, and 2.0 cpd. See Phenomenon 12 in Table 1.

#### 3.3.3 Orientation and spatial-frequency tuning

The orientation and spatial-frequency tuning functions of the DNM neuron are shown in Figure 2 (Phenomena 13 and 14). The measured peaks of the tuning functions were consistent with the weighting function of the stimulus drive of the neuron. The measured bandwidths of the tuning functions were 31.8° in the orientation domain and 1.11 octaves in the frequency domain. Other parameter sets of the model were also tested and the measured bandwidths tended to become narrower as 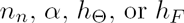 increase or 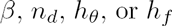 decrease. Note that the measured bandwidths were substantially narrower than those of the weighting function of the linear filter in Equation 5 (*h_θ_* = 40°, *h*_*f*_ = 1.5 oct). This narrowing effect is attributed to the rectification and the expansive nonlinearity (*n*_*n*_ > 1) in the numerator of Equation 13.

Note that the bandwidths of the numerator of Equation 13 are a little narrower (29.2° and 1.04 oct) than those of the DNM neuron itself (31.8° and 1.11 oct). This occurs because the denominator of Equation 13, which represents suppression, tends to widen the tuning curves of the DNM neuron. The tuning functions of the denominator are unimodal and become maximal at the same orientation and frequency as those of the numerator. The divisive operation then has a widening effect, and becomes stronger as the denominator becomes more sharply tuned. This is why the bandwidths of the DNM neuron become wider as 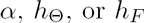 decrease or as *n*_*d*_ increases because they make the tuning functions of the denominator narrower. In particular, the 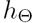 and 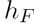 parameters (Table 2) allow an almost independent control of the widening of the orientation-and frequency tuning, respectively.

There is evidence that the orientation (Okamoto, Naito, Sadakane, Osaki, & Sato, 2009; C.-Y. Li & Li, 1994; see also Maffei & Fiorentini, 1976, for a report of a few exceptional neurons) and spatial-frequency (Osaki et al., 2011; Maffei & Fiorentini, 1976) bandwidths of real neurons become narrower when the size of the grating patch increases. The model simulated this effect well (Fig. 13A). This property of the model neuron is attributed to its stimulus drive. The bandwidths of the stimulus drive (and of the numerator) in the model also became narrower as the size of the patch increased (Fig. 13E, F). This suggests that the effect of the patch size on the tuning widths of the virtual neuron is *not* due to surround suppression. The weighting function of the linear stage of the model neuron is partially hidden by the surround suppression. This leads to a systematic underestimation of the measured receptive field (R. L. De Valois, Thorell, & Albrecht, 1985). Then, its measured bandwidths become wider if the grating patch just fills the underestimated RF. Note that the peak of the frequency tuning curve of real neurons tends to shift to a higher frequency as the size of the stimulus decreases (Teichert, Wachtler, Michler, Gail, & Eckhorn, 2007; Osaki et al., 2011). Note, however, that the model neuron did not show this trend. The peak of its frequency tuning curve was *not* affected by the size of the grating patch.

**Figure 13:**
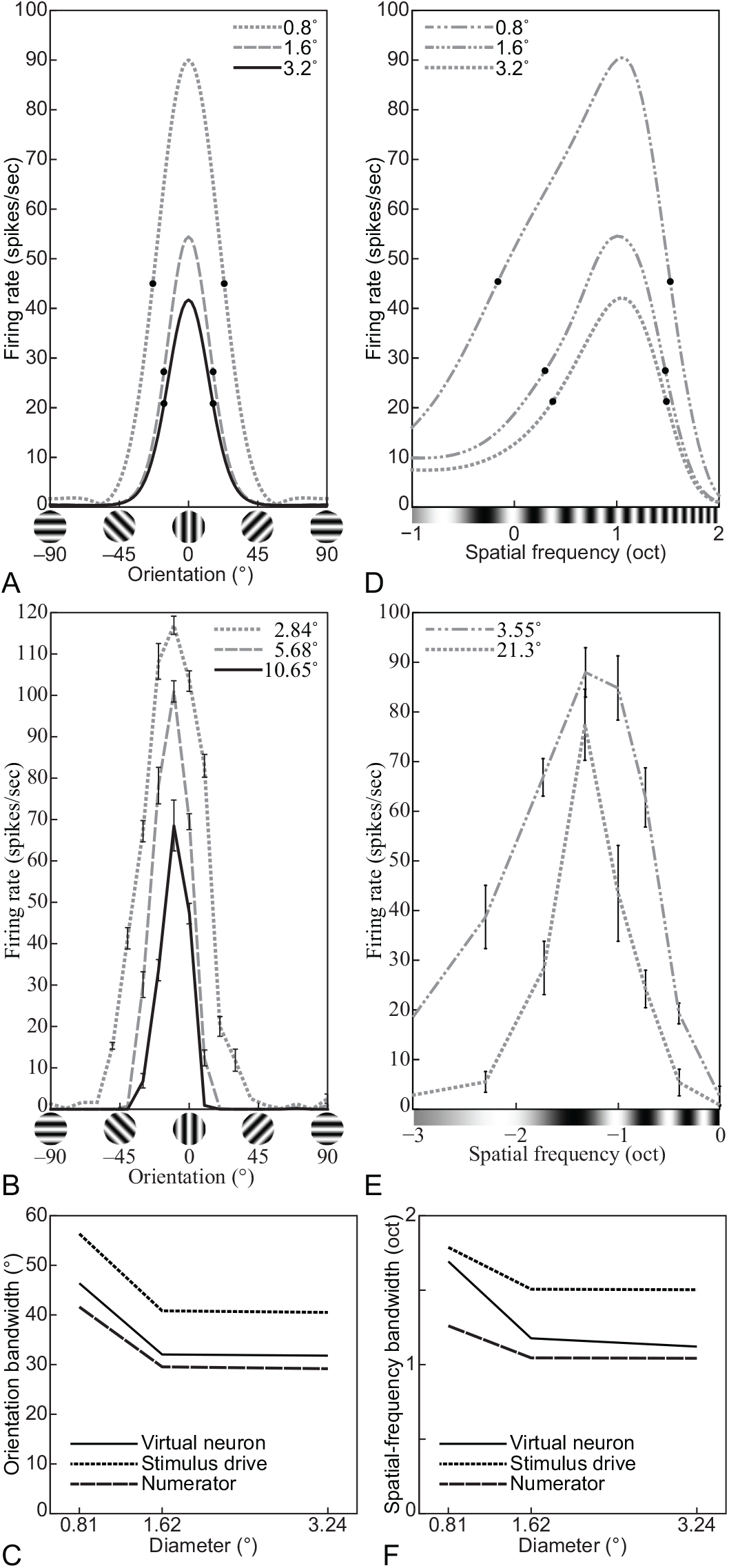
Orientation and spatial-frequency tuning functions of (A, D) the DNM neuron with standard parameters, (B) a V1 neuron (replotted from Okamoto et al., 2009, Fig. 1, anesthetized cat, error bars = ±SE), and (E) a V1 simple cell (replotted from Osaki et al., 2011, Fig. 5B, anesthetized cat, error bars = ±SE) for gratings with different diameters (indicated in legend). (C, F) The effect of the size on the bandwidths of the DNM neuron, its stimulus drive, and the numerator in Equation 13. The contrast of the grating was 100% for (A, D), its frequency was 2.0 cpd for (A), and its orientation was 0° for (D). (DNM = divisive normalization model. See Phenomena 15 and 16 in Table 1).

The orientation and spatial-frequency bandwidths of real neurons are affected by the contrast of the grating patches. As the contrast decreases, these bandwidths become slightly wider or narrower (or are invariant) depending on the individual neurons. Overall, the orientation bandwidths tend to be relatively invariant against the contrast (Phenomenon 17, Fig. 14A, B; Somers et al., 1995; Skottun et al., 1987; C.-Y. Li & Creutzfeldt, 1984; Sclar & Freeman, 1982; Anderson, Lampl, Gillespie, & Ferster, 2000; Alitto & Usrey, 2004; Troyer, Krukowski, Priebe, & Miller, 1998), but the spatial-frequency bandwidths tend to become narrower as the contrast decreases (Ph. 18, Fig. 14C, D; Sceniak et al., 2001; Albrecht & Hamilton, 1982; Skottun et al., 1987). These trends in the neurophysiological data could be emulated to some extent by the model neuron with a non-standard parameter set with smaller 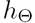 (40°) and *h*_*F*_ (1.0 oct). Under the standard parameterization, the bandwidths of the DNM neuron in both domains became slightly wider as the contrast decreased (Figure 14). Note that these trends in the neurophysiological data are rather weak and also note that there are many neurons in V1 that show opposite trends (Sclar & Freeman, 1982; Sceniak et al., 2001; Alitto & Usrey, 2004; Kim, 2011).

**Figure 14:**
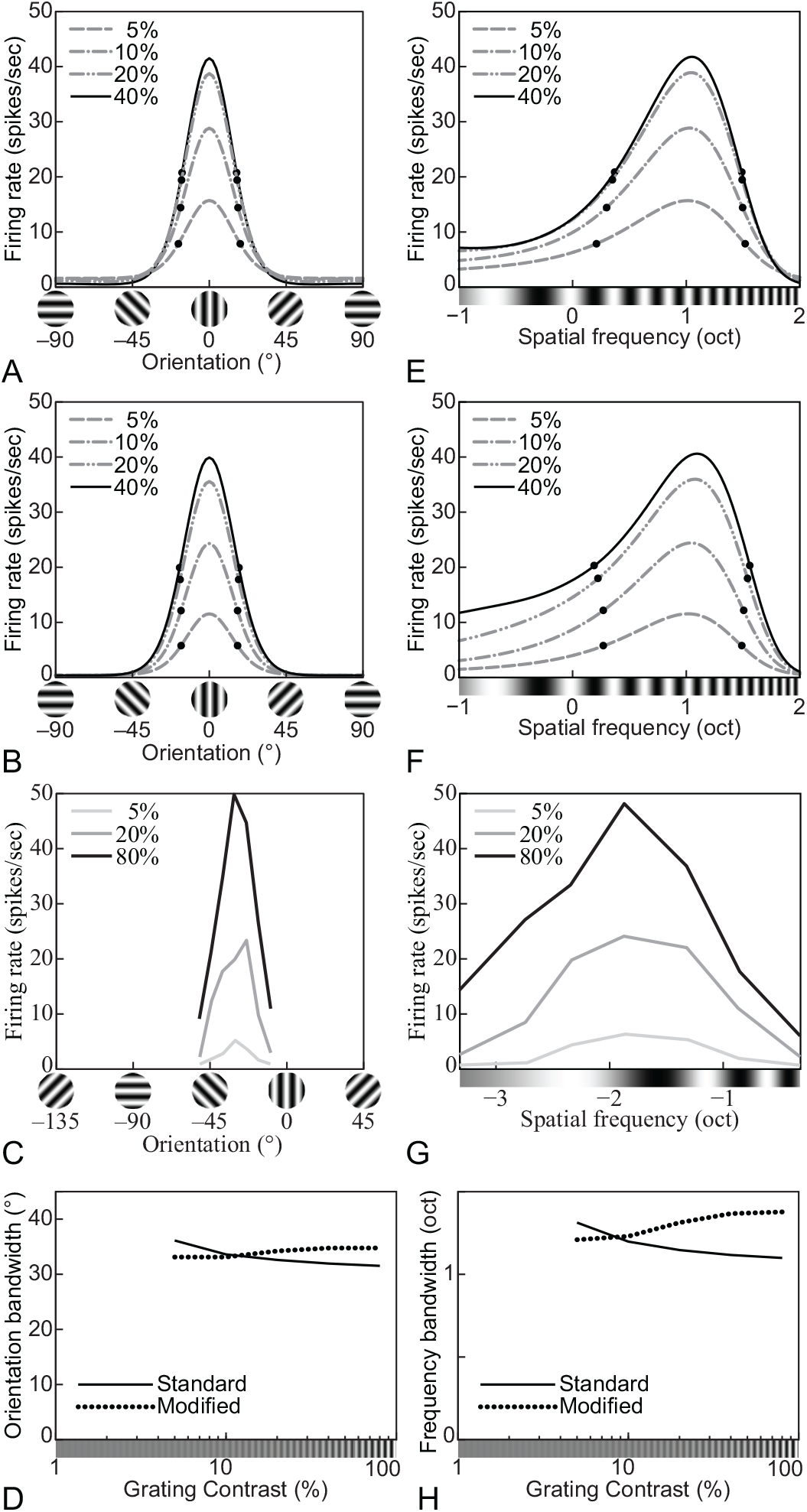
Orientation and spatial-frequency tuning functions for gratings with different contrasts (indicated in legend). (A, E) The DNM neuron with the standard parameter set; (B, F) The DNM neuron with a modified parameter set (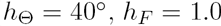 oct); (C) Orientation tuning of a simple striate cell (replotted from Skottun et al., 1987, Fig. 3A, anesthetized cat); (G) Frequency tuning of a simple striate cell (replotted from Skottun et al., 1987, Fig. 4A, anesthetized cat). (D, H) The effect of stimulus contrast on the bandwidths of the DNM neuron in the orientation (D) and frequency (H) domains for the two parameter sets (legend). (A, B, E, F) The size of the grating patch was 2.88°. The spatial-frequency of the grating was 2.0 cpd for (A, B) and its orientation was 0° for (E, F). (DNM = divisive normalization model. See Phenomena 17 and 18 in Table 1).

The effect of stimulus contrast on the orientation- and frequency-bandwidths depends on the *β* parameter in the numerator and on the denominator of Equation 13. The bandwidths of the numerator of the model neuron become narrower as the contrast decreases if *β* < 0 (the *iceberg effect,* Heeger, 1992a; Sceniak et al., 2001; Tadmor & Tolhurst, 1989) and become wider if *β* > 0. The bandwidths of the model neuron are also widened by the denominator depending on stimulus contrast. Recall that the widening effect of the denominator becomes stronger as the denominator becomes more sharply tuned. The tuning function of the denominator widens as the contrast decreases and it becomes a constant (*α^nd^*) at zero contrast. This contrast dependence of the widening effect of the denominator is consistent with the trend that the bandwidths of the model neuron become narrower as the contrast decreases.

Pollen and Ronner (1982) measured the spatial-frequency tuning of real neurons using sinusoidal and square gratings. They observed that, when measured with square gratings, the tuning function showed two peaks in the spatial-frequency domain (Phenomenon 19). The primary peak appeared at the neuron’s preferred frequency and the secondary peak appeared at one-third of the preferred frequency (Figure 15B). The height of the secondary peak was between 0.6 and 0.8 times the height of the primary peak for most neurons. The secondary peak of some neurons was even as high as the primary peak. These observations partly agree with the predictions of the linear model (Section 2.1). According to Fourier-series decomposition, a square grating with frequency *f_s_q* can be represented by a sum of sinusoidal gratings at the odd harmonics of *f*_*sq*_ with magnitudes proportional to reciprocals of the orders of the harmonics: 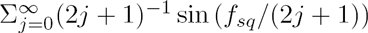. Thus, if a neuron satisfying the assumption of the linear model is tuned to a sinusoidal grating with frequency *f*_*sin*_, it is predicted that its tuning function to a square grating should show multiple local maxima in the spatial-frequency domain. The positions of these maxima are *f*_*sin*_/(2*j* + 1) and their heights are 1/(2*j* + 1) of the global maximum at *f*_*sin*_, where *j* is an integer. However, the results of Pollen and Ronner (1982) showed some deviations from the linear predictions. Only the primary (*j* = 0) and secondary peaks (*j* = 1) of the tuning functions could be reliably identified reliably in the data, and the heights of the observed secondary peaks were higher than the predicted height. The DNM neuron with a non-standard parameter set (*h*_*f*_ = 0.8, *h*_*F*_ = 0.4) can emulate these results, including the discrepancy to some extent (Figure 15A). The tuning function of the model neuron shows the secondary peak at one-third of the tuned frequency and its height is about 0.9 of the height of the primary peak. On the other hand, the model neuron also showed the tertiary peak at one-fifth of the tuned frequency (*j* = 2). Note also that the DNM neuron responds more strongly to the square grating than the sinusoidal grating, especially when their contrast is moderately low (Figure 15C). The contrast sensitivity function of the DNM neuron with the square grating shifts horizontally to the left (Phenomenon 20). This is because the outputs of both stimulus- and suppressive drives are always larger to the square grating than to the sinusoidal grating with the same frequency and orientation (see Equation 5).^8^

**Figure 15:**
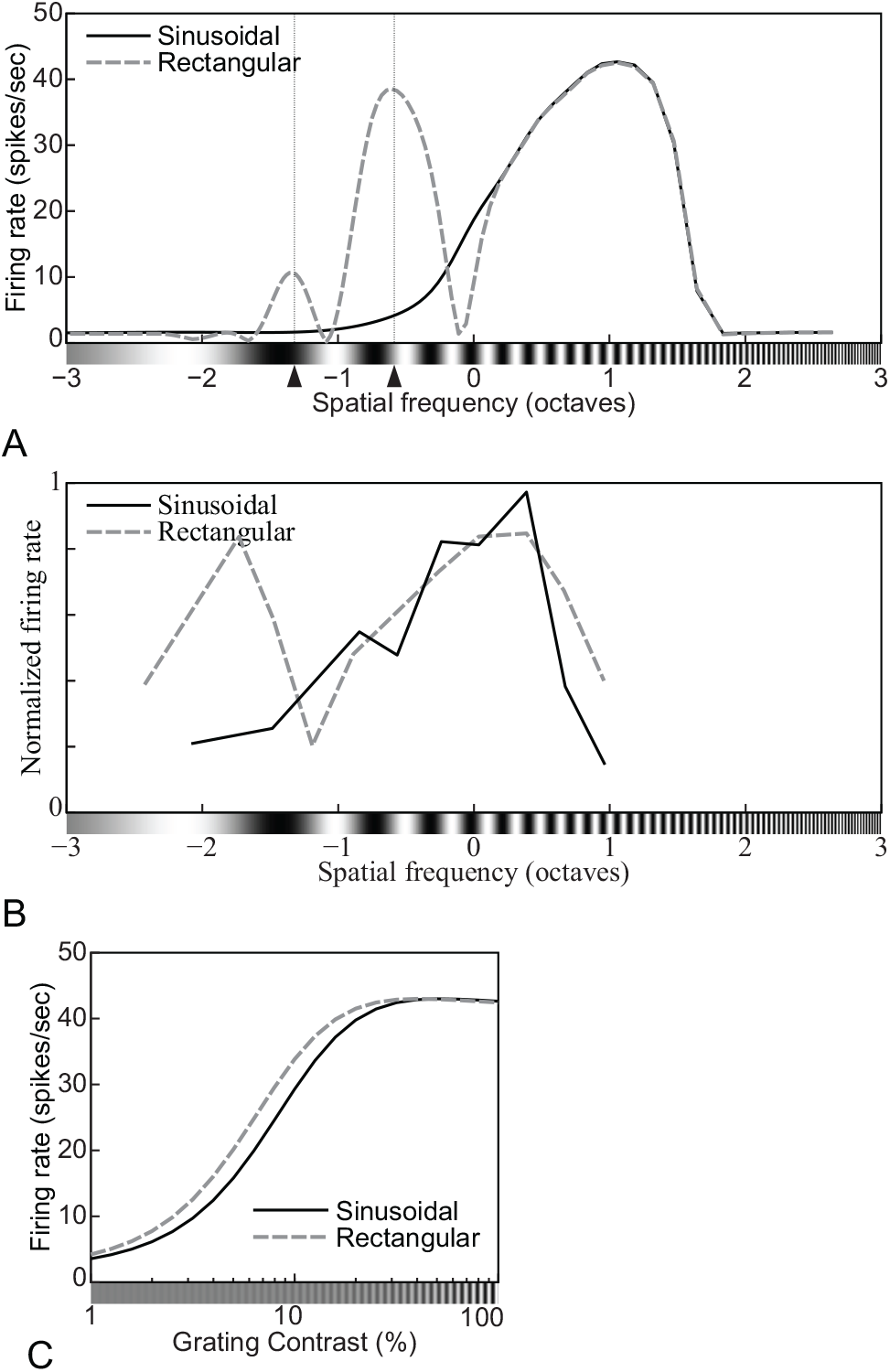
Spatial-frequency tuning functions for sinusoidal and square gratings: (A) of the model neuron with a modified parameter set (*h*_*f*_ = 0.8, *h*_*F*_ = 0.4) and (B) of a complex cell (replotted from Pollen & Ronner, 1982, Fig. 6E, anesthetized cat). Note the secondary peak at one-third of the tuned frequency for square gratings (dashed line). (C) The contrast sensitivity functions of the model neuron with the modified parameter set for sinusoidal and square gratings (indicated in legend). See Phenomena 19 and 20 in Table 1.

#### 3.3.4 Cross-orientation suppression

The responses of real neurons in V1 to a grating (signal) tend to be suppressed by another grating (mask) superimposed onto the signal grating within the neuron’s measured RF. This *cross-orientation suppression* occurs in wider ranges of the orientation and spatial-frequency domains than the excitatory tuning of the neuron (DeAngelis et al., 1992; see also K. K. De Valois & Tootell, 1983). The strongest suppression occurs when the mask orientation is the same as the tuned orientation (Phenomenon 21, Morrone et al., 1982; DeAngelis et al., 1992, see also Bonds, 1989 for a mask-induced facilitation effect). These trends were captured by the DNM neuron (Figure 16A). As for spatial frequency, the most suppressive frequency of the mask grating does not necessarily coincide with the tuned frequency of the real neuron (Phenomenon 22, K. K. De Valois & Tootell, 1983; Morrone et al., 1982; DeAngelis et al., 1992). The most suppressive frequency of the DNM neuron (Figure 16C) seems to be lower than its tuned frequency (1.0 oct). One possible explanation of the latter result is that the diameter of the measured RF could have been too small for a grating with a spatial-frequency lower than the tuned frequency.

**Figure 16:**
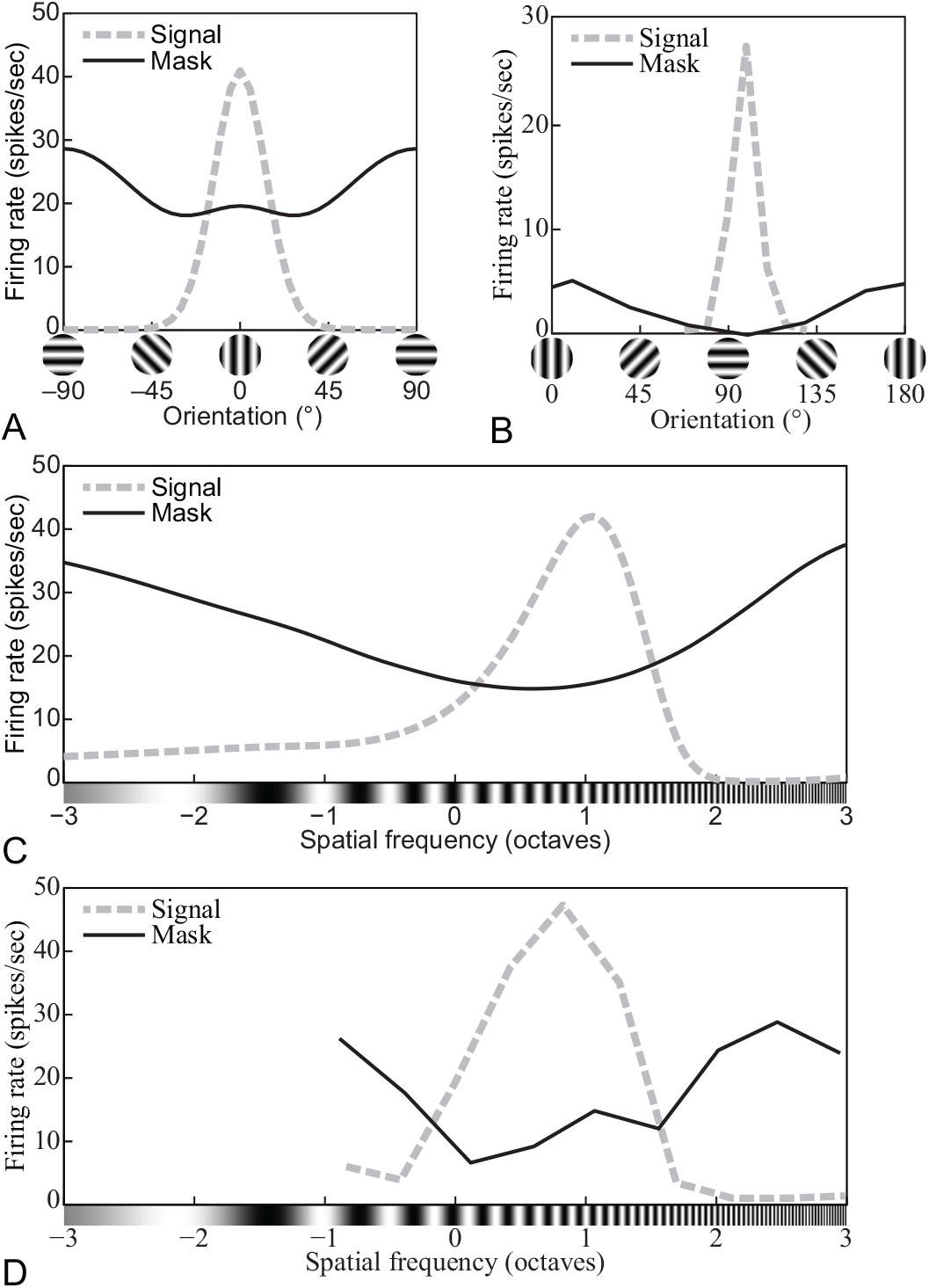
The effect of *cross-orientation suppression* on: (A, C) the orientation and frequency tuning functions of the model neuron with standard parameters, (B) the orientation tuning function of a complex cell (replotted from DeAngelis et al., 1992, Fig. 7CD, anesthetized cat), and (D) the frequency tuning function of a simple cell (replotted from DeAngelis et al., 1992, Fig. 3AB, anesthetized cat). In all panels, the standard tuning function (as probed with a non-masked grating) is plotted by the dashed gray line. The tuning function of orthogonal suppression is plotted by a solid black line. (A) The contrast and frequency of the signal grating were 20% and 1.0 oct. The contrast and frequency of the mask grating were 40% and 0.0 oct. The size of the stimuli was 2.88°. (C) The contrast and orientation of the signal grating were 20% and 0°. The contrast and orientation of the mask grating were 80% and 90°. The size of the stimuli was 2.88°. See Phenomena 21 and 22 in Table 1.

The mask grating affects the contrast response function (CF) of real neurons (Phenomenon 23, Carandini, 2004; Freeman et al., 2002; Morrone et al., 1982). Panel B of Figure 17 shows representative physiological data (Freeman et al., 2002) and panel A shows simulation results of a DNM neuron. The CF of the DNM neuron shifts horizontally right if the contrast of the mask is high enough (≥ 12%). The suppression effect is negligible if the mask contrast is lower (< 12%). This trend agrees with the physiological results. The responses are also plotted as functions of the mask contrast in panels C and D. In both the model and the data, the response functions shift upward and scale up vertically as the contrast of the signal grating increases.

**Figure 17:**
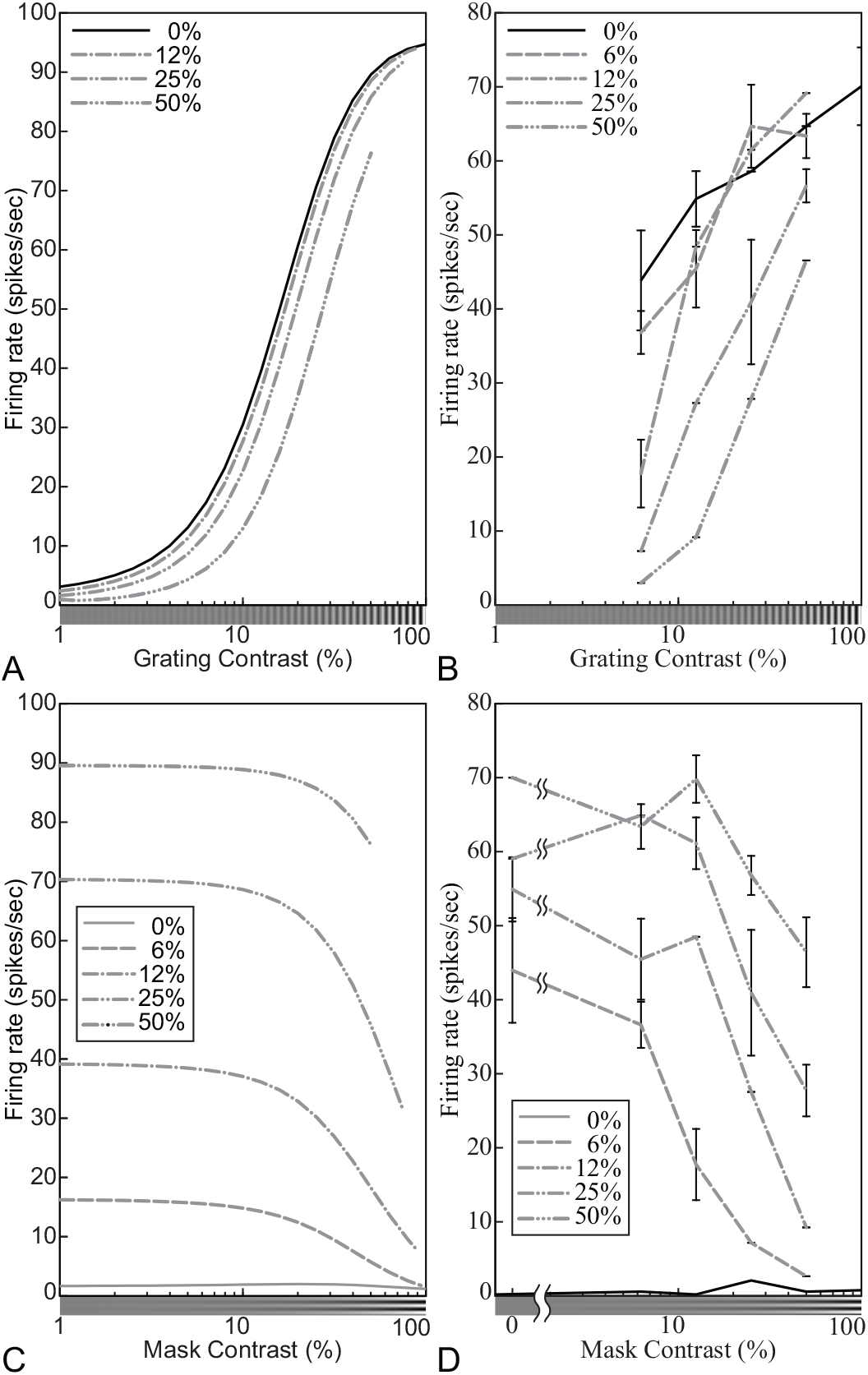
The effect of *cross-orientation suppression* on the contrast response functions of (A, C) the model neuron with standard parameters and (B, D) a V1 neuron (replotted from Freeman et al., 2002, Fig. 6, anesthetized cat). In (A, B), the responses of the model and real neurons are plotted against the contrast of the signal grating, with mask contrast shown in the legend. In (C, D), the responses are plotted against the contrast of the mask grating, with signal contrast in the legend. (A, C) The size of the stimuli was equal to the measured RF diameter (0.81°). The orientations of the signal and mask gratings were 0° and 90°, and their spatial-frequency was 2.0 cpd. See Phenomenon 23 in Table 1.

#### 3.3.5 Surround suppression

The responses of real neurons to a grating patch within the neurons’s measured RF tend to be suppressed by an annular grating surrounding the RF. This *surround suppression* is strongest when the orientation and frequency of the annular grating are the same as those the neuron is tuned to (Phenomena 24 and 25, Nelson & Frost, 1978; C.-Y. Li & Li, 1994; Blakemore & Tobin, 1972; DeAngelis et al., 1994; Ozeki, Finn, Schaffer, Miller, & Ferster, 2009; Ozeki et al., 2004; Cavanaugh et al., 2002b; Sillito, Grieve, Jones, Cudeiro, & Davis, 1995). The ranges of the surround suppression in both domains are wider than the respective tuned bandwidths. These trends were captured by the DNM neuron with standard parameters (Figure 18).

**Figure 18:**
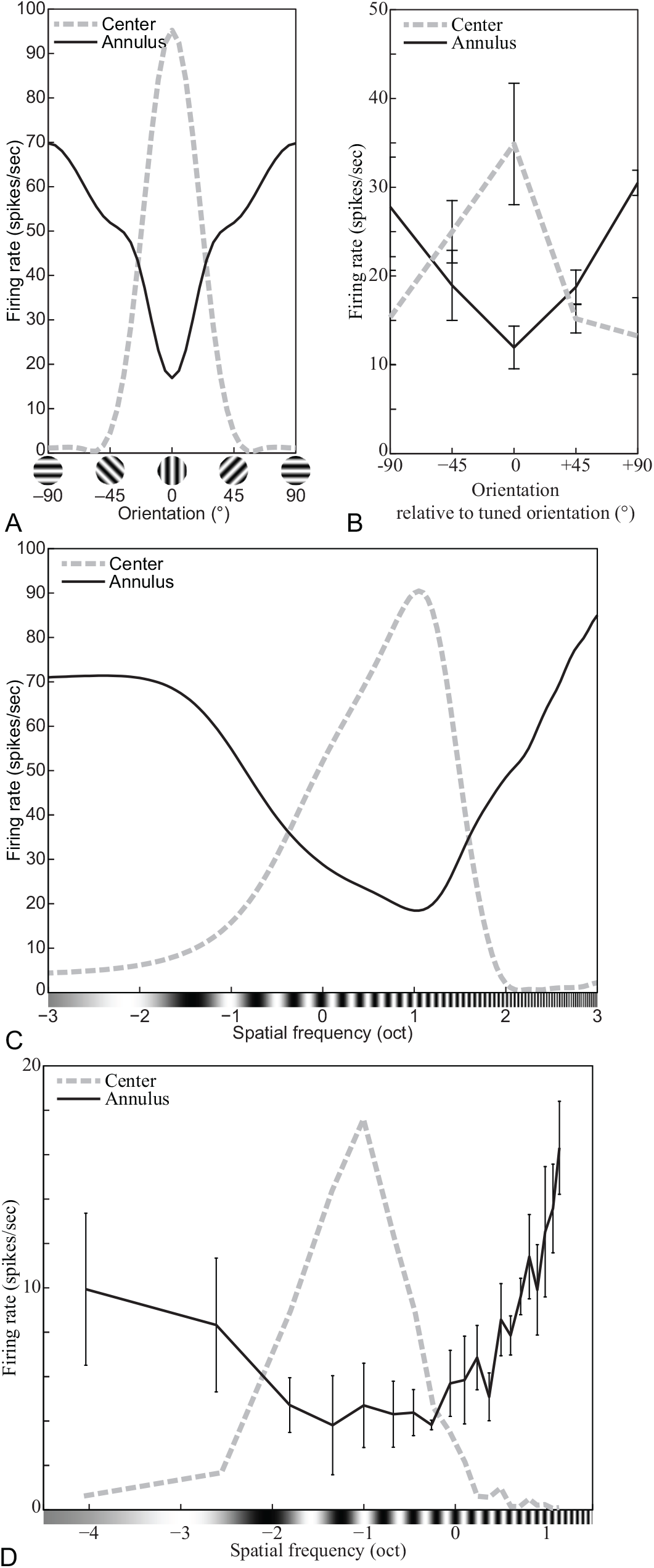
The effect of surround suppression on the response of (A, B) the model neuron with standard parameters, (C) a V1 complex cell (replotted from Cavanaugh et al., 2002b, Fig. 2B, anesthetized macaque, error bars = ±SE), and (D) a V1 simple cell (replotted from C.-Y. Li & Li, 1994, Fig. 7C, anesthetized cat, error bars = ±SE) in the orientation and the spatial-frequency domains. In all panels, the standard tuning function (as probed with no surrounding annulus) is plotted by the dashed grey line. The tuning function of the surround suppression is plotted by a solid black line. (A, C) The contrast of the center grating was 100% and its diameter was 0.81° (the measured RF). The contrast of the annulus grating were 100% and 5.76°. The frequency of the center and annular gratings was 2.0 cpd for (A). Their orientation was 0° for (C). See Phenomena 24 and 25 in Table 1.

The contrast response function of real neurons is affected by the contrast of an annular grating surrounding the classical RF (Phenomenon 26, Cavanaugh et al., 2002a; Carandini, 2004; DeAngelis et al., 1994). Panel B of Figure 19 shows the CF of a complex cell in cat V1 with different contrasts of the annular grating (Carandini, 2004). Panel A shows the corresponding plot for a DNM neuron with standard parameters. Both empirical and simulated CFs shift horizontally to the right as the contrast of the annular grating increases. Note that the slope of the CF of the model neuron is not necessarily invariant against the annular grating. With a different parameter set (*n*_*n*_ < *n*_*d*_), the CF of the DNM neuron became shallower as the contrast of the annular grating increased (Figure 19C). The same trend has been observed for some V1 neurons (Figure 19D, Carandini, 2004).

**Figure 19:**
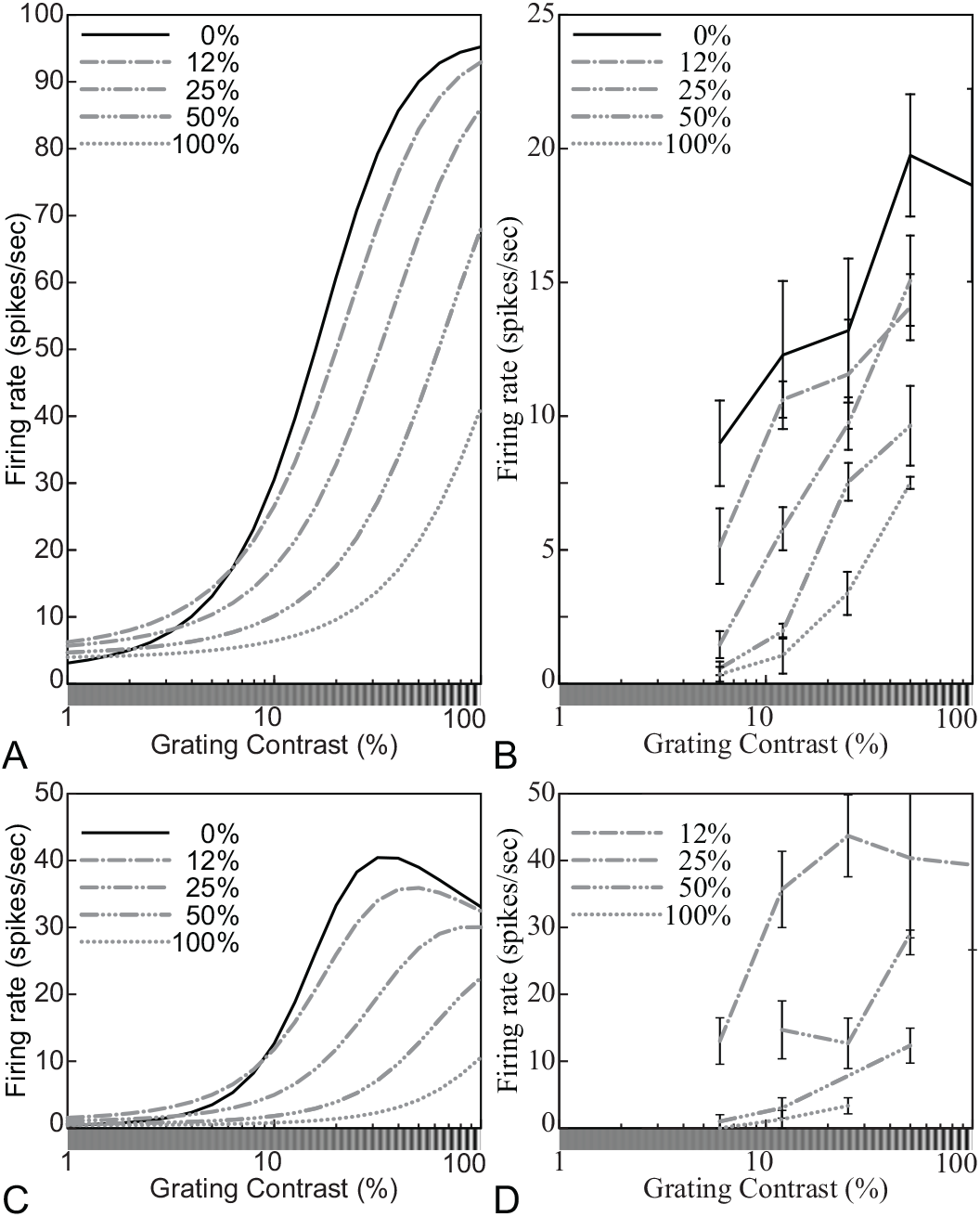
Surround-suppression effects on the contrast response functions for different contrasts of the annular grating. The contrast of the central disk is plotted on the horizontal axes, and the contrast of the annular surround is indicated in the legend. (A) The model neuron with the standard parameter set (cf. Table 2); (B) a complex cell in cat V1 (replotted from Carandini, 2004, Fig. 6); (C) the model neuron with a modified parameter set (*n*_*n*_ = 2.8, *n*_*d*_ = 3.0, *M* = 10); and (D) a complex cell in cat V1 (replotted from Carandini, 2004, Fig. 7). (A, C) The orientation and spatial-frequency of the center and annulus gratings were 0° and 2.0 cpd. The diameter of the center grating patch was 0.81° and it is equal to the measured RFs of the model neuron for both parameter sets. The size of the annulus grating was 5.76°. See Phenomenon 26 in Table 1.

The effect of the annular grating also depends on its orientation for both real neurons (Figure 20B, Phenomenon 27, Cavanaugh et al., 2002b) and the model neuron (Figure 20A). Note that the cross-orientation suppression and the surround suppression of real neurons show analogous trends to one another, so they can be modeled jointly by the suppressive drive in Equation 13. Note that this similarity at the functional level does not necessarily imply similarity at the mechanistic level. In fact, physiological results indicate that the two types of suppression should be attributed to different mechanisms (see DeAngelis et al., 1992, 1994, for examples). We will discuss the difference between these two types of suppression in Section 4.3 below.

**Figure 20:**
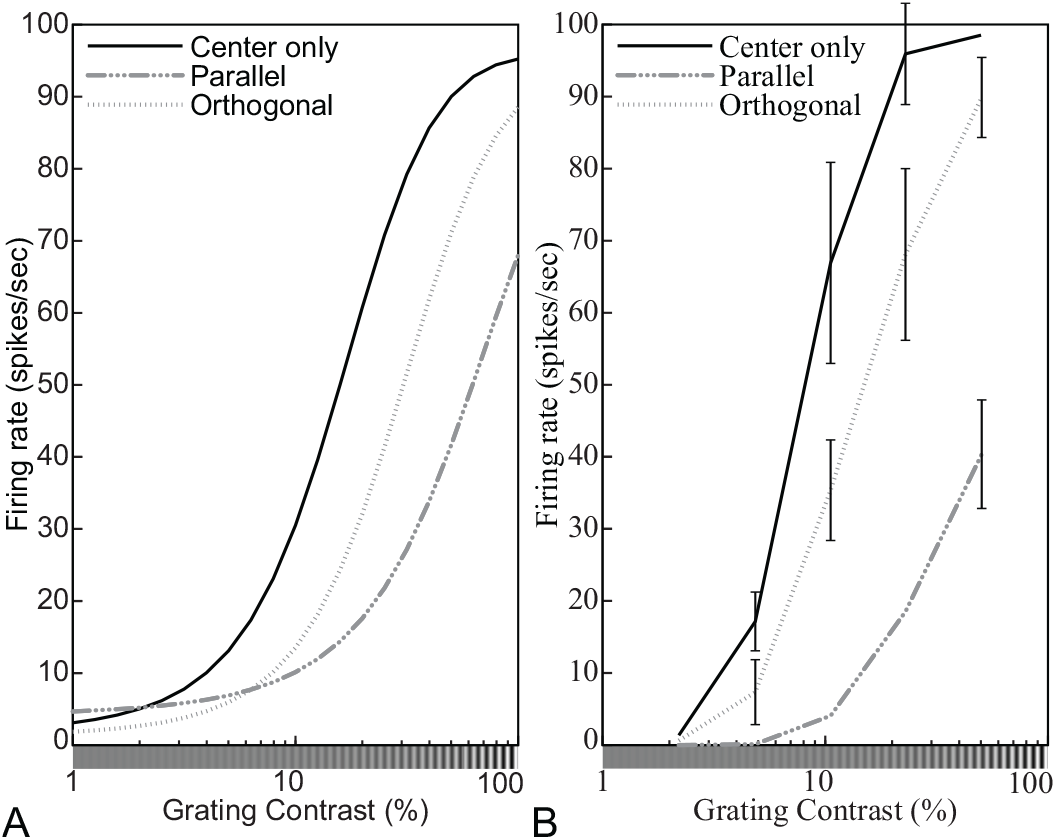
Surround-suppression effects on the contrast response functions for different orientations of the annular grating (indicated in the legend). (A) The DNM neuron with standard parameters and (B) a simple cell (replotted from Cavanaugh et al., 2002b, Fig. 5A, anesthetized macaque, error bars = ±SE). (A) The orientation of the center grating was 0°. The spatial-frequency of the center and annulus gratings was 2.0 cpd. The size of the center grating patch was equal to the measured RF diameter (0.81°). The size of the annulus grating was 5.76°. (DNM = divisive normalization model. See Phenomenon 27 in Table 1.)

#### 3.3.6 Receptive fields of simple cells

The RF of a real simple cell is composed of excitatory and inhibitory sub-regions. This composition has been mapped using local stimulus probes such as a light spot (Hubel & Wiesel, 1959; Volgushev, Vidyasagar, & Pei, 1996), a light bar (Andrews & Pollen, 1979), light and dark bars (Movshon et al., 1978b; Tadmor & Tolhurst, 1989; Kulikowski & Bishop, 1981; Kulikowski & Vidyasagar, 1986; Glezer, Tsherbach, Gauselman, & Bondarko, 1982), and the *reverse correlation* method (Nishimoto, Ishida, & Ohzawa, 2006; Ringach, 2002; Moore IV & Freeman, 2012; see also Gardner, Anzai, Ohzawa, & Freeman, 1999b). The classical RF of the DNM simple cell was also mapped using these methods (Figures 21 and 22, Phenomena 28 and 29). Note that a firing rate of a real simple cell becomes lower than its maintained discharge 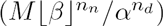 when the light spot stimulates its inhibitory sub-regions (Hubel & Wiesel, 1959, Figure 21) or the light and dark bars stimulate its inhibitory and excitatory sub-regions respectively (Andrews & Pollen, 1979, Figure 23C). This effect can be observed with the DNM simple cell only when its maintained discharge in the absence of external stimulation is high enough to reveal the inhibitory effect of a light-spot probe. This occurs when 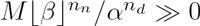.

**Figure 21:**
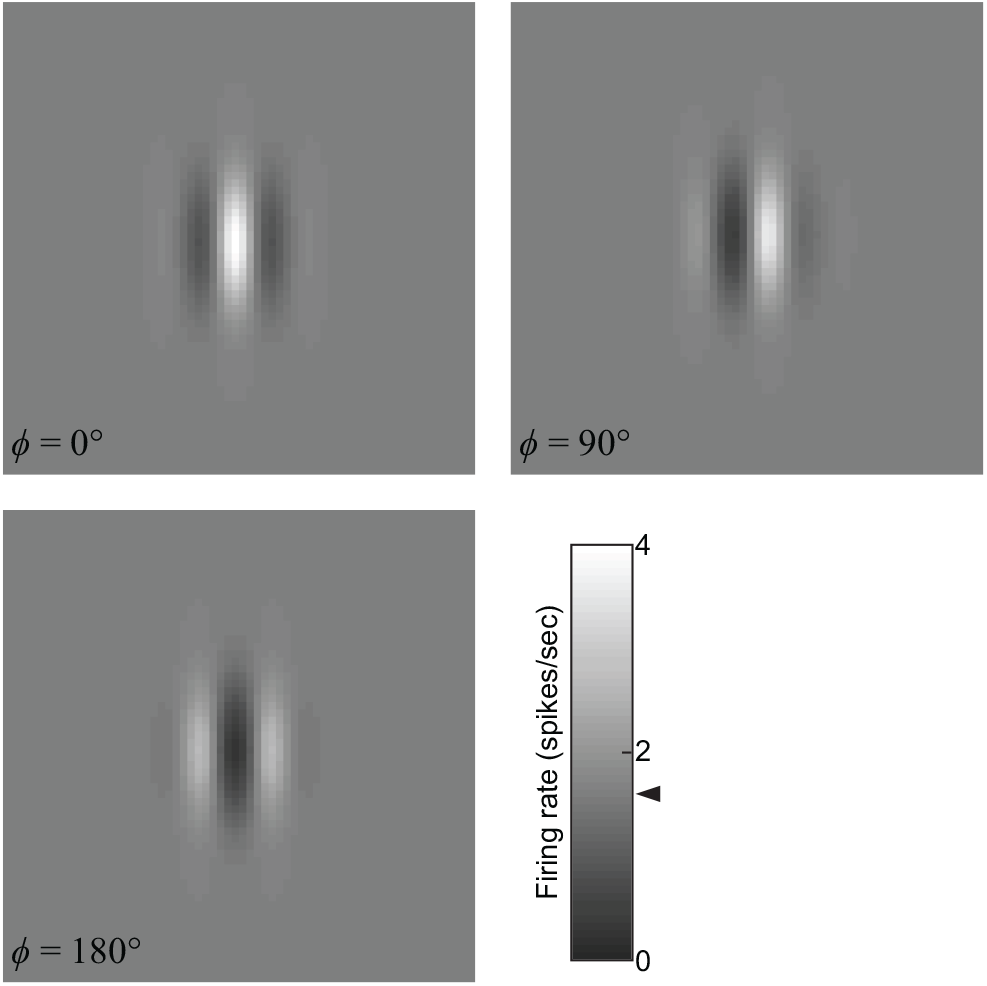
Receptive-field maps obtained with a light spot as a probe for three DNM simple cells with different phases *ϕ*. The grayscale levels indicate firing rates. The maintained discharge (the response to a uniform gray field) was the same for the three cells and is indicated by the triangle on the color bar. The size of the light-spot probe was 0.045° × 0.045° (1 pixel). The stimulus background was uniform gray and the luminance of the spot was twice as high as the background gray. (DNM = divisive normalization model, standard parameters. See Phenomenon 28 in Table 1.)

**Figure 22:**
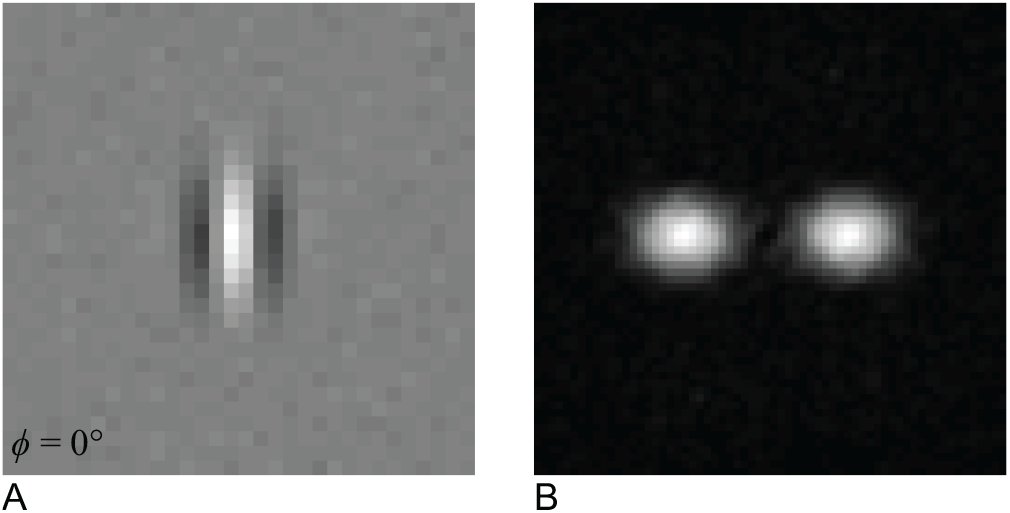
(A) The receptive field of a DNM simple cell obtained via the reverse correlation method. Brighter and darker regions represent excitatory and inhibitory regions, respectively. (B) The 2D Fourier spectrum energy distribution of the RF pattern in (A). The stimuli were 32 × 32 mosaics of random luminance spots (white noise). The size of the stimuli was 2.88° and that of the spots was 0.090° (2 × 2 pixels). (DNM = divisive normalization model, standard parameters. See Phenomenon 29 in Table 1.)

Consider a hypothetical simple cell that acts as a purely linear filter of the visual stimulus. Under this linearity assumption (cf. Section 2.1), the bandwidths in the orientation and spatial-frequency domains can be derived from the 2D composition of its excitatory and inhibitory sub-regions in its RF in the image domain (Eq. 3 and 4; Graham, 1989; Lathi, 2005). However, the derived bandwidths of real simple cells tend to be wider than these measured directly with gratings of various orientations and frequencies (Tadmor & Tolhurst, 1989; Gardner et al., 1999b; Nishimoto et al., 2006; Ringach, 2002; see also Section 3.3.3). These results suggest that real simple cells are non-linear. This discrepancy of the bandwidths is also observed with the model simple cell (Figures 22 and 23B). The derived bandwidths in the orientation and the frequency domains were 42.6° and 1.53 oct using the reverse correlation method and were wider than those measured with a grating (31.8° and 1.11 oct). This trend could be observed from the DNM cell if *n_n_ >* 1 (see Gardner et al., 1999a). Note that the derived bandwidths were rather close to the specified bandwidths of the stimulus drive (40° and 1.5 oct). Several other parameter sets (including *n*_*n*_ < 1) were tested and this similarity was observed reliably. Note that the stimulus probe itself could also affect the results. Recall that the RF could be measured with a bar as the probe. However, as the bars become wider, the derived bandwidth became wider and the tuned frequency derived was lower (Figure 23C). This underestimation of the tuned frequency was also observed in the physiological data (Tadmor & Tolhurst, 1989).

**Figure 23:**
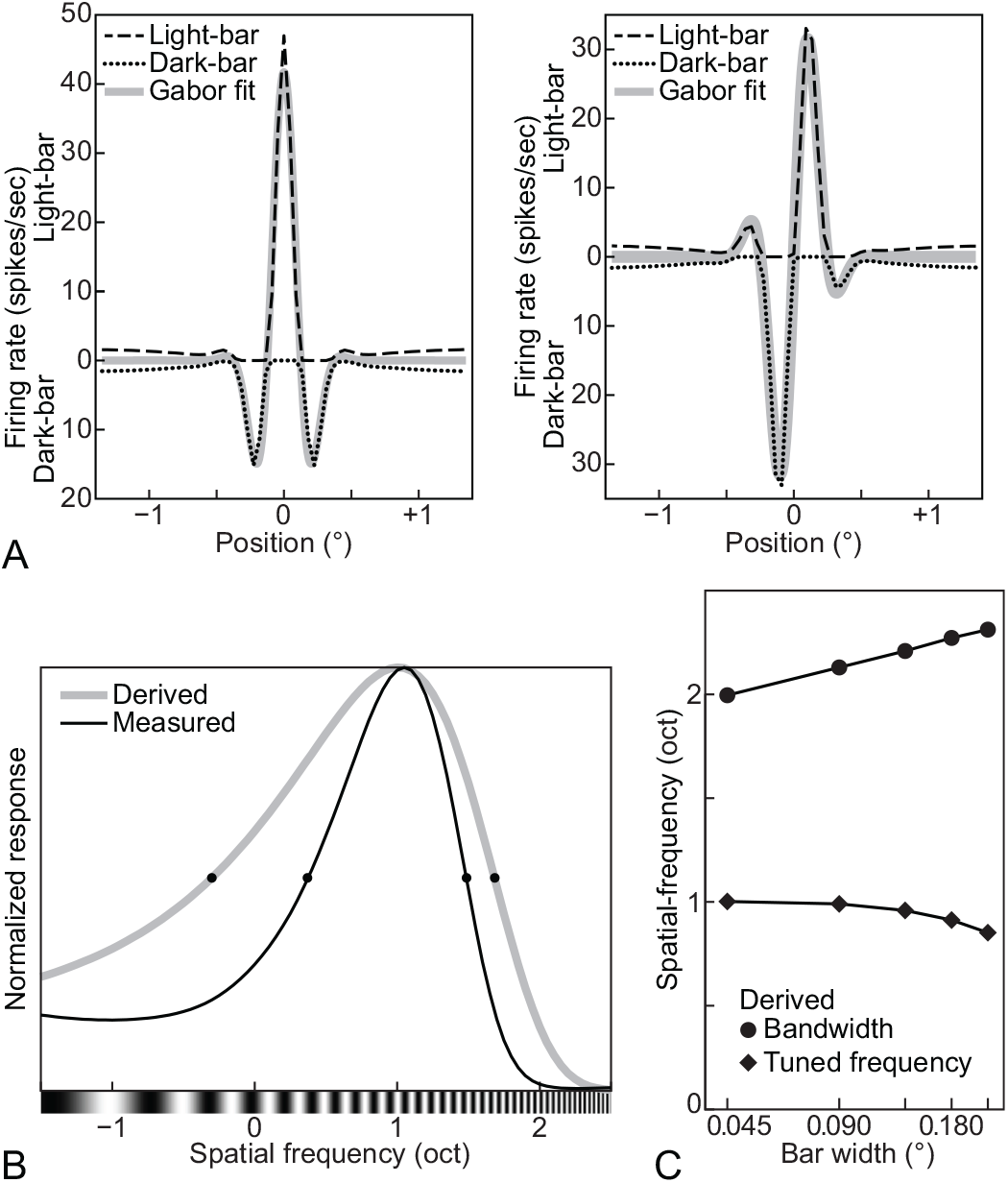
(A) A 1D distribution of the responses to light and dark bars parallel to the tuned orientation of the DNM simple cell. The abscissa of the graph represents the positions of the bars along an orientation perpendicular to the tuned orientation. The upper half of the graph is for the light bar and the lower half of the graph is for the dark bar. The best-fitting 1D Gabor pattern is superimposed (in gray) for comparison. The background of the stimulus was uniform gray and the luminance of the bars was 0 (dark) or twice as high as the background gray (light). The width of the bars was 0.045° (1 pixel). (B) The spatial-frequency tuning functions of the DNM simple cell derived (gray line) from the 1D pattern of its receptive field in (A) and measured (black line) using a grating. (C) The effect of the width of the bars on the tuned spatial-frequency derived from the measured RF as in (A). (DNM = divisive normalization model, standard parameters. See Phenomenon 30 in Table 1.)

### 3.4 Summary and discussion

The results of our simulation experiments show that the model neuron based on the divisive normalization Equation 13 can account for many physiological phenomena (Table 1) with a standard parameter set (Table 2). A few other phenomena can also be accounted for, but they require customized parameter sets. Based on these simulation results and on mathematical analyses, we can make some falsifiable predictions that can be used to test the divisive normalization model (DNM).

All predictions involve probing a single neuron with multiple stimuli. The theoretical constraints stem from the fact that the model parameters must be fixed for each individual neuron. Thus, certain patterns are expected to occur together in the responses of a given neuron because they all depend on a single model parameter. In particular, the baseline parameter *β* in Equation 13 gives rise to interesting constraints. We have shown that the following three phenomena can be produced by the model only when *β* is sufficiently large:

(A) The inhibitory sub-regions of a simple cell probed with a single light spot (Hubel & Wiesel, 1959) can be observed only when 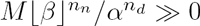. This condition ensures that the DNM simple cell has a substantial maintained discharge in the absence of external stimulation. This is necessary to reveal the inhibitory effect of a light-spot probe (Phenomenon 28, Figure 21).

(B) The supersaturation effect (Ph. 8, Fig. 10) can occur in the model only when 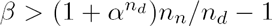 (Eq. 21). Recall from Section 3.3.2 that this effect refers to the non-monotonicity of the contrast sensitivity function of some V1 neurons. (Note that the majority of V1 neurons have monotonically increasing CFs.)

(C) Finally, the widening of the bandwidths in the orientation and in the spatial-frequency domains with decreasing stimulus contrast (Ph. 18, Fig. 14D, H, see also Ph. 17) can occur in the model only when *β >* 0.

Because of their dependence on a common parameter, the divisive normalization model predicts a correlation among these thee phenomena if they are tested for each individual cell in a sample of V1 neurons recorded in a physiological experiment.

These interlocking patterns are methodologically important because they show that the divisive normalization model is constrained enough to be falsifiable even though it has 10 free parameters that give it the flexibility it needs to account for the diversity of V1 neurons. This novel theoretical result emerges from the examination of a comprehensive suite of phenomena within the framework of a single model with consistent parameters.

The model also provided alternative interpretations of a few phenomena observed in physiological studies. First, the tuning bandwidths of real simple cells are often narrower when measured directly in the orientation and the spatial-frequency domains compared to those derived from the mapped spatial patterns of the receptive fields (see Fig. 23). This departure from linearity has often been explained by the so-called *iceberg effect,* which occurs in our model when *β* < 0 (Carandini & Ferster, 2000; Sompolinsky & Shapley, 1997; Volgushev, Pernberg, & Eysel, 2000). The present simulations (Fig. 23) show that the model can produce such departures from linearity even when *β* > 0. Across its parameter range, the model produces departures from linearity due to the non-linear character of the numerator of the divisive-normalization equation (Gardner et al., 1999a).

Another phenomenon is the effect of stimulus size on the orientation- and frequency-bandwidths of real V1 neurons (see Fig. 13). The measured bandwidths depend on the size of the grating patch, and the size that maximizes the bandwidths is larger than the measured RF of the neuron. This occurs because RF mapping methods tend to underestimate the true extent of the weighting function (WF) of the linear filtering stage of the model neuron. The periphery of the WF of the model neuron is hidden by an annular region of the surround suppression, which narrows down the measured RF of the model neuron. This means that the stimulus patch must be larger than the measured RF to fill the true WF. Note that our simulations show that the surround suppression itself actually makes the bandwidths of the DNM neuron wider (see Section 3.3.3).

A few studies have shown that the size of the grating patch also bears some relationship to the preferred frequency of real neurons. The peak of the frequency tuning curve of real neurons tends to shift to a higher frequency as the size of the grating patch decreases (Teichert et al., 2007; Osaki et al., 2011). Also, the peak of the size tuning curve of the real neuron tends to shift to a smaller size if the spatial-frequency of the grating is higher than the tuned frequency of the neuron (Osaki et al., 2011). These trends were not observed with the DNM neuron used in the simulation experiments. Such trends, however, can be emulated by a model with some additional flexibility. For example, the suppressive drive of the DNM model can be modified so that the suppression in the spatial-frequency domain changes depending on the eccentricity from the center of the RF. These trends can also be expected to be emulated by the modified DNM if the peak of the suppression in the spatial-frequency domain shifts to a higher frequency as eccentricity increases. Another possible modification involves the stimulus drive. Note that Naito and Sato (2015, August) proposed that if you want to model these trends, your model neurons should be composed of multiple Gabor filters. These filters can be tuned to different spatial-frequencies and the sizes of their WFs can be correlated with the tuned frequencies, i.e., smaller WFs are tuned to higher frequencies. Also, following the suggestions of Naito and Sato (2015, August), analogous modifications can be made to the DNM, composing its stimulus drive with multiple Gabor filters.

## 4 General Discussion

Our simulation experiments demonstrated that the divisive normalization model (DNM) can account for a comprehensive set of neurophysiological studies of both simple and complex cells in V1 (Table 1). Moreover, a mathematical analysis of Equation 13 predicts interdependence between certain observable phenomena. In the prior section, we explained how these predictions can be tested experimentally, which means that the standard formulation of the DNM specified in Section 2.3 is a falsifiable theory. If this formulation, especially Equation 13, is modified, the concrete mathematical results proven in the appendices will no longer apply. It seems plausible, however, that analogous results would hold for any formulation based on similar principles. In other words, our theoretical conclusions probably extend to the entire class of models based on a combination of linear filtering, half-wave rectification and squaring, and response normalization (Carandini et al., 2005). Our results ultimately rest on the fact that, although adjustable parameters are needed to accommodate the diversity across neurons, they must be fixed for each individual neuron. This fixedness gives rise to falsifiable constraints when a single neuron is probed with a judiciously chosen battery of stimuli.

With the standard parameter set in Table 2, the DNM qualitatively replicates most of the empirical phenomena listed in Table 1. Quantitative comparisons of the DNM with real neurons are also possible but, they would require quantitative estimation of all parameters of the DNM for the individual neurons from physiological data. The interpretability of such quantitative tests would depend on the precision of the estimation of the parameters. Finding the parameters that best fit the physiological data often involves the numerical optimization of an objective function. Depending on the data, it is possible that this function has multiple local minima that correspond to alternative parameter settings. When the parameter search procedure is trapped in a local optimum, it becomes difficult to compare the model predictions to empirical observations because different local minima would lead to different predictions. Such instability of the parameter estimation would jeopardize the status of the DNM even as a descriptive model unless the parameter search is guaranteed to find the global optimum.

### 4.1 Correlation between the parameters of the model

The parameters of the model are conceptually independent from one another but physiological studies have shown correlations among some of them. These correlations can potentially decrease the number of independently adjustable parameters or limit the ranges of these parameters. This subsection briefly reviews some studies that suggest correlations among various DNM parameters.

The preferred spatial-frequency *F\** of the divisive normalization model correlates with some parameters of the contrast response function (CF). Dean (1981) reported that the preferred spatial-frequencies of neurons in cat striate cortex correlates positively with their contrast threshold and negatively with the slope of their CF. Consider the hyperbolic-ratio model for simplicity (Eq. 10). Its mathematical analysis shows that the hyperbolic-ratio model can account for both correlation patterns above if the semisaturation contrast parameter *α*_*HB*_ correlates positively with the preferred frequency *F\** across the population of model neurons (see Section 2.2). Some studies have shown that the bandwidths of V1 neurons tend to be narrower in neurons with higher preferred spatial frequencies (Kulikowski & Bishop, 1981; Kulikowski & Vidyasagar, 1986; R. L. De Valois, Albrecht, & Thorell, 1982; Yu et al., 2010). This effect can be emulated by the divisive normalization model if either *β*, *n_d_,* or *h*_*f*_ decrease or *α, n_n_,* or *h*_*F*_ increase as the tuned frequency *F\** increases. (See Table 2 for the meaning of these symbols.) Note that the bandwidths in the spatial-frequency and orientation domains correlate with one another (Zhu, Xing, Shelley, & Shapley, 2010). Both of those bandwidths would be affected together if *α, β*, *n*_*n*_, or *nd* covary with *F*.*

The receptive-field locations of neurons affect their orientation- and frequency tuning. The DNM can emulate this by introducing a correlation between the RF location parameters *(x, y)* and the tuning preference parameters Θ* and *F*.* First, as the retinal eccentricity of the positions increases, the tuned frequencies of real neurons decrease (Movshon et al., 1978b; Yu et al., 2010; Henriksson, Nurminen, Hyvärinen, & Vanni, 2008) and this correlation can be well represented well by the cortical magnification factor (Swindale, 1996; Daniel & Whitteridge, 1961; Tootell, Silverman, Switkes, & De Valois, 1982; Schwartz, 1980; Duncan & Boynton, 2003). Next, the observed distribution of tuned orientations has a preponderance of vertical or horizontal preferences for neurons with RFs in or near the fovea (R. L. De Valois, Yund, & Hepler, 1982; G. H. Henry, Dreher, & Bishop, 1974) and a preponderance of radial orientations for neurons in the visual periphery (Schall, Vitek, & Leventhal, 1986; see also Y. Sasaki et al., 2006, for fMRI results). The results of an fMRI study suggest that human V1 contains more vertically tuned neurons than horizontally tuned neurons (Yacoub, Harel, & Uğurbil, 2008). On the other hand, the number of the vertical and horizontal neurons found in cat striate cortex are almost equal (B. Li, Peterson, & Freeman, 2003), whereas rat V1 apparently has a preponderance of neurons with horizontal preferences (Girman, Sauvé, & Lund, 1999). Note that the neurons tuned to vertical or horizontal orientations, and to high spatial-frequencies, have narrower bandwidths in the orientation domain when compared to neurons tuned to oblique orientations (Orban & Kennedy, 1981; B. Li et al., 2003).

### 4.2 The stimulus drive

The current formulation of the divisive normalization model specifies the weighting function of simple cells as a 2D Gabor filter (Eq. 5). There are, however, some other filters that can fit physiological data better than the Gabor filter does (see Stork & Wilson, 1990; Wallis, 2001, for reviews). All those models of the simple cell use a linear filtering stage (cf. Eq. 5) as their first processing step. This linear filtering predicts that the cell’s response will be maximal for a square wave grating with their tuned frequency, orientation, and phase.

Simple and complex cells are represented in qualitatively different ways in the current formulation of the DNM (see Section 2.1). Note, however, that it has been shown that these two types of neurons are better conceptualized as endpoints of a continuum rather than as a categorical distinction (Mechler & Ringach, 2002; Priebe, Mechler, Carandini, & Ferster, 2004; Martinez & Alonso, 2003; Chance, Nelson, & Abbott, 1999, 2000). Note that the original classification was based on 2D patterns of the classical RFs (Hubel & Wiesel, 1959, 1962, 1968; Bishop & Henry, 1972). Later proposals classified the neurons on the basis of the temporal modulation of their firing rates evoked by a drifting grating (e.g., Maffei & Fiorentini, 1973; Movshon et al., 1978c, 1978a; Andrews & Pollen, 1979). Simple cells tend to show greater modulation than complex cells. The classifications based on these two methods often agree with one another (R. L. De Valois, Albrecht, & Thorell, 1982; Skottun et al., 1991; Dean & Tolhurst, 1983; Mata & Ringach, 2005; Sengpiel et al., 1997; C. A. Henry & Hawken, 2013) but the agreement is not always complete (see Chen et al., 2009, for a review). There are always some neurons whose behaviors are intermediate between the “pure” types defined by either method (Van Kleef, Cloherty, & Ibbotson, 2010; Crowder, van Kleef, Dreher, & Ibbotson, 2007; Hietanen et al., 2013; Meffin, Hietanen, Cloherty, & Ibbotson, 2015; Mata & Ringach, 2005; Kagan, Gur, & Snodderly, 2002). These intermediate neurons would be difficult to emulate by models that make a categorical distinction between simple and complex cells. Also, the temporal modulation of the firing rates can be affected by other properties of the visual stimuli. For example, some complex cells show temporal modulation if the contrast of the drifting grating is low (Crowder et al., 2007; Van Kleef et al., 2010, see also C. A. Henry & Hawken, 2013). The temporal phase of the modulation also changes as a function of stimulus contrast (Albrecht, 1995). The temporal modulation of firing rate is also affected by stimulation of the surround regions of the neuron’s classical RF (Bardy, Huang, Wang, FitzGibbon, & Dreher, 2006). The effect of the surround stimulation on the modulation is either facilitatory or suppressive depending on the individual neurons.

Neurons in V1 are tuned to gratings with particular orientations in static images and many of the neurons are selective to directions of drifting gratings (e.g., Livingstone, 1998; H. E. Jones et al., 2001). This *direction selectivity* could be modeled by a drifting 2D Gabor filter, that is, by a 2D Gabor filter whose phase changes over time (e.g., Heeger, 1993). This formulation of the direction selectivity predicts that the model responses would remain constant for a drifting grating whose orientation, spatial frequency, direction, and speed match the corresponding tuning preferences of the model neuron. Heeger (1993) showed that some other temporal properties of V1 neurons for drifting gratings can also be also emulated by integrating the drifting 2D Gabor filter and using it as the stimulus drive into the divisive normalization model. It is possible that many temporal properties of the neuronal responses to stimuli within their RFs can be well represented well by a relatively simple model that is based on divisive-normalization principles (see Albrecht et al., 2002, for an example).

### 4.3 The suppressive drive

The standard DNM formulation includes both surround- and cross-orientation suppression in a single *suppressive drive* (Eq. 16) for simplicity. Despite the fact that these two forms of suppression show some analogous properties, there is evidence that they should be attributed to different neurophysiological mechanisms (Sengpiel et al., 1998; Angelucci & Bullier, 2003). It has been suggested that cross-orientation suppression consists of a binocular factor arising from intracortical connections within V1 (B. Li, Peterson, Thompson, Duong, & Freeman, 2005; Endo, Kaas, Jain, Smith III, & Chino, 2000; Sengpiel & Vorobyov, 2005) and a monocular factor from thalamo-cortical connections between LGN and V1 (B. Li, Thompson, Duong, Peterson, & Freeman, 2006; DeAngelis et al., 1992; Priebe & Ferster, 2006; Smith, 2006; Freeman et al., 2002), but not from neurons in the LGN (Bauman & Bonds, 1991). It has also been suggested that surround suppression can come from LGN inputs to V1 (Webb, Dhruv, Solomon, Tailby, & Lennie, 2005), and from intracortical connections within V1 itself (Ozeki et al., 2009), and from feedback from higher cortical areas (Angelucci & Bullier, 2003; Nassi, Lomber, & Born, 2013; Bair et al., 2003; W. Li, Thier, & Wehrhahn, 2001; see also Angelucci & Bullier, 2003 for a review). In addition, the responses of some neurons are *facilitated* in a non-linear manner by stimuli outside their classical RFs (Fitzpatrick, 2000; C.-Y. Li & Li, 1994; Maffei & Fiorentini, 1976; Sillito et al., 1995; Cavanaugh et al., 2002b; Polat, Mizobe, Pettet, Kasamatsu, & Norcia, 1998; H. E. Jones et al., 2001; Nelson & Frost, 1985), as well as by stimuli within the classical RFs (K. K. De Valois & Tootell, 1983). The facilitatory stimuli are different depending on the idividual neurons. A judicious combination of surround suppression and surround facilitation can tune these neurons to more complex visual features (Grossberg & Mingolla, 1985; Fitzpatrick, 2000; Z. Li, 1998, 2000; Craft, Schütze, Niebur, & von der Heydt, 2007).

The various sources of suppression are expected to have different temporal properties. There is evidence that neuron-to-neuron signals propagate much faster along feedforward and feedback connections than along intracortical connections in V1 (Bringuier, Chavane, Glaeser, & Frégnac, 1999; Grinvald, Lieke, Frostig, & Hildesheim, 1994). This difference in temporal properties is expected to be larger for surround suppression because the temporal delay of the signal caused by the intracortical connection depends on the cortical distance, and the cortical distance depends on the distance between the corresponding points on the retina. The latency of the surround suppression becomes longer as the surround stimuli become further from the neurons’ classical RFs (Bair et al., 2003). It has been shown that the feedback signal of the surround suppression from area MT is delayed less than 10 ms from the initial responses of the neurons (anesthetized macaque monkey: Hupé, James, Girard, Lomber, et al., 2001). The feedback signal from V2 and V3 is delayed by about 20 ms (alert macaque monkey: Nassi et al., 2013). The shorter latency of the surround suppression from MT can be attributed to a direct connection between the LGN and MT (Sincich, Park, Wohlgemuth, & Horton, 2004; Stepniewska, Qi, & Kaas, 1999).

In addition to the differences in their latencies, surround-suppression effects can be differentiated by their properties in different domains. Bair, Cavanaugh, and Movshon (2003) reported a negative correlation between strength and latency of the surround suppression across the neurons in their sample. The delay of the surround suppression from the initial response of the neurons could be as long as 30 ms for weak suppression, whereas it could be almost 0 ms (as early as the initial response) for strong suppression. Knierim and Van Essen (1992) tested the temporal properties of surround suppression while they controlled the orientation of the surround stimuli. Their results suggested that the surround suppression with a short delay (about 7 ms) was isotropic, but with a long delay (20 ms) it was selective to the orientation of the surround stimuli. Knierim and Van Essen argued that this isotropic surround suppression should be attributed to the neurons or to their processes in the LGN (or in the retina) because many of the subcortical neurons are insensitive to orientations of stimuli in their classical RFs or in their surround regions (see Bonin et al., 2005; Zaltsman, Heimel, & Van Hooser, 2015, for reviews).

Recall that the response of some neurons is facilitated by their surround stimuli (Fitzpatrick, 2000; C.-Y. Li & Li, 1994; Maffei & Fiorentini, 1976; Sillito et al., 1995; Cavanaugh et al., 2002b; Polat et al., 1998; H. E. Jones et al., 2001; Nelson & Frost, 1985). This facilitatory effect comes with a longer delay (40 ms) from the initial response than the facilitation produced by the suppressive effect (20 ms) (Hupé, James, Girard, & Bullier, 2001). It is worth pointing out that the delay of the suppressive effect (20 ms) observed by Hupé et al. (2001, anesthetized macaque monkey) is close to the delay observed in Nassi et al. (2013, alert macaque monkey). This suppressive effect was not affected by inactivating V2 (Hupé, James, Girard, & Bullier, 2001) but it was weakened (or eliminated) by inactivating V2 and V3 (Nassi et al., 2013).

Some effects of the surround stimuli can be observed only by studying the temporal properties of the neuron’s responses. Briggs and Usrey (2011) found that the initial response of the neurons becomes faster or slower by about 5 ms with the surround stimuli depending on the particular neuron. Also, the surround stimuli, themselves, can cause an excitatory signal and evoke firing responses in many neurons without any stimulation within the classical RFs with a very long latency (≥ 100 ms, W. Li et al., 2001; Rossi, Desimone, & Ungerleider, 2001). The latency of this excitatory effect from the surround stimuli is not affected by the distance of the surround stimuli from the RFs (Rossi et al., 2001). This constant latency of the excitatory effect suggests that this effect cannot be attributed to intracortical connections because the temporal delay of a signal mediated by these connections depends on the distance on the retina.

In sum, the surround suppression appears to be a combination of effects from different sources with different temporal and spatial properties. Cross-orientation suppression probably is similarly heterogeneous. It will be difficult to account for such complexity with a simple functional model.

### 4.4 Directions for future work

In this study, we considered only simple synthetic visual stimuli within a limited spatial context in a static (or steady-state) experimental condition. We did not assess the performance of the model under more ecologically-valid stimulation. There is a growing number of physiological studies that recorded responses from V1 neurons to photographs and to videos of real visual scenes (e.g., Touryan, Felsen, & Dan, 2005; Smyth et al., 2003), but Rust and Movshon (2005) warned about the interpretative difficulties inherent in the use of such complex stimuli. Furthermore, our review focused on relatively simple (first-order) properties of the stimuli, despite the fact that the responses of many real neurons can be affected by higher-order visual information including figure-ground organization (Hupé et al., 1998; Zhou, Friedman, & von der Heydt, 2000; Zhang & von der Heydt, 2010; Rossi et al., 2001), 3D context (Murray, Kersten, Olshausen, Schrater, & Woods, 2002; Murray, Boyaci, & Kersten, 2006), perceptual filling-in (Fiorani Jr., Rosa, Gattass, & Rocha-Miranda, 1992; Komatsu, 2006), and visual illusions (Ramsden, Hung, & Roe, 2001; see also Jancke, Chavane, Naaman, & Grinvald, 2004 for a brain imaging study). This means that no model can fully replicate the response of a neuron to natural stimuli unless such higher order information is taken into account. On the other hand, extracting such higher-order information from a retinal image is an open research problem. One practical way to circumvent this problem would be to prepare the higher-order information in advance and make it available to the model as its inputs from a top-down process.

The divisive normalization model (DNM) has also been used to fit physiological results of populations of neurons in V1 (Busse et al., 2009; Goris et al., 2009). Note that modeling populations is different from modeling single cells, for the following reasons. First, a population may be able to process visual information better than individual neurons by forming a *population* code in which different neurons specialize in encoding different aspects of the input signal (deCharms & Zador, 2000; Pouget et al., 2003). For example, the neuronal population in V1 can systematically represent the second-order visual information while only a subset of the individual cells in V1 respond selectively to second-order stimuli (e.g., An et al., 2014). This complicates the derivation of model predictions about the population response because a given physiological phenomenon may have two possible explanations: one in terms of a population code and another in terms of the properties of individual neurons (e.g., Footnote 8 above). Another difficulty arises from the selective sampling of neurons in physiological experiments. It has been pointed out that recordings from some neurons are often excluded from the data set, producing a selection bias that can affect the experimental results (Olshausen & Field, 2005). So, the physiological results of the population can change depending on which neurons are included/excluded in the population. This can add extra parameters to the model of the population. On the other hand, the selection bias is less critical for single-cell recording because the model aims to account for the response properties of individual neurons. These differences suggest that different approaches may be required to model populations of neurons as opposed to modeling individual neurons. One fruitful area for future research is to explore the degree to which the divisive-normalization equation is applicable to both cases.

In conclusion, the divisive normalization model provides a useful functional characterization of the responses of simple and complex cells in primary visual cortex. It deserves to be designated as “the standard” model for many present purposes. We hope that the standard formulation proposed here, the standard parameter set, and the accompanying software implementation will facilitate future research based on this influential and successful model. Of course, the DNM will be supplanted and/or subsumed by future, more advanced models, just as it subsumed the “standard” linear model of the 1980s (Rust & Movshon, 2005). It seems particularly desirable to augment the DNM with mechanisms to account for the temporal dynamics of neuronal responses (e.g., Heeger, 1993; Brosch & Neumann, 2014). Accounting for this temporal dimension, however, poses interesting challenges. It is hard to see, for example, how a simple functional model can account for the temporal properties of surround suppression (and facilitation), which are attributable to several different sources as discussed above. Such an augmented model would need additional components and new adjustable parameters. Ideally, the components and parameters should have physiological interpretations that relate to physiological and/or anatomical data. Developing the DNM along those lines will bring it closer to a structural model.

## 5 Acknowledgement

The authors thank Kayla Higginbotham for her help in digitizing data from published figures.

## Appendix A A 2D Gabor filter and its tuning functions

Consider orientation and spatial-frequency tuning functions of a 2D Gabor filter (Equations 2 and 5) tuned to a spatial-frequency *F* (cycles/deg, cpd) and an orientation Θ (°). The orientation tuning function of the 2D Gabor filter is cyclic and becomes maximal at Θ and Θ + 180°. The shapes of those two peaks are the same and the orientation bandwidth *h_θ_* (°) is defined as a width of each peak at its half-height. The orientation tuning function for a grating with spatial-frequency *F* is equivalent with a circular cross section of the 2D Fourier transform of the filter with a radius *F.* Note that a Fourier spectrum energy distribution of the 2D Gabor filter is a 2D Gauss distribution (see Figures 2.10 and 2.11 in Graham, 1989). Then, a relation between the orientation bandwidth and the bandwidth 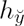 of the Gabor filter along *θ* can be written as follows (see Tables 2.2 and 2.4 in Graham, 1989):

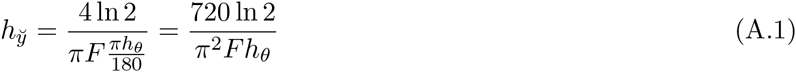

The spatial-frequency tuning function is unimodal and becomes maximal at *F* (cpd). Its bandwidth *h*_*f*_ (oct) is defined as a distance between two frequencies *F*_*low*_ and *F*_*high*_ (cpd) at the half-height of the function where *F*_*low*_ < *F*_*high*_. Then, the relation between *F*, *F*_*low*_, *F*_*high*_, and *h*_*f*_ is:

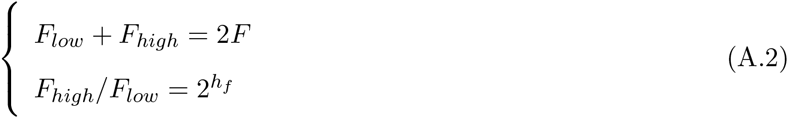

From Equation A.2, the bandwidth *F*_*high*_ − *F*_*low*_ in cpd can be derived as follows:

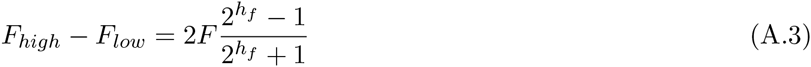

Recall that the Fourier spectrum energy distribution of the 2D Gabor filter is the 2D Gauss function. The spatial-frequency tuning function of the filter for a grating with an orientation *θ* is equivalent with a straight cross section of the Gauss function along an orientation *θ* + 90°. Then, a relation between the frequency bandwidth *F*_*high*_ − *F*_*low*_ and the bandwidth 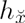 of the Gabor filter for its tuned orientation Θ can be written as follows (see Tables 2.2 and 2.4 in Graham, 1989):

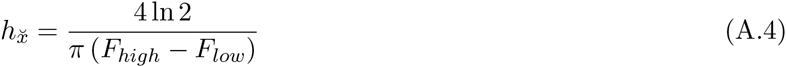

From Equations A.3 and A.4, a relation between 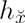 and *h*_*f*_ can be written as follows:

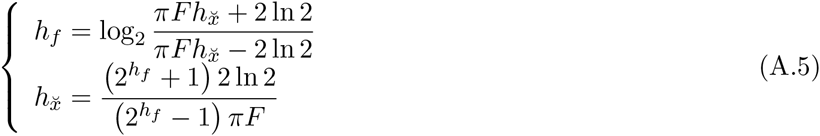

## Appendix B The Contrast Response Function of the hyperbolic-ratio model along the linear contrast axis

We analyze the contrast response function (CF) of the hyperbolic-ratio model (Equation 10) on a linear contrast axis. The CF is a function of the luminance contrast *c* of a grating stimulus. Note that the orientation and spatial-frequency of the grating are consistent with the tuning of the model and the size of the grating is large enough to fill the entire receptive field of the model.

The model becomes most sensitive to a change in luminance contrast *c* where the slope of the CF becomes maximal (see Itti et al., 2000; Wilson, 1980, see also Section 3.3.2). The maximal CF slope is:

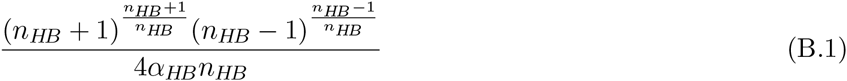

This maximum occurs for contrast

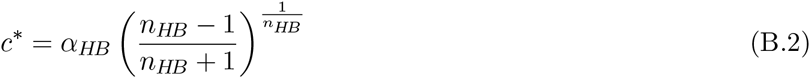

if *n*_*HB*_ > 1 and for c* = 0 otherwise (Figure 24). If *n*_*HB*_ > 1, the CF is convex downward around *c* = 0. This trend of the model CF can account for the shapes of CFs of real neurons around *c* = 0 (Albrecht & Hamilton, 1982; Albrecht et al., 2003).

Some other studies (e.g., Tadmor & Tolhurst, 1989; Dean, 1981; Heeger, 1992a) represent the shapes of the CFs of real neurons around *c* = 0 using a piecewise-linear model (Section 2.1) with a threshold *β_IB_*:

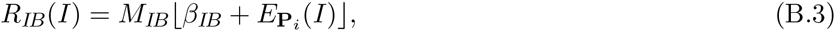

where *β_IB_* < 0. Note that CFs of real neurons often start to saturate at high contrasts (see Figure 4 for CFs of real neurons plotted on the log contrast axis). This trend can also be captured by the hyperbolic-ratio model (Figure 24, see also Figure 4 for the CFs of the hyperbolic-ratio model on the log contrast axis) but cannot be captured by the linear model.

**Figure 24:**
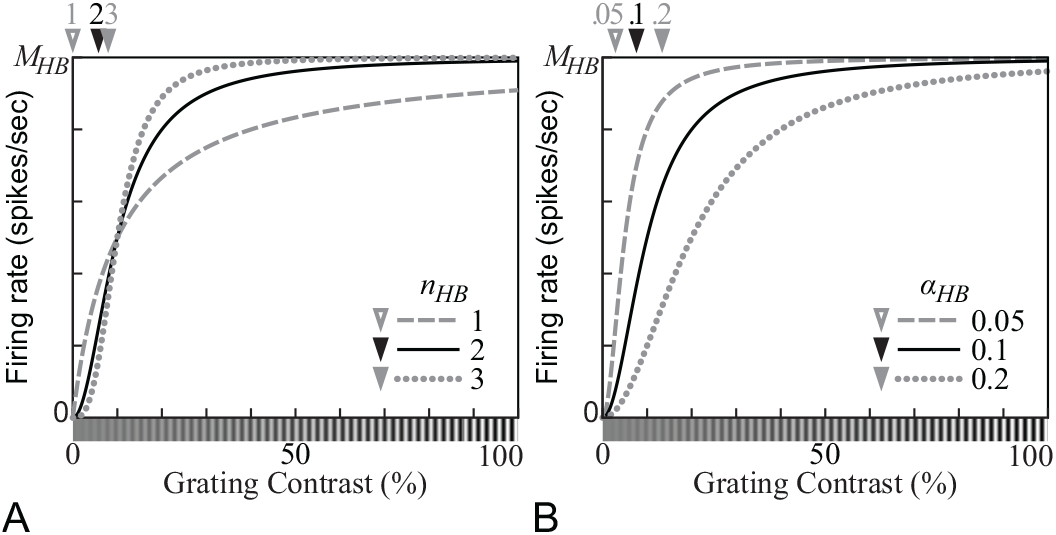
Contrast response functions (CFs) produced by the hyperbolic-ratio model (Equation 10) on the linear-contrast axis. Positions of the maximal slopes of the CFs are indicated by triangles. (A) The CFs with three different values of the exponent parameter *n_HB_.* (B) The CFs with three different values of the *semisaturation contrast* parameter α_*HB*_. A stimulus with contrast *α*_*HB*_ elicits one-half of the saturation level *M*_*HB*_. (*α*_*HB*_ = 0.1 for panel A; *n*_*HB*_ = 2 for panel B.)

## Appendix C The shape of the contrast response function of the divisive normalization model

The shape of the contrast response function (CF) of the divisive normalization model (DNM) is mathematically analyzed in this appendix. Consider the DNM (Equation 13) and a grating *g*(*c*) with contrast *c*, the model’s tuned orientation and frequency (and phase for a simple cell), and spatial extent large enough to fill both the entire receptive and suppressive fields of the model. Then, the CF of the DNM (Equation 13) for *g*(*c*) is:

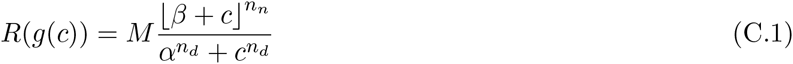

(Note that this equation is equivalent to Equation 15 in Section 2.3.) The first derivative of Equation C.1 with respect to *c* is:

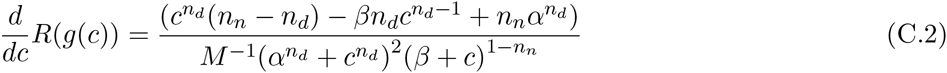

for 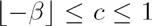. Note that *dR*(*g*(*c*))/*dc* >0 at *c* = 0 if *β* > 0 and at 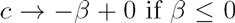. This means the CF is increasing at low contrasts. The numerator of Equation C.2 is further differentiated:

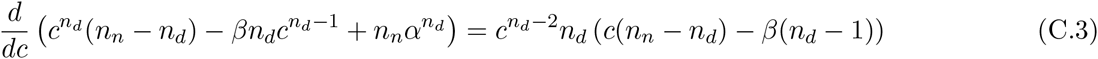

The derivative of the numerator becomes 0 at *c* = 0 and *c = c_ex_* where

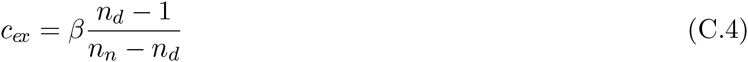

Hence, the numerator has two local extrema: 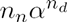 at *c* = 0 and

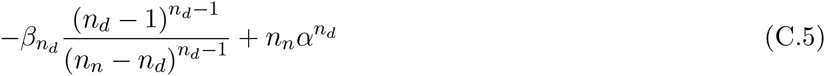

at *c* = *c*_*ex*_. Besides, the denominator of Equation C.2 is always positive between 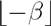 and 1, exclusive. Then, the possible shapes of the contrast sensitivity curves between 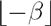 and 1 can be categorized into the following three types.

(1) The sensitivity curve is unimodal and convex-upward if 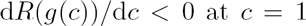. This condition is satisfied if *β* is large enough:

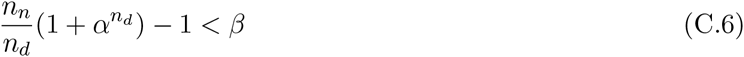

The CF is decreasing at high contrasts and must have a local maximum between 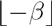 and 1, exclusive. Namely, the curve shows the super-saturation effect. This is the sufficient condition for the super-saturation effect for *g*(*c*).

(2) The CF is with both local maximum and minimum if d*R*(*g*(*c*))/d*c* > 0 at *c* = 1, d *R*(*g*(*c*))/d*c* < 0 at *c* = *c*_*ex*_ (see Equation C.5), and 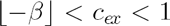. The local maximum appear between 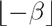 and 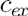 and the local minimum appears between *c*_*ex*_ and 1. Note that it may not be practically well-conditioned to judge those conditions based on physiological results. Many physiological studies have shown that the DNM can fit data well under an assumption that *n*_*n*_ = *n_d_.* Then, *n*_*n*_ − *n*_*d*_ can be too small compared with *n*_*d*_ − 1 to have stable estimates of *c*_*ex*_ and d*R*(*g*(*c*))/d*c* at *c* = *c*_*ex*_ (Equation C.5).

(3) The CF is monotonically increasing otherwise.

## Appendix D Contrast response function for a non-preferred stimulus

Consider the divisive normalization model (DNM, Equation 13) and a grating *g′*(*c*) with contrast *c* but with non-preferred orientation, spatial-frequency, or size. The stimulus drive (Equations 5 and 7) and the suppressive drive are still proportional to *c* between 0 ≤ *c* ≤ 1 because *g′*(*c*) = *cg′*(*1*). Then, the contrast response function (CF) of the DNM (Equation 13, see also Equation C.1 above) for *g′*(*c*) is:

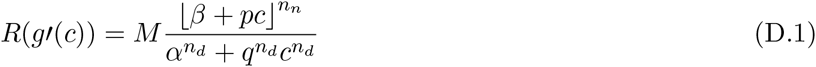

where *p* and *q* are constants. Equation D.1 can be modified as follows:

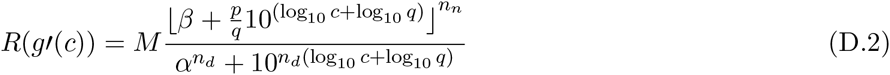

Equation D.2 shows that a relation between plots of *R*(*g*′(*c*)) and *R*(*g*(*c*)) (Equation C.1) can be represented as a subset of affine transformation with *c* in a logarithmic scale and *β* sufficiently smaller than *pc.* The subset is translation − log_10_ *q* along the contrast axis and scaling (*p/q*)*^nn^* along the response axis.

## Appendix E Computational implementation of the divisive normalization model

Computational implementation of the divisive normalization model (DNM, Equation 13) allows the model to be applied to any visual stimulus used in physiological and also psychophysical studies. Its computational efficiency and practical usage should be also taken into consideration.

The suppressive drive of the DN model (Equation 16) is computationally demanding. It is represented as a weighted sum of 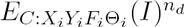 (Equation 7) in the orientation, spatial frequency, and 2D retinal space domains, where 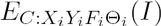 is a square root of a sum of 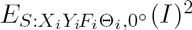 and 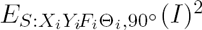 (Equation 5). A part of the weighted sum of the suppressive drive in the 2D retinal space domain can be computed using 2D cross-correlation of a retinal image *I*(*x, y*) (Equation 1) with two 2D Gabor filters 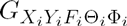 in Equation 2:

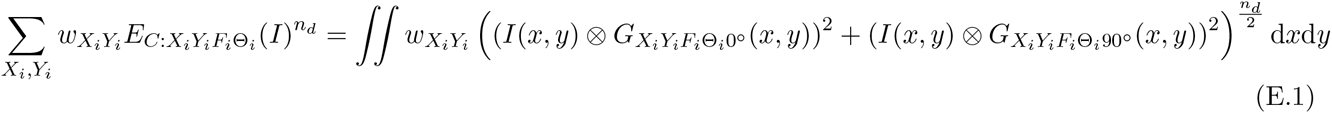

where an operator ⊗ represents a cross-correlation^9^. This is equivalent with computing 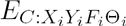 at every pixel of the input image. Then, the number of the 2D spatial sampling *N*_*X*_ × *N*_*Y*_ becomes equivalent with that of pixels in the filtered image.

Note that the computational cost of the cross-correlation is an order of 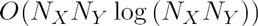 and it has to be processed for 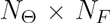 times to compute the suppressive drive. Then, the total cost becomes 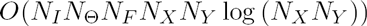 if the number of the input images is *N_I_.* This process can be optimized using the convolution theory. Namely, the image filters in Equation 2 are applied to the input image in a 2D Fourier space:

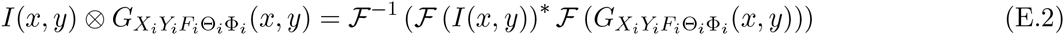

where operators 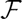 and 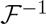 represent a Fourier transform and its inverse and * represents a conjugation. The Fourier transforms of the image filters 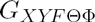 can be computed in advance and can be used for processing every image. Then, its total cost is 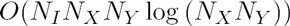.

Note that Equations E.1 and E.2 become approximations because the sizes of *I* and 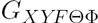 are practically both finite. The approximation becomes better if *I* and 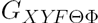 are surrounded by sufficiently large regions with uniform gray. The size of *I* with the surrounding region should be the same as that of 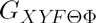 and be 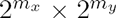 where *m*_*x*_ and *m*_*y*_ are positive integers. It allows the model to process the image efficiently using the Fast Fourier Transform algorithm in Equation E.2.

The process is further optimized so that the model can compute the suppressive drives of multiple “virtual neurons” to the single input image at once. The results of the image filtering process in Equation E.2 can be shared among the virtual neurons. It generates a set of the 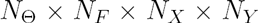 different channels. The suppressive drives of the multiple virtual neurons can be computed from the same set of the channels with different distributions of the weights 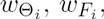 and 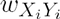.

In the process of computing the suppressive drives, 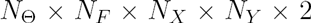 different 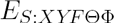 and 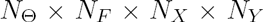 different 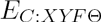 are generated. They can be also used as the stimulus drives of the virtual neurons. Note that we practically need a limited number *N_C_,* instead of *N*_*X*_ × *N_Y_,* of the 2D spatial positions of the virtual neurons, especially when the resolution of the input image is high. Also, not all *N*_*F*_ spatial-frequencies are appropriate for the stimulus drives because of the edge effect. The domain of spatial-frequency ranges between ±∞ but 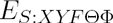 and 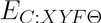 can be sampled within its limited range. Consider virtual neurons whose tuned frequencies are on the upper (or lower) bound of the sampling range in the domain. They cannot be suppressed by the channels whose tuned frequencies are higher (or lower) than the tuned frequency of the neurons. The edge effect can be lightened by removing some neurons whose tuned frequencies are close to the bounds of the sampling range. Hence, there are 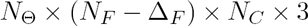 virtual neurons generated in total: 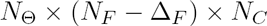 complex cells and 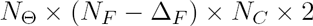 simple cells (see Equation 7), where *∆_F_* is the number of the removed neurons for the edge effect.

## Footnotes

1 A time-varying stimulus such as a drifting grating also has temporal frequency. As the temporal properties are beyond our scope, however, the term “frequency” refers to spatial frequency throughout this article unless explicitly stated otherwise.

2 Published physiological data were captured from their respective figures using PlotDigitizer (http://plotdigitizer.sourceforge.net) and replotted here in a unified format.

3 *I* and *G*_*XYFΘΦ*_ are treated as (long) vectors in the computation of the dot product.

4 The abbreviation CRF should be to be avoided because it tends to be misread as a “classical receptive field.”

5 The function *R*(*c*) = *c^n^*/(α^n^ + c^n^) is sometimes called the *Naka-Rushton* function. In biochemistry, it is called the *Michaelis-Menten* function and is the centerpiece of a standard model of enzyme kinetics.

6 The corresponding term in Grossberg (1973, 1988; Ellias & Grossberg, 1975) is *mass action.*

7 This assumption ignores non-separable effects such as end stopping and contour integration, which require coordination among components with aligned orientations at neighboring locations (Kapadia, Ito, Gilbert, & Westheimer, 1995; Z. Li 1998).

8 This prediction seems to be consistent with psychophysical results showing that human performance in contrast detection is better with a square grating than with a sinusoidal grating (Campbell & Robson, 1968). Note, however, that the square grating is expected to stimulates a wider variety of V1 neurons because a square grating is composed of multiple sinusoidal gratings, so these psychophysical results can be explained by properties of either single V1 neurons or by a population of V1 neurons.

9 If the resolution of the image I is high enough, the following approximation also works: 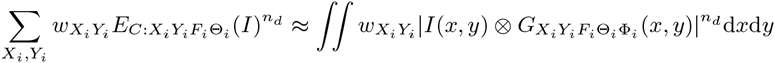 where Φ_*i*_ is arbitrary.

